# Hydroxychloroquine-functionalized Ionizable Lipids Mitigate Inflammatory Responses in mRNA Therapeutics

**DOI:** 10.1101/2025.03.02.641033

**Authors:** Kepan Chen, Xuechun Li, Shuo Feng, Ye Li, Ting Jiang, Yan Liu, Na Guo, Xiaoling Zeng, Hongwei Yao, Min Qiu, Jing Lu, Jinzhong Lin

## Abstract

Lipid nanoparticle (LNP)-based mRNA therapeutics, highlighted by the success of SARS-CoV-2 vaccines, face challenges due to inflammation caused by ionizable lipids. These ionizable lipids can activate the immune system, particularly when co-delivered with nucleic acids, leading to undesirable inflammatory responses. We introduce a novel class of anti-inflammatory ionizable lipids functionalized with hydroxychloroquine (HCQ), which suppresses both lipid-induced and nucleic acid-induced immune activation. These HCQ-functionalized LNPs (HL LNPs) exhibit reduced proinflammatory responses while maintaining efficient mRNA delivery. Structural and physicochemical analyses revealed that HCQ-functionalization results in a distinct particle structure with significantly improved stability. The efficacy of HL LNPs was demonstrated across various therapeutic contexts, including a prophylactic vaccination model against varicella-zoster virus (VZV) and CRISPR-Cas9 gene editing targeting PCSK9. Notably, HL LNPs showed robust mRNA expression after repeated administration, addressing concerns of inflammation and ensuring sustained therapeutic effects. These findings highlight the potential of HCQ-functionalized LNPs in expanding the safe use of mRNA therapeutics, particularly for applications requiring repeated dosing and in scenarios where inflammation-induced side effects must be minimized.

## Introduction

The success of mRNA vaccines, such as Moderna’s mRNA-1273^1^ and Pfizer/BioNTech’s BNT162b2^2^ against SARS-CoV-2, has accelerated the development of mRNA-based medicines^3^. Recent approvals, including Moderna’s mRNA-1345 against respiratory syncytial virus (RSV), underscore the potential of mRNA technologies in combating diverse diseases. This progress is largely due to the inherent advantages of mRNA vaccines, including potent immunogenicity, short development timelines, and scalable production^4^. However, naked mRNA, as a negatively charged macromolecule, cannot penetrate cell membranes and is prone to degradation by nucleases, necessitating the use of specialized delivery systems, such as lipid nanoparticles (LNPs), lipoplexes, virus-like particles (VLPs), dendritic cells, and exosomes^5^. Among these, LNPs have emerged as the most effective and widely used delivery system^6,7^, including in the three currently approved mRNA vaccines.

LNPs typically consist of four main components: ionizable lipids, neutral phospholipids, cholesterol, and poly(ethylene glycol) (PEG)-modified lipids^7^. Ionizable lipids, which can be protonated in an acidic environment, are crucial for mRNA packaging, protection, intracellular delivery, and endosomal escape of mRNA^8^. For example, the ionizable lipid in the mRNA-1273 vaccine is SM-102, and that in the BNT162b2 vaccine is ALC-0315. These lipids also have an important impact on the in vivo fate of LNPs, including the distribution and expression of mRNA within different organs or cells^9,10^. LNP outperforms other vehicles mainly because of their stable and controllable quality, high loading efficiency, and tunable targeting property^6,7^.

Despite these advantages, the mRNA-LNP partners face challenges, particularly related to their inflammatory side effects. Even though chemical modifications have reduced the immunogenicity of mRNA, ionizable lipids can still induce innate immune responses^11–13^. Both ionizable lipids and exogenous mRNA can activate pattern recognition receptors (PPRs) such as Toll-like receptors (TLRs)^14,15^, triggering the release of proinflammatory cytokines like tumor necrosis factor-α (TNF-α), interferon-γ (IFN-γ), interleukin-6 (IL-6), and interleukin-1 (IL-1)^16–18^. These cytokines may enhance the efficacy of therapeutic tumor vaccines^19^, but are undesirable in other applications, such as prophylactic vaccines and protein replacement therapies, where excessive immune stimulations provoke adverse inflammation-related effects. Moreover, inflammation caused by ionizable lipids is particularly concerning in scenarios involving repeated administration of mRNA-LNPs, where repeated immune activation can further affect mRNA expression and cause cumulative damage^20–22^. As ionizable lipids are essential for LNP formation, there is an urgent need to develop novel ionizable lipids that do not provoke inflammatory reactions, thereby improving the safety of LNPs and expanding the applications of mRNA-based medicines^20,23^. Additionally, ionizable lipids that can suppress inflammatory responses would be particularly beneficial for highly immunogenic nucleic acid drugs, such as self-amplifying RNA (saRNA)^24^.

Given the expanding application fields of LNP-based mRNA medicines, there is growing interest in developing the next generation of multifunctional ionizable lipids or LNPs^25–28^. While current research has focused on designing organ-targeted ionizable lipids (e.g., AID-lipids^29^, nAcx-Cm lipids^30^), there have also been efforts to functionalize ionizable lipids by incorporating small-molecule drugs into their structures. For example, TLR7/8 agonists, such as R848, have been conjugated with lipid tails to form functionalized lipids like RAL2^31^ and C12-TLRa^32^, which enhance the innate immunity of mRNA vaccines against melanoma and SARS-CoV-2, respectively. Similarly, non-nucleotide agonists of the stimulator of interferon genes (STING) have been integrated into ionizable lipids to enhance protective immunity by activating the STING pathway^33^. These studies, which aim to enhance the immunogenicity of mRNA vaccines, also provide a new approach for the functionalization of ionizable lipids.

Herein, we aimed to mitigate the inflammatory side effects of LNP-based mRNA medicines by developing a class of anti-inflammatory ionizable lipids. Hydroxychloroquine (HCQ), originally developed as an antimalarial drug, has gained widespread use as an anti-inflammatory agent in autoimmune diseases such as rheumatoid arthritis (RA) and systemic lupus erythematosus (SLE) ^34^. HCQ exerts its effects through multiple mechanisms, including the inhibition of autophagy, endosomal TLR activation, and the cGAS-STING signaling pathway^34–36^. These pathways are crucial for the production of proinflammatory cytokines, and their inhibition by HCQ leads to a reduction in cytokine levels, including TNF-α, IFN-γ, and IL-6^34^. Given HCQ’s broad anti-inflammatory properties, we propose the functionalization of ionizable lipids with HCQ to mitigate the inflammatory side effects associated with LNP-based mRNA therapeutics. Here, we introduce a new class of HCQ-derived ionizable lipids (HLs) designed to mitigate the inflammatory side effects of LNPs while maintaining efficient mRNA delivery (Fig. 1a). We demonstrate that HL-based LNPs retain the mRNA expression capacity of conventional LNPs, but exhibit a distinct ability to suppress cytokine production. Notably, they exhibited completely different physicochemical and structural properties from the LNP formed of conventional ionizable lipids. Through multiple in vivo models, including repeated intravenous administration, a prophylactic mRNA vaccine against varicella zoster virus (VZV), and CRISPR-Cas9 mRNA-based gene editing, we show that HL LNPs significantly reduce the induction of proinflammatory cytokines compared to conventional SM-102 LNP. These results highlight the potential of HCQ-functionalized lipids in expanding the safe use of LNP-based mRNA medicines.

**Fig. 1.**
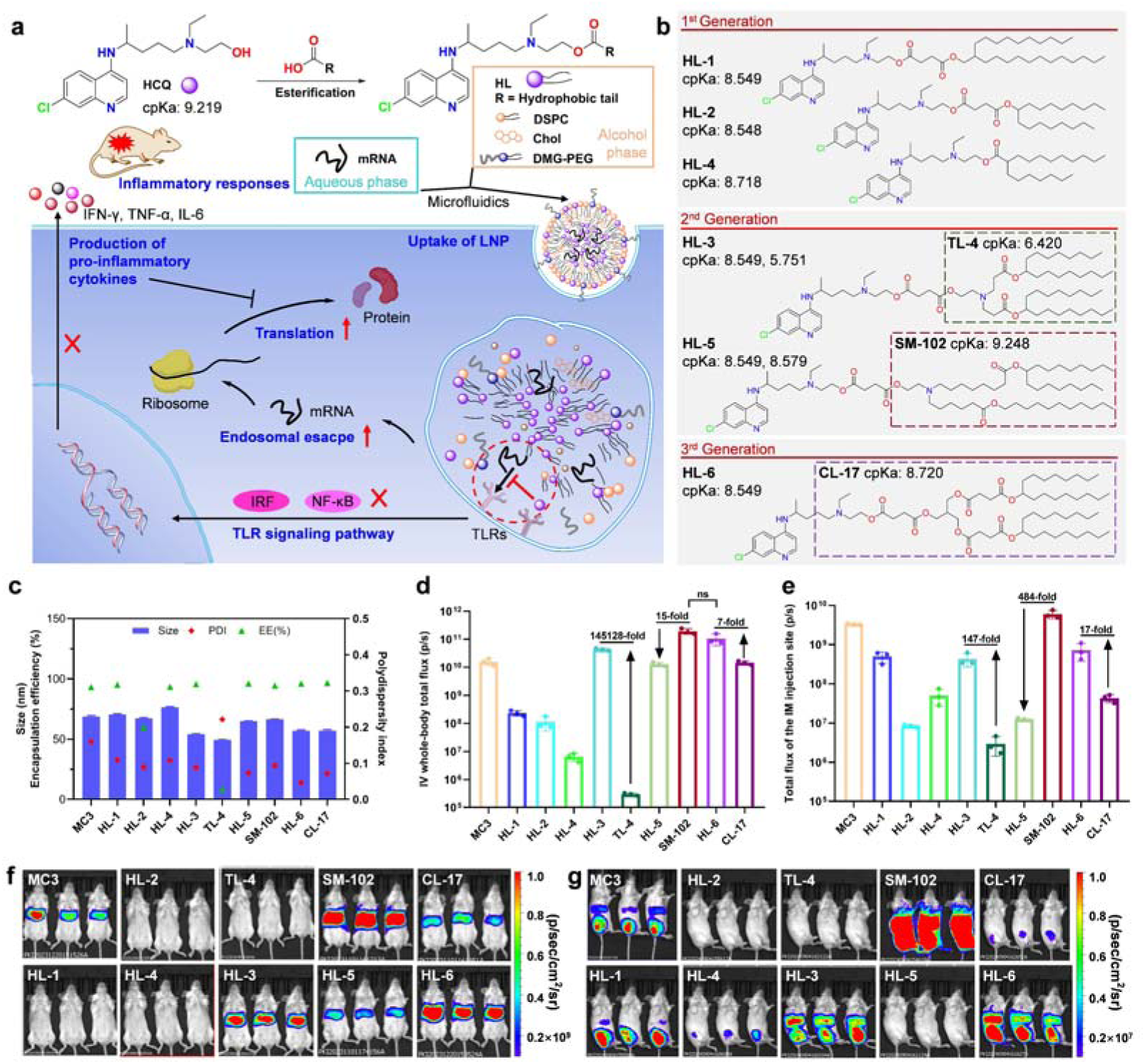
Anti-inflammatory design, characterization and in vivo delivery efficiency of HL LNPs. **a**, Procedures for the synthesis of HLs, preparation of LNPs, and intracellular inhibition of cytokine production. HL compounds were synthesized via the esterification reaction between HCQ and hydrophobic tails containing carboxyl groups. Subsequently, a mixed lipid alcohol solution comprising HL, DSPC, cholesterol and DMG-PEG, along with an acidic mRNA solution, was squeezed through a microfluidic chip to formulate LNP preparations. Following the uptake of LNPs by cells, HLs facilitated the escape of mRNA from endosomes and inhibited the TLR signaling pathway, ultimately leading to the reduction in the release of pro-inflammatory cytokines and the enhancement of mRNA translation. **b**, Chemical structures and cpKa values of HLs and their corresponding tail-controlled ionizable lipids. **c**, Particle size, PDI and encapsulation efficiency of LNP loaded with FLuc mRNA. **d**, Intensity of whole-body bioluminescence observed 4 h following intravenous administration (IV). **e**, Intensity of bioluminescence at the injection site 4 h following intramuscular administration (IM). **f**, Bioluminescence images of all groups obtained 4 h following intravenous administration. **g**, Bioluminescence images of all groups obtained 4 h following intramuscular administration. The data are presented as mean ± SD (n = 3).

## Results

### Rational design and synthesis of hydroxychloroquine-functionalized lipids

HCQ, a 4-aminoquinoline antimalarial drug, processes an ionizable tertiary amine in its side chain. This feature, along with a hydroxyl group on its side chain, distinguishes it from chloroquine and gives it a calculated pKa (cpKa) of 9.219 (Fig. 1a). The intrinsic nitrogen in HCQ’s structure makes it a highly suitable candidate for lipid derivation, allowing it to function as an ionizable head group or be conjugated to other ionizable lipids.

Therefore, in our designs, we utilized this flexibility in two ways: in some HL variants, the intrinsic nitrogen of HCQ servers as the ionizable head, while in others, HCQ is conjugated to existing ionizable lipids that contained their own ionizable nitrogen, allowing for multi-functional lipid structures (Fig. 1a). To synthesize the HCQ-derived ionizable lipids, we esterified hydrophobic tails containing carboxyl groups to the hydroxyl group of HCQ (Fig. 1a). Based on the structure of the tail, we developed three generations of HLs (Fig. 1b). For the first generation, the hydrophobic tail consisted of a simple long-chain aliphatic acid (HL-4) or a succinate derivative formed by reacting a long-chain aliphatic alcohol (HL-1, HL-2). These designs prioritize simplicity in the tail structure while using HCQ as the ionizable head group. In the second generation, HCQ was conjugated to other ionizable lipids: TL-4 for HL-3 and SM-102 for HL-5, each containing their own ionizable nitrogen in the tail. In the third generation of HLs, the design similar to the first generation except that the tails are more complicated, with multiple branches as in HL-6, which was synthesized by linking heptadecan-9-ol with tri-hydroxymethyl-methane and succinic anhydride. Unlike HL-3 and HL-5, the tail of HL-6 does not contain a tertiary amino group.

To isolate the contribution of HCQ, we also synthesized tail-controlled lipids without HCQ or the aminoquinoline group, namely TL-4, SM-102, and CL-17 for comparison with HL-3, HL-5, and HL-6, respectively (Fig. 1b).

In summary, the HCQ-derived lipids (HLs) leverage the intrinsic ionizable nitrogen in HCQ’s side chain, either as the primary ionizable head group or in conjunction with another ionizable group in the lipid tail. All these lipid compounds were synthesized according to the designed synthetic routes (Supplementary Fig. 1-6), purified by silica gel column chromatography, and their structures were confirmed by NMR and MS (Supplementary Fig. 7-15).

### Preparation, characterization and in vivo evaluation of HL LNPs

To validate the functionality of HCQ-derived ionizable lipids, we prepared HL lipid nanoparticles (HL LNPs) by mixing HLs with neutral phospholipids, cholesterol, and PEG-modified lipids at a mol/mol ratio of 50:10:38.5:1.5. mRNA was encapsulated in LNPs, and the nitrogen-to-phosphate (N:P) ratio between the ionizable lipid and mRNA was maintained at 6:1, with one ionizable nitrogen considered for all HLs. mRNA-loaded LNPs were formulated using a microfluidic mixing method. Tail-controlled lipids (TL-4, SM-102, CL-17) were formulated into LNPs using the same formulation and procedure. The approved ionizable lipid DLin-MC3-DMA (MC3) was used as a positive control.

Firefly Luciferase (FLuc) mRNA was encapsulated into different LNPs, and the particle size, polydispersity index (PDI) and encapsulation efficiency were characterized (Fig. 1c). All HL LNPs exhibited uniform particle sizes (54-76 nm), with PDIs below 0.2, indicating good colloidal stability. The encapsulation efficiency of HL LNPs was generally over 90%, except for HL-2, which had an encapsulation efficiency of around 60%. This result of HL-2 with a shorter tail suggests that tail length plays an important role in encapsulating mRNA. A comparison between HL-3 and its tail-controlled lipid TL-4 further revealed HCQ’s role in mRNA encapsulation. TL-4 failed to encapsulate mRNA efficiently, with efficiency below 5%, likely due to its low cpKa of 6.4, which is too low to be protonated and capture mRNA efficiently. However, after conjugation with HCQ to form HL-3, the encapsulation efficiency significantly improved to above 95%, suggesting that HCQ contributes to the encapsulation process, most likely through its intrinsic ionizable nitrogen on the alkaline side chain. Consequently, HL-6 also performed well. Further characterization revealed that the aminoquinoline ring of HCQ also contributes to mRNA encapsulation and LNP stability through extensive interactions with mRNA, which will be discussed later.

Next, we evaluated the in vivo mRNA delivery efficiency. Each LNP loaded with FLuc mRNA was injected intravenously into BALB/c mice at a dose of 5 μg mRNA per mouse, and the luciferase expression was monitored at 4 h, 24 h and 48 hours post-injection using bioluminescence imaging (Fig. 1d, f and Supplementary Fig. 16). The bioluminescence-time curve was plotted, and the area under the curve over 4 h to 48 h (AUC_0-48h_) was calculated to quantify total expression over time. In addition, the LNPs were tested for in vivo efficiency via the intramuscular administration route (Fig. 1e, g and Supplementary Fig. 17).

The results show that the first-generation HL LNPs (HL-1, HL-2, and HL-4) failed to deliver mRNA efficiently via intravenous administration compared to MC3. However, via the intramuscular route, HL-1 and HL-4 LNPs were able to deliver mRNA, with HL-1 being more efficient than HL-4 and reaching the same level of performance as MC3. The second-generation HL-3 LNP outperformed MC3 in both intravenous and intramuscular administration, demonstrating the importance of tail structure in delivery. This becomes more evident in the third-generation HL-6 LNP, which was significantly more efficient than MC3 LNP and similar to SM-102 LNP via intravenous administration, though slightly less efficient than SM-102 via intramuscular administration. In comparison, the tail-controlled lipid CL-17 performed poorly, suggesting that the aminoquinoline structure plays a crucial role in delivery.

These results provide two key insights. First, direct lipid derivatization of HCQ can achieve both efficient mRNA encapsulation and delivery. Second, this activity relies on proper selection of the lipid tail. Based on these findings, we selected HL-3 and HL-6 LNPs for further investigation.

### Physicochemical stability and structural characterization of HL LNPs

Due to the inherent instability of mRNA, LNPs typically need to be stored at freezing temperatures to prevent mRNA degradation. This requires LNP formulations to withstand the challenges of the freeze-thaw cycles, which can affect their pharmaceutical properties. To assess the stability of HL LNPs, we subjected them to five freeze-thaw cycles at either −80 or −20, followed by thawing at 4. We then analyzed particle size, PDI, encapsulation efficiency, and morphological feature using cryo-electron microscopy (cryo-EM) (Fig. 2a, b). During these freeze-thaw cycles, the particle size of SM-102 LNP gradually increased, especially in the −20 cycles. In contrast, HL LNPs, which initially had a smaller particle size of about 60 nm, showed greater stability in both the −80 cycles and the −20 cycles. Cryo-EM analysis revealed that SM-102 LNPs developed bleb-like structures at the particle edges during the freeze-thaw process, potentially causing the observed increase in particle size and aggregation. In contrast, HL LNPs showed no such structural abnormalities. These differences may be attributed to the stronger interaction between HLs and mRNA, resulting in a more compact and stable particle structure. The enhanced stability of LNP particles suggests that, compared with conventional LNPs, HL LNPs are more resistant to temperature fluctuations during storage, transportation, and use.

**Fig. 2.**
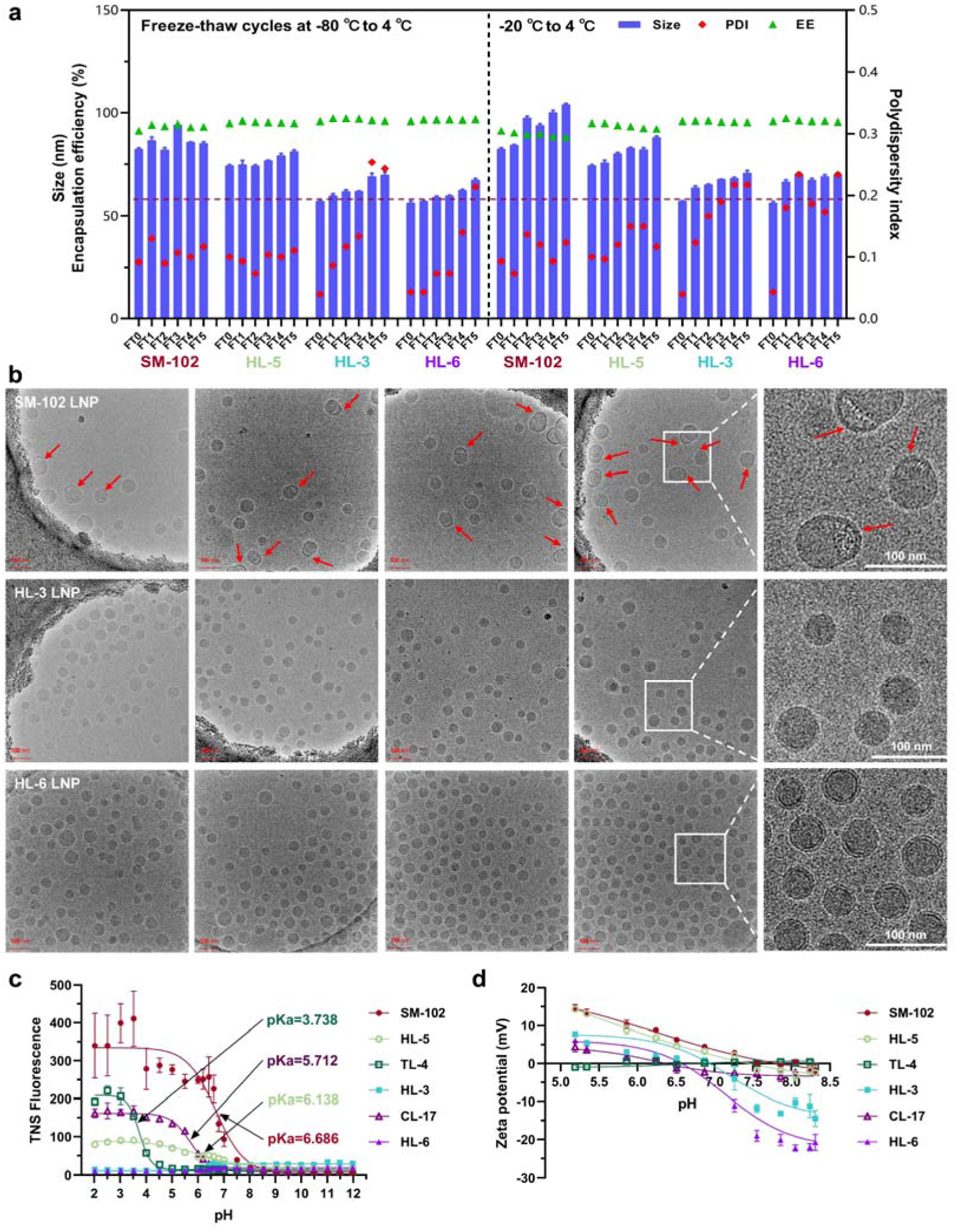
Stability and physicochemical properties of HL LNPs. **a**, Changes in particle size and encapsulation efficiency of LNPs during five freeze-thaw cycles. **b**, Cryo-EM morphology of LNPs after the first freeze-thaw cycle at −20. **c**, Apparent pKa of LNP determined from the TNS fluorescence-pH curve. **d**, Zeta potential-pH curve of LNP over a pH range of 5.2 to 8.3. The data are presented as mean ± SD (n = 3).

The apparent pKa of LNPs is another critical parameter influencing their potential and delivery efficiency, with high-performing LNPs typically showing a pKa around 6.5^37,38^. We determined the apparent pKa of HL LNPs and their tail-controlled counterparts using a TNS-binding assay^38^ (Fig. 2c). Under acidic conditions, the conventional ionizable lipids like SM-102, TL-4, and CL-17 become protonated, which allow TNS to bind to the LNP surface, generating fluorescence. In a basic environment, these LNPs lost their positive charge, and the fluorescence disappeared, enabling the calculation of their apparent pKa from the fluorescence-pH curve. However, this behavior was not observed in HL LNPs. For HL-3 LNP and HL-6 LNP, the TNS fluorescence signal remained at the baseline across the entire pH range, preventing the calculation of the apparent pKa. In HL-5 LNPs, fluorescence was detected, but it was significantly lower than that of its tail control SM-102, suggesting distinct structural properties. These observations indicate that HL LNPs may have a novel particle structure, differing substantially from conventional LNPs. The same pKa trends were observed for empty LNPs formulated without mRNA, confirming the structural differences (Supplementary Fig. 18).

To further characterize the physicochemical differences, we measured the zeta potentials of the LNPs across different pH buffers (Fig. 2d). Using a phosphate buffer ranging from pH 5.2 to 8.3, we observed that the zeta potential curves for conventional LNPs (SM-102, TL-4, CL-17) matched their fluorescence-pH profiles. However, HL-3 and HL-6 LNPs exhibited consistently strong negative zeta potentials in neutral and basic environments, implying a significantly higher negative surface charge compared to their conventional counterparts. For HL-5 LNP, the zeta potential curve aligned with SM-102, likely because of the structural similarity in their tails. These data suggest that the lipid distribution within HL LNPs can be quite different from that in conventional LNPs, despite having the same formulation.

To assess the surface lipid distribution of LNPs, we dialyzed the particles against phosphate-buffered saline (PBS) containing 5% D_2_O and subsequently analyzed the resulting ^1^H NMR spectra. Despite identical formulations, HL-3 and HL-6 LNPs exhibited significant differences in surface group distributions compared to SM-102 and HL-5 LNPs. Specifically, chemical groups corresponding to shifts at approximately 1.5 ppm, 2.4 ppm, and 4.0 ppm were notably reduced in HL-3 and HL-6 LNPs, particularly those associated with tertiary amine and ester-linked methylene groups (Fig. 3a, b). We further quantified the lipid composition on the LNP surface by examining the abundance of hydrogen atoms in the key functional groups of DSPC, DMG-PEG, and ionizable lipids (Fig. 3c and Supplementary Fig. 19-22). The percentage of ionizable lipid present on the surface relative to total lipids was determined to be 74.6% and 71% for SM-102 and HL-5 mRNA-loaded LNPs, respectively. In contrast, this percentage was significantly lower for HL-3 and HL-6, at 45.6% and 29%, respectively. Similar trends were observed for empty LNPs (Fig. 3c and Supplementary Fig. 23-27).

**Fig. 3.**
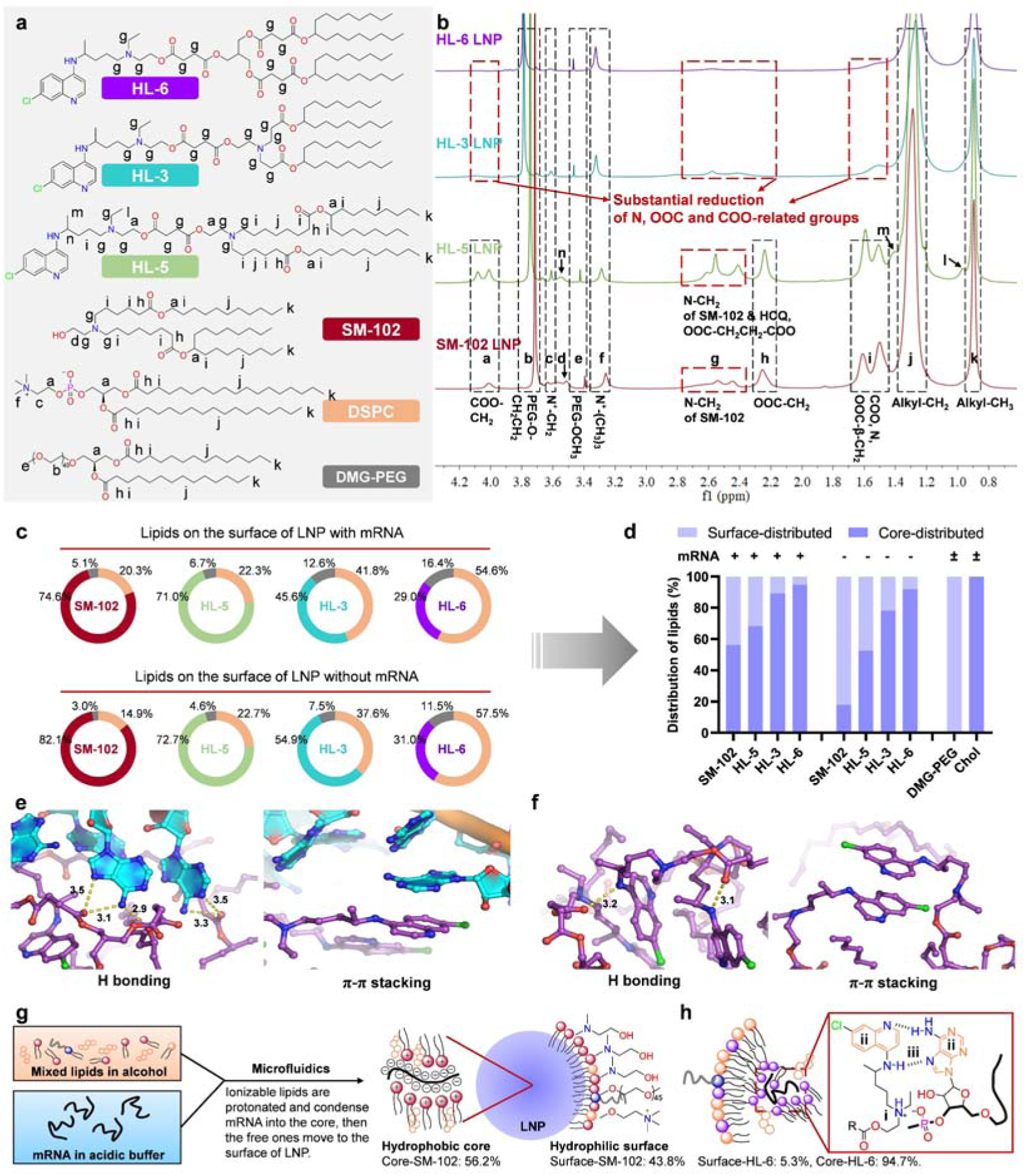
Structural analysis of HL LNPs. **a**, Chemical structures of the lipids distributed on the surface of LNPs. **b**, Merged ^1^H NMR spectra of different LNPs with mRNA. **c**, Proportions (mol/mol) of the three lipids on the surface of different LNPs both with and without mRNA. DSPC is represented in orange, while DMG-PEG is illustrated in grey. **d**, Proportions of the surface- and core-distributed ionizable lipids in LNPs both with and without mRNA. **e**, Non-electrostatic interactions between HL-6 and mRNA. **f**, Non-electrostatic interactions among HL-6 lipids themselves. **g**, Mechanism of LNP formation utilizing conventional ionizable lipids. **h**, Distribution of HLs and interactions between HLs and mRNA: i) electrostatic interaction, ii) π-π stacking interaction, iii) hydrogen bonding.

Given that DMG-PEG is primarily distributed on the LNP surface, the surface-to-core ratio of ionizable lipids was estimated using their ratio to DMG-PEG on the surface and the overall formulation ratio of total ionizable lipid to DMG-PEG (Fig. 3d). For SM-102 LNP, approximately 43.8% of the ionizable lipid was present on the surface of mRNA-loaded LNPs, increasing to 82.1% in empty LNPs. In contrast, for HL-6 LNP, only 5.3% of the ionizable lipid was found on the surface of mRNA-loaded LNPs, with a slight increase to 8.1% in empty HL-6 LNP (Fig. 3d). These findings indicate that HL-6 lipids are more inclined to undergo phase separation and form strong interactions with mRNA, resulting in their predominant localization within the LNP core.

This trend was further supported by molecular dynamics (MD) simulations^39^, which examined interactions between ionizable lipids both in the presence and absence of RNA (Supplementary Fig. 28). MD simulations revealed extensive interactions between HL lipids and RNA (Fig. 3e), as well as between HL lipids themselves (Fig. 3f). These interactions were primarily mediated by hydrogen bonding and significant π-π stacking involving the aminoquinoline groups and RNA bases. This distinctive interaction profile highlights the unique potential of HL-functionalized lipids, distinguishing them from conventional lipid formulations. Consequently, the strong affinity between HL lipids and mRNA results in fewer HL lipids being available on the LNP surface (Fig. 3g, h), which contributes to unique characteristics such as reduced TNS responsiveness and a more pronounced negative zeta potential during mRNA encapsulation.

### Assessment of mRNA expression and immune response under repeated administration

As the mRNA-LNP system gain wider application, particularly beyond vaccines to fields such as protein replacement therapies, repeated dosing becomes increasingly relevant. However, repeated dosing also raises concerns about inflammatory side effects, which can compromise mRNA expression and therapeutic efficacy^21^. To address these challenges, we investigated mRNA expression, cytokine release, and safety profiles of HL LNPs under repeated dosing conditions.

BALB/c mice were administered FLuc mRNA-loaded LNPs intravenously at a dose of 5 μg of mRNA per mouse, for a total of four injections, with a three-day interval between each administration (Fig. 4a). Bioluminescence signals were recorded at 4 h and 72 h following each injection. Blood samples were collected at 4 h after the first and second injections to measure serum levels of key proinflammatory cytokines, including TNF-α, IFN-γ, IL-6, and monocyte chemoattractant protein-1 (MCP-1). In addition, the body weights of the mice were monitored throughout the course of repeated administration, with weight growth curve plotted to assess overall health (Supplementary Fig. 29).

**Fig. 4.**
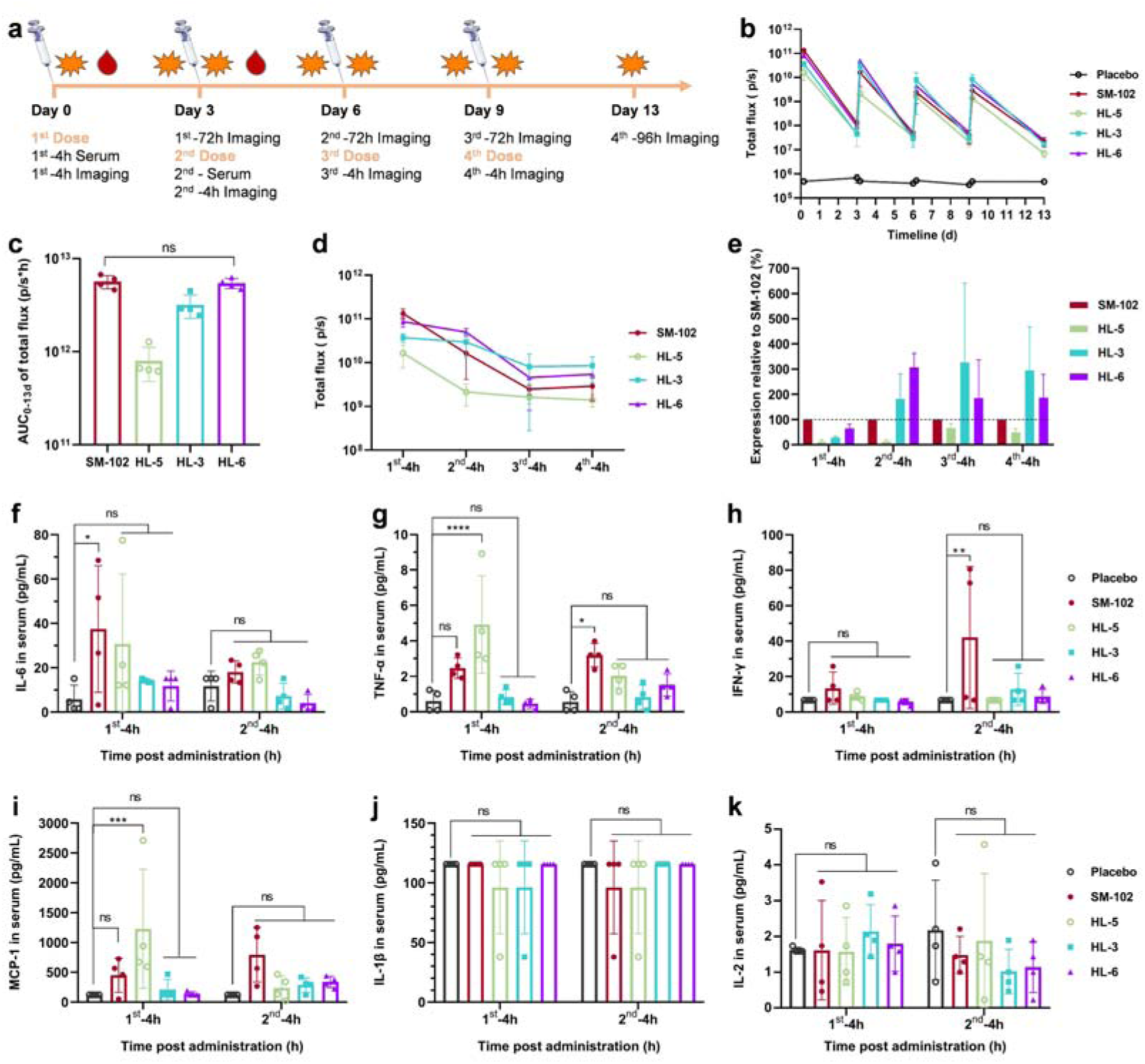
Repeated administration of luciferase mRNA loaded in different LNPs. **a**, Scheme for the repeated administration, sampling and detection. **b**, Bioluminescence-time curve during the repeated administration. **c**, AUC_0-13d_ following the repeated administration. **d**, Bioluminescence observed 4 hours following each administration. **e**, The ratio of mRNA expression in the HL groups relative to that of SM-102 at 4 h after each administration. **f-k**, Serum concentrations of proinflammatory cytokines at 4 h after administration, including IL-6, TNF-α, IFN-γ, MCP-1, IL-1β and IL-2. The dispersing agent for LNP, a Tris-acetate buffer containing 8% sucrose (8%Su Tris-Ac), was utilized as a placebo. The data are presented as mean ± SD (n = 4). The P values were determined by one-way ANOVA with one-sided Dunnett’s test. (ns, not significant; *, p < 0.0332; **, p < 0.0021; ***, p < 0.0002; ****, p < 0.0001.)

To visualize the kinetics of mRNA expression across the repeated administration regimen, bioluminescence signals over 13 days were plotted (Fig. 4b) and the AUC_0-13d_ was calculated for each group (Fig. 4c). Additionally, mRNA expression levels at 4 h post-injection were extracted and plotted separately (Fig. 4d and Supplementary Fig. 30), and the ratio of mRNA expression in the HL groups to the SM-102 group was calculated for each time plot (Fig. 4e). Despite a gradual decline in mRNA expression across all groups with the repeated administration, the decrease was significantly less pronounced in the HL-3 and HL-6 groups compared to the SM-102 group (Fig. 4d). By the second dose, mRNA expression levels in the HL-3 and HL-6 groups surpassed those of SM-102, exhibiting expression levels 2-3 time higher (Fig. 4e). While the initial expression after the first dose was higher in the SM-102 group, the AUC for the HL-6 group remained comparable to SM-102 over the entire course of repeated administration, highlighting the sustained expression levels achieved by HL LNPs.

Cytokine analysis revealed that SM-102 LNP and HL-5 LNP triggered significantly higher levels of cytokines, including IL-6, TNF-α, IFN-γ, and MCP-1, compared to HL-3 and HL-6 LNPs (Fig. 4f-k). The elevated cytokine release observed in the SM-102 and HL-5 groups likely contributed to reduced mRNA translation during repeated dosing. In contrast, HL-3 and HL-6 LNPs induced substantially lower levels of cytokines, suggesting a more controlled inflammatory response and better mRNA translation efficiency over time. Interestingly, HL-5, which is a conjugate of HCQ and SM-102, was as inflammatory as SM-102, suggesting that the SM-102 component can trigger a strong inflammatory response even when combined with HCQ.

### Mechanistic validation of immunosuppression by HL Lipids

To better understand how HCQ-derived ionizable lipids suppress inflammatory cytokine release, we examined their effect on innate immune signaling pathways in vitro using THP1-Dual cells. These cells are widely used to study the activation of the nuclear factor κB (NF-κB) and interferon regulatory factor (IRF) pathways, which can be monitored via secreted embryonic alkaline phosphatase (SEAP) and Lucia luciferase^40^, respectively. To stimulate a strong immune response, high molecular weight polyinosinic-polycytidylic acid (poly(I:C)), a strong TLR agonist, was co-delivered with FLuc mRNA at a mass ratio of 1:5 in LNPs and transfected into THP1-Dual cells. Lipofectamine 2000 (Lipo2000) and the LNP solvent (8% sucrose Tris-acetate buffer) served as controls. 24 hours post-transfection, we evaluated cell viability (Fig. 5a), FLuc expression (Fig. 5a), the activity of Lucia luciferase (Fig. 5b) and activity of SEAP (Fig. 5c). Poly(I:C)-loaded SM-102 LNP activated both the IRF and NF-κB pathways, as evidenced by a significant increase in SEAP and Lucia luciferase activity. In contrast, HL LNPs effectively suppressed this activation, indicating their ability to mitigate immune activation induced by poly(I:C). In comparison, when using Lipo2000, the strong immune stimulation caused a marked reduction in cell viability and mRNA expression, aligning with the in vivo results observed during repeated administration.

**Fig. 5.**
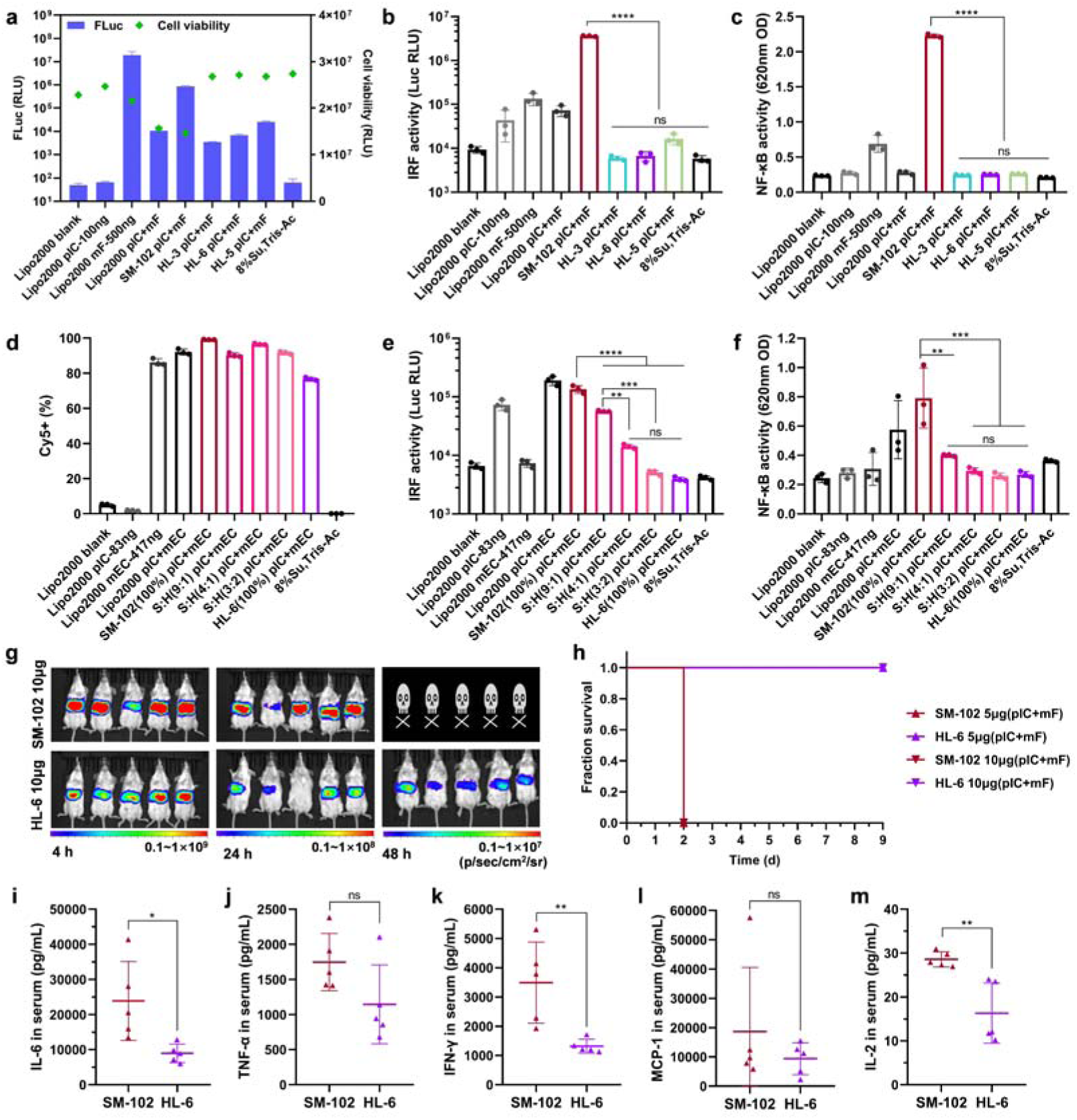
Validation of HLs blocking IRF and NF-κ B pathways and protecting mice from the toxicity of poly(I:C). THP1-Dual cells were co-transfected with poly(I:C) (pIC) and FLuc mRNA (mF) at a mass ratio of 1:5 using HL LNPs, SM-102 LNP or Lipo2000. **a**, Expression of FLuc mRNA and cell viability 24 h post-transfection. **b**, Activity of IRF 24 h post-transfection. **c**, Activity of NF-κB 24 h post-transfection. Hybrid LNPs loaded with poly(I:C) and Cy5-labeled EGFP mRNA (mEC) were formulated using specific ratios of SM-102 (S) and HL-6 (H). **d**, Cy5-positive rate 24 h post-transfection. **e**, Activity of IRF 24 h post-transfection. **f**, Activity of NF-κB 24 h post-transfection. **g**, Bioluminescence images of the high dose groups. **h**, Kaplan-Meier curves of both 5 μg and 10 μg groups. **i-m**, Serum concentrations of proinflammatory cytokines at 4 h after administration in the 5 μg groups, including IL-6, TNF-α, IFN-γ, MCP-1 and IL-2. The data are presented as mean ± SD (n = 3 for in vitro, and n = 5 for in vivo). The P values were determined by one-way ANOVA with one-sided Dunnett’s test. (ns, not significant; *, p < 0.0332; **, p < 0.0021; ***, p < 0.0002; ****, p < 0.0001.)

Next, to explore whether the differences in immune activation were due to disparities in mRNA uptake, we generated hybrid LNPs combining SM-102 and HL-6 at various ratios, which were loaded with poly(I:C) and Cy5-labeled EGFP mRNA at the same mass ratio of 1:5. These hybrid LNPs were then transfected into THP1-Dual cells. The comparable Cy5-positive rates in different hybrid LNPs indicated that the amounts of poly(I:C) internalized by cells in different groups were almost the same (Fig. 5d). Despite similar uptake, SM-102 LNP loaded with poly(I:C) strongly activated the innate immune pathways. However, as the proportion of HL-6 in the hybrid LNPs increased, the activity of both the IRF and NF-κB pathways gradually decreased (Fig. 5e, f), indicating that HL-6 has a clear inhibitory effect on innate immune pathways.

To further validate the in vivo inhibitory effects of HL lipids on poly(I:C)-induced immune activation, we conducted a toxicity challenge experiment using LNPs loaded with a mixture of poly(I:C) and FLuc mRNA at a 1:1 mass ratio. Each mouse received a single intravenous injection of either 5 μg or 10 μg of the nucleic acid mixture loaded in SM-102 LNP or HL-6 LNP (Fig. 5g-m). Proinflammatory cytokine analysis was performed on blood collected 4 hours post-administration, and bioluminescence imaging was conducted at 4, 24, 48, and 120 hours following injection. Subsequently, the survival of mice was monitored. No significant difference in luciferase expression was observed between the SM-102 and HL-6 groups at either dose (Fig. 5g and Supplementary Fig. 31). However, survival curves exhibited a striking difference (Fig. 5h). While the SM-102 groups experienced complete mortality within 48 hours, even at the lower dose (5 μg), the HL-6 groups maintained 100% survival, even at the higher dose (10 μg), demonstrating the superior protective efficacy of HL lipids against poly(I:C) toxicity. In contrast to the repeated administration of mRNA alone (Fig. 4f-k), the addition of poly(I:C) strongly stimulated the immune system, triggering a cytokine storm with an approximately 800-fold increase in IL-6 and a 100-fold increase in IFN-γ, indicating the lethal effects of the high poly(I:C) dose. Cytokine profiling at the 5 μg dose revealed that the levels of proinflammatory cytokines, including IL-6, IFN-γ, and IL-2, were significantly lower in the HL-6 group compared to the SM-102 group (Fig. 5i-m). This advantage diminished at the 10 μg dose, potentially due to excessive poly(I:C) dosage (Supplementary Fig. 32). These in vivo results strongly support the in vitro findings, demonstrating that HL LNPs effectively suppress nucleic acid-induced innate immune activation, thereby reducing the release of proinflammatory cytokines.

### Immunogenicity of HL LNP-based mRNA vaccine against VZV

The proinflammatory cytokines responses induced by mRNA vaccines remains a significant concern, particularly for prophylactic vaccines where excessive immune activation can be detrimental. To explore the potential of HL LNPs as a safer and more effective delivery platform, we developed an mRNA vaccine targeting varicella-zoster virus (VZV). VZV is a member of the herpesvirus family responsible for both chickenpox (varicella) and shingles (zoster)^41^. We selected glycoprotein E (gE) as the target antigen, as it has been proven to be effective in approved subunit vaccine Shingrix (GlaxoSmithKline)^42^.

To assess the immunogenicity and safety of HL-based mRNA vaccines, we formulated gE mRNA into HL-3, HL-6, and SM-102 LNPs, yielding HL-3-gE, HL-6-gE, and SM-102-gE vaccines. C57BL/6 mice were immunized intramuscularly with doses of 2 μg mRNA each, following a prime-boost schedule with a three-week interval between doses (Fig. 6a). Serum samples were collected three weeks post-prime dose and two weeks post-boost dose to measure gE-specific antibody titers. The results showed that both HL-3-gE and HL-6-gE induced robust humoral immune responses, with antibody titers comparable to those elicited by SM-102-gE (Fig. 6b). This demonstrates that even at a relatively low dose, HL LNPs can drive a strong humoral response.

**Fig. 6.**
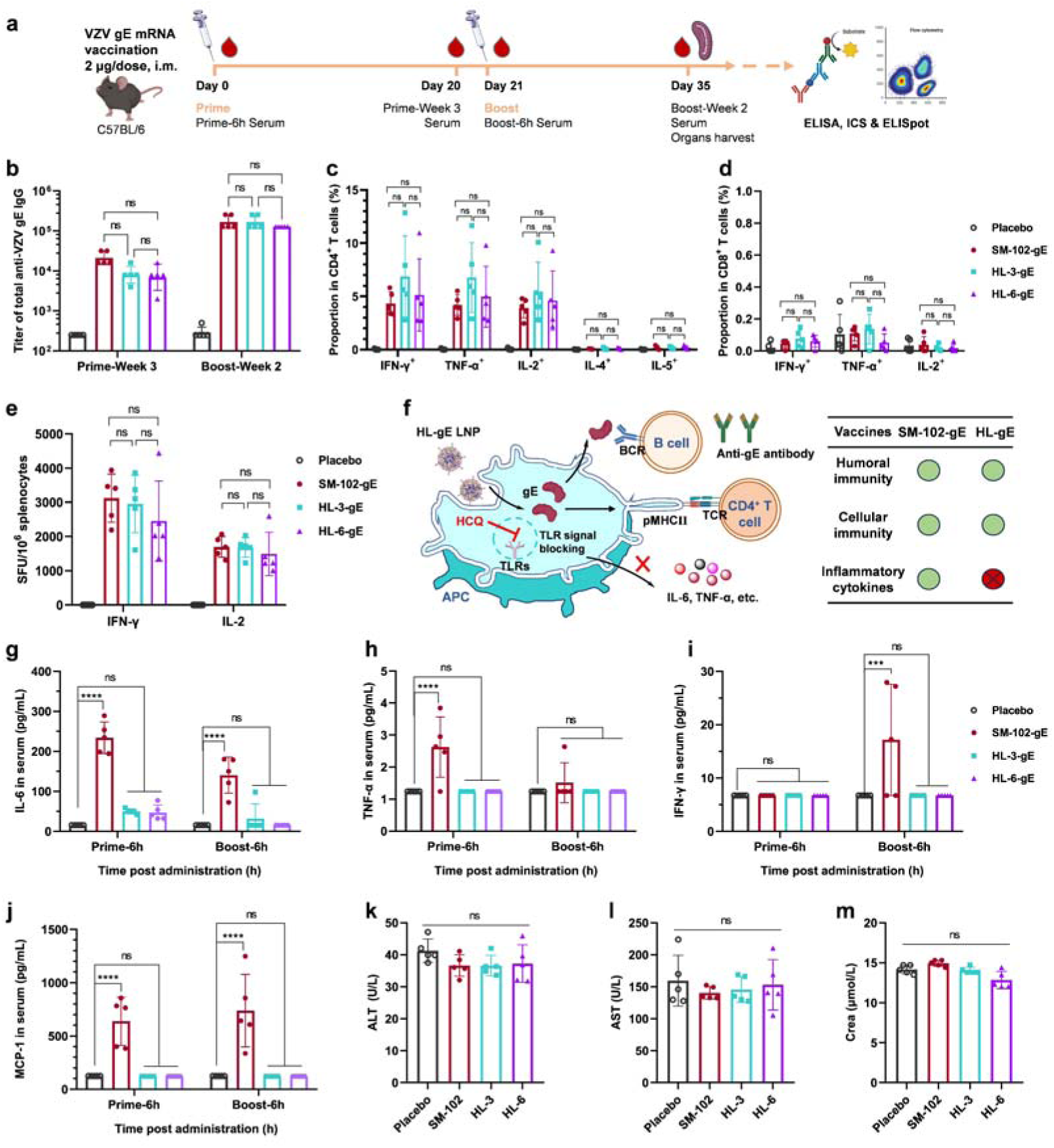
HL LNP-based gE mRNA vaccine against VZV. **a**, Scheme for vaccination and sample collection. **b**, Endpoint dilution titers of total gE-specific IgG antibody. **c**, Antigen-specific CD4^+^ T cells expressing different cytokines in spleen quantified by ICS. **d**, Antigen-specific CD8^+^ T cells expressing different cytokines in spleen quantified by ICS. **e**, Antigen-specific IFN-γ-secreting and IL-2-secreting cells in spleen measured by ELISpot. **f**, Illustration of HL-gE vaccines. After LNPs are taken in by antigen presenting cells (APCs), the gE protein is expressed, and then it is either secreted to be recognized by B cells, or processed into antigen peptides to be presented to T cells, thus triggering adaptive humoral immunity and cellular immunity. At the same time, HCQ blocks the TLR signal pathway and inhibits the production of inflammatory cytokines. **g-j**, Release of proinflammatory cytokines post each immunization, including IL-6, TNF-α, IFN-γ and MCP-1. **k-m**, Blood biochemistry at Week 2 post the booster dose, including alanine aminotransferase (ALT), aspartate aminotransferase (AST) and creatinine (Crea). The data are presented as mean ± SD (n = 5). The P values were determined by one-way ANOVA with one-sided Dunnett’s test. (ns, not significant; *, p < 0.0332; **, p < 0.0021; ***, p < 0.0002; ****, p < 0.0001.)

We next evaluated the cellular immune responses through intracellular cytokine staining (ICS) and enzyme-linked immunosorbent spot (ELISpot) assays two weeks after the boost dose. Splenocytes from vaccinated mice were stimulated with a gE peptide pool to quantify the presence of cytokine-secreting CD4^+^ (Fig. 6c) and CD8^+^ T cells (Fig. 6d). The gating strategy for flow cytometry was shown in Supplementary Figure 33. Both HL-3-gE and HL-6-gE vaccines generated strong gE-specific CD4^+^ T cells, characterized by Th1-type cytokine production, including IFN-γ, IL-2, and TNF-α. Interestingly, HL-3-gE led to a significantly higher percentage of gE-specific CD4+ T cells compared to SM-102-gE (Supplementary Fig. 34), while none of the vaccines elicited significant Th2 cytokine responses (e.g., IL-4 and IL-5), indicating a robust Th1-biased immune profile. Moreover, the gE-specific T cell response was predominantly mediated by CD4^+^ T cells with minimal CD8^+^ T cell response (Supplementary Fig. 34), consistent with previous reports^43,44^. These trends were corroborated by IFN-γ and IL-2 ELISpot assays, which showed similar results (Fig. 6e and Supplementary Fig. 35), further validating that HL LNPs promote strong antigen-specific Th1-biased T cell immunity.

In addition to immune response analysis, we examined proinflammatory cytokine levels in serum collected 6 hours post-immunization. SM-102-gE vaccine induced significant levels of proinflammatory cytokines, such as IL-6, TNF-α, IFN-γ, and MCP-1, even at a low dose of 2 μg mRNA. In contrast, the cytokine levels in the HL-3-gE and HL-6-gE groups were similar to those in the placebo group (Fig. 6g-j and Supplementary Fig. 36), further echoing with previous results. This suggests that HL LNPs effectively suppress the production of excessive proinflammatory cytokines, while still enabling potent immune responses (Fig. 6f).

Two weeks after the booster immunization, mice were sacrificed, and the blood and organs were collected for histopathological analysis and blood biochemistry evaluation. Throughout the immunization process, body weight was also monitored to assess any adverse effects. All results showed no significant abnormalities in mice treated with HL-based vaccines, further confirming the safety profile of HL LNPs (Fig. 6k-m and Supplementary Fig. 37, 38).

### Gene editing of PCSK9 via CRISPR-Cas9 mRNA delivered by HL LNPs

To further demonstrate the versatility of HL LNPs for mRNA delivery beyond vaccines, we investigated their potential in gene editing through CRISPR-Cas9. Specifically, we targeted the proprotein convertase subtilisin/kexin type 9 (PCSK9) gene, which plays a key role in regulating low-density lipoprotein receptor (LDLR) degradation and cholesterol metabolism. Disruption of PCSK9 through CRISPR-Cas9 can lower cholesterol levels by enhancing LDLR expression and promoting cholesterol clearance from the bloodstream^45,46^.

To create a model of hypercholesterolemia, C57BL/6 mice were fed a high-fat diet for two months. Each model mouse was then intravenously injected with Cas9 mRNA and sgPCSK9^47^, loaded into HL-6 or SM-102 LNPs, at a dose of 3 mg per kg. Mice were euthanized 14 days post administration, and the serum and liver samples were collected. HL-6 LNP loaded with EGFP mRNA was used as a negative control. Genomic DNA from the liver was extracted, and insertion/deletion (Indel) changes in the PCSK9 gene were analyzed using next-generation sequencing (NGS) (Fig. 7a, e). In both treatment groups, PCSK9 was significantly knocked down, with editing efficiency reaching up to approximately 30%.

**Fig. 7.**
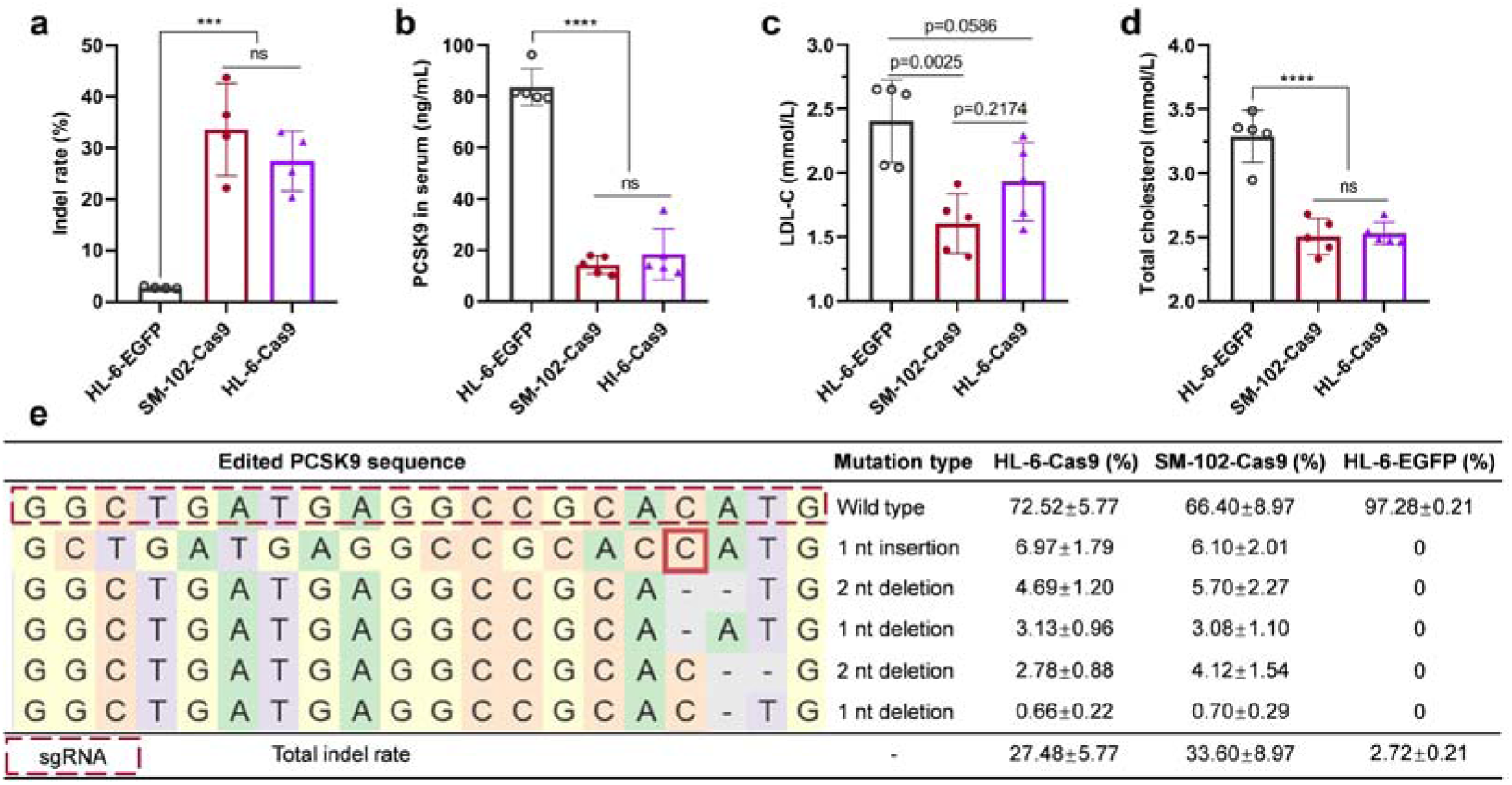
CRISPR-Cas9 mRNA-based gene editing of PCSK9. **a**, NGS analysis of the total indel rate of PCSK9. **b**, Concentration of PCSK9 protein in serum. **c**, Concentration of LDL-C in serum after PCSK9 gene editing. **d**, Concentration of TC in serum after PCSK9 gene editing. **e**, Mutation type and frequency of different indels. The data are presented as mean ± SD (n = 5). The P values were determined by one-way ANOVA with one-sided Dunnett’s test. (ns, not significant; *, p < 0.0332; **, p < 0.0021; ***, p < 0.0002; ****, p < 0.0001.)

Compared to the negative control group, the serum PCSK9 concentrations in the HL-6-Cas9 and SM-102-Cas9 groups decreased sharply by 78% and 83%, respectively (Fig. 7b). In addition, LDL cholesterol (LDL-C) and total cholesterol (TC) levels in the serum were also reduced, indicating that PCSK9 editing exerted a cholesterol-lowering effect (Fig. 7c, d). These results demonstrate that HL-6 LNPs are comparable to the commercially available SM-102 LNP in terms of gene editing efficacy with Cas9 mRNA, supporting the potential of HL LNPs for broader applications in gene editing.

## Discussion and conclusion

In this study, we introduce a novel approach to mitigate the inflammatory side effects of LNPs by incorporating hydroxychloroquine (HCQ) into the design of ionizable lipids. HCQ has long been recognized for its immunomodulatory properties, particularly in the treatment of autoimmune disorders, cancer, and infectious diseases such as COVID-19. Its mechanism of action includes the inhibition of endosomal Toll-like receptors (TLRs)—specifically TLR3, TLR7, and TLR9, which recognize nucleic acids such as DNA and double-stranded RNA (e.g., poly(I:C)). HCQ impairs these receptors by blocking endosomal acidification through its ionizable group at high concentrations. Additionally, it can directly bind nucleic acids, altering their structure and preventing their interaction with TLRs, even at lower concentrations that do not affect endosomal pH. HCQ also blocks cyclic GMP-AMP synthase (cGAS) binding, reducing the production of IFN-β and pro-inflammatory cytokines.

Leveraging these properties, we investigated whether HCQ could serve as a foundation for designing ionizable lipids for mRNA delivery. Our objectives were threefold: (1) to assess whether HCQ’s nucleic acid-binding properties could improve the encapsulation and stability of nucleic acids within LNPs; (2) to explore whether HCQ’s intrinsic ionizable group could facilitate mRNA delivery in a manner similar to established ionizable lipids; and (3) to determine whether HCQ could mitigate the inflammatory side effects commonly associated with LNP systems.

To achieve this, we synthesized HCQ-derived ionizable lipids (HLs). Our findings showed that HL-3 and HL-6, which do not contain additional functional nitrogen, were effective in encapsulating and delivering mRNA both in vitro and in vivo. Notably, HL-LNPs displayed a highly stable structure, likely due to enhanced interactions between the mRNA and the aminoquinoline moiety of HCQ, in addition to the positive charge from its ionizable nitrogen. NMR analysis revealed that HLs were significantly less distributed on the surface of LNPs compared to SM-102, resulting in a negative zeta potential across a wide pH range. This structural stability suggests that HL-LNPs have an improved safety profile, as lower surface distribution of ionizable lipids may reduce recognition by the innate immune system.

Both HL-3 and HL-6 exhibited significantly lower inflammatory responses compared to SM-102, which is widely used in current mRNA vaccines. Interestingly, when HCQ was conjugated to SM-102 in HL-5, it did not suppress the activation of inflammatory cytokines, suggesting that HCQ alone could not counteract the inherent pro-inflammatory properties of SM-102. This highlights the importance of the lipid architecture in determining the inflammatory potential of LNP formulations.

Repeated administration of SM-102 LNPs led to significant cytokine release and reduced mRNA expression. In contrast, HL-3 and HL-6 LNPs not only suppressed cytokine production but also sustained mRNA expression over multiple doses. This finding underscores the potential of HL-lipids in improving the tolerability and efficacy of mRNA-based therapies, particularly in contexts requiring repeated dosing.

Further studies using poly(I:C), a model nucleic acid, confirmed that HL-lipids effectively suppressed nucleic acid-induced immune activation and protected mice from the toxicity of poly(I:C), in line with previous findings that HCQ inhibits TLR signaling by directly binding nucleic acids. Importantly, this suppression did not impair adaptive immune responses, as demonstrated by robust B-cell and T-cell responses in a VZV vaccine model. This indicates that HL-LNPs may be particularly suitable for a wide range of nucleic acid delivery applications, including DNA and self-amplifying RNA (saRNA), where excessive innate immune activation is undesirable.

We also observed that the tail structure of the HL-lipids significantly influenced their encapsulation and delivery efficiency. This suggests that further optimization of lipid tail design could enhance delivery performance. The ability of HLs to suppress immune responses triggered by exogenous nucleic acids positions them as a promising system for delivering DNA and saRNA, both of which are often associated with strong inflammatory responses. Future research will focus on this aspect.

In conclusion, the integration of HCQ into ionizable lipids provides a promising strategy for reducing inflammation in LNP systems while maintaining effective mRNA delivery. HCQ’s dual role—facilitating endosomal escape and reducing immune activation—combined with the favorable structural properties of HL-LNPs, makes them a strong candidate for a wide range of therapeutic applications, including gene editing and protein replacement therapies.

## Methods

### Materials

Chemicals used in the synthesis of lipids: 2-decyltetradecan-1-ol (58670-89-6, Bidepharm), succinic anhydride (108-30-5, Macklin), 4-dimethylaminopyridine (1122-58-3, Bidepharm), hydroxychloroquine (118-42-3, Bidepharm), 1,3-dicyclohexylcarbodiimide (538-75-0, Bidepharm), heptadecan-9-ol (624-08-8, Bidepharm), 2-heptylundecanoic acid (22890-21-7, Macklin), triethylamine (121-44-8, Sinopharm), acryloyl chloride (814-68-6, Adamas), ethanolamine (141-43-5, Adamas), SM-102 (2089251-47-6, RNACure), 2-hydroxymethyl-1,3-propanediol (4704-94-3, Bidepharm), 2-diethylaminoethanol (100-37-8, Adamas). For the formulation of LNPs, 1,2-distearoyl-sn-glycero-3-phosphocholine (DSPC, 816-94-4) and cholesterol were procured from Nippon Fine Chemical Co., Ltd., while 1,2-dimyristoyl-rac-glycero-3-methoxypolyethylene glycol-2000 (DMG-PEG, 160743-62-4) was obtained from Guobang Pharma Group Co., Ltd. For the determination of LNP apparent pKa, 6-(p-toluidino)-2-naphthalenesulfonic acid sodium salt (TNS, 53313-85-2) was acquired from Nanjing Xize Pharmaceutical Technology Co., Ltd. Poly(I:C) (31852-29-6, size: 1.5 - 8 kb) was purchased from InviGen (catalog code: vac-pic). The antibodies, assay kits, and other reagents can be found in the corresponding method sections.

### Synthesis of lipid compounds

The following is a brief summary of the synthesis steps of HLs: i) The hydrophobic tails with carboxyl heads were formed by the ring-opening reaction of succinic anhydride and the esterification of carboxylic acid; ii) The esterification reaction between the carboxylic acid-headed tail and the hydroxyl group of HCQ produced the final products. The lipid compounds were purified using silica gel column chromatography, and their structures were verified by NMR and MS. The production procedures, synthesis routes, NMR spectra and MS of all lipid compounds are listed in the Supplementary Information.

### mRNA synthesis

The FLuc mRNA, VZV gE mRNA, and Cas9 mRNA were synthesized in vitro by T7 RNA polymerase-mediated transcription from a linearized DNA template. Modified uridine-5′-triphosphate (1-methylpseudouridine) was incorporated in a certain ratio to reduce the immunogenicity of the mRNA and Cap1 was employed to enhance the translation efficiency.

### LNP preparation

Ionizable lipid (HLs, TL-4, CL-17 or SM-102), DSPC, cholesterol, DMG-PEG were co-dissolved in alcohol at a mol ratio of 50:10:38.5:1.5, resulting in a total lipid concentration of about 10 mM. The mRNA (with or without poly(I:C)) was dissolved in a 25 mM sodium acetate buffer with a pH of 5.0. The aqueous phase and the alcohol phase (at an ionizable lipid/nucleotide ratio of 6:1) were mixed using a microfluidic device at a volume ratio of 3:1 with a total flow rate of 12 mL/min. The resulting formulations were dialyzed against 20 mM Tris-acetate buffer (pH∼7.5) to remove alcohol using Slide-A-Lyzer dialysis cassettes (10 kDa MWCO, Thermo Fisher). Following dialysis, the solutions were concentrated using Amicon ultra-centrifugal filters (10 kDa MWCO, Millipore) and subsequently filtered through a 0.22 μm filter. LNPs without mRNA were prepared using the sodium acetate buffer devoid of mRNA, and the same procedures for preparing LNPs with mRNA were followed. For the SM-102/HL-6 hybrid LNPs, the mol ratios were as follows: S/H(9:1) = 45:5:10:38.5:1.5, S/H(4:1) = 40:10:10:38.5:1.5, S/H(3:2) = 30:20:10:38.5:1.5.

### LNP characterization

The particle size and PDI of LNPs were measured using a Zetasizer Pro (Malvern). The encapsulation efficiency of mRNA within the LNPs was determined using a Quant-iT RiboGreen RNA assay kit (Invitrogen, R11490). The concentration of mRNA was quantified using a Stunner analyzer (Unchained Labs). The apparent pKa of the LNPs was determined through the TNS binding assay.

### Freeze-thaw challenges

LNPs were prepared and dispersed in a Tris-acetate buffer added with 8% sucrose. In each freeze-thaw cycle, the LNPs were subjected to a freezing temperature of −20 or −80 for a minimum of 20 h, followed by thawing at 4 for about 2 h. The particle size, PDI and encapsulation efficiency were determined, and the morphology of the LNPs was assessed using a Cryo-EM microscope.

### Cryo-EM

Cryo-EM specimens were prepared on the ANTcryoTM Au300 R1.2/1.3 grids (Nanodim). The grids were glow-discharged using an easyGlow device (PELCO) with the parameters set to 15 mA for a duration of 100 seconds. A Vitrobot plunge freezer (Vitrobot Mark IV, Thermo Fisher) was utilized for preparing cryo grids. A droplet of 3 μL was applied to the freshly glow-discharged grid, and the grid was subsequently blotted (bolt time: 4 s, bolt force: 0, wait time: 30 s) and plunge-frozen in liquid ethane, then stored in liquid nitrogen until further use. Autogrids were assembled under liquid nitrogen and then loaded into the electron microscope for further examination. Data collection was performed using a Glacios transmission electron microscope (Thermo Fisher) operating at 200 kV, equipped with a Ceta-D camera. Images were captured using the EPU software (Thermo Fisher).

### ^1^H-NMR spectrum of LNP

The lipid distribution on the surface of LNP was characterized using an ultra-high field NMR spectrometer (800 MHz). The LNPs were dialyzed into 0.2 × PBS, followed by adding 5% D_2_O for field locking. By combining the chemical shifts of each lipid with two-dimensional NMR spectra, the spectral peaks corresponding to each lipid were identified.

### Analysis of ionizable lipid distribution

The lipid composition on the surface of LNPs was quantitated by analyzing the spectrum abundance of hydrogen (H) in the -N+(CH_3_)_3_ group of DSPC (n = 9, theoretically), the –(OCH_2_CH_2_)_45_-group (n = 180, theoretically) and the -OCH_3_ group of DMG-PEG, and the tertiary amine-(e.g., for SM-102, n = 6, theoretically) and the ester-related groups from ionizable lipids.

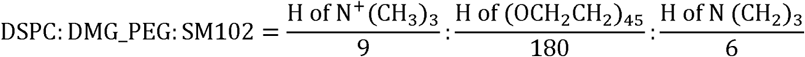

Given that all the DMG-PEG was distributed on the LNP surface, the surface-to-core ratio of ionizable lipids within LNPs could be estimated according to the ratio of surface-distributed ionizable lipids to DMG-PEG and the overall formulation ratio of total ionizable lipid to DMG-PEG (50:1.5).

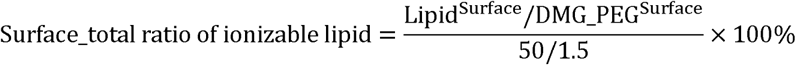

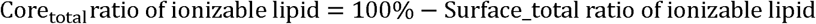

### Molecular dynamics simulations

MD trajectories were acquired using Gromacs2022 package, GPU-accelerated, employing the AMBER14SB and GAFF all-atom forcefields. Each simulation box, maintained at a 1:1 base to lipid ratio, housed a 10-nucleotide mRNA, 10 ionizable lipids either HL-5/SM-102 or HL-6/CL-17, and was solvated with explicit TIP3P water molecules within a vacuum box placed 2 nm away from the border. For investigation of self-interactions, the box only housed ionizable lipids, without mRNA. The systems were initialized with bond length, angle, and dihedral values for the lipids generated using Sobtop, and the RESP charges computed via Gaussian and Multiwfn. Following energy minimization using a steepest descent approach, a pre-equilibration phase was conducted through a 100 ps isothermal-isobaric simulation (NPT). The subsequent 100 ns production MD simulation was carried out, and a coupling constant of 2.0 ps, in alignment with the Parrinello−Rahman algorithm. The temperature was held steady at 298.15 K, with a coupling constant of 0.2 ps as per v-rescale algorithm. Short-range interactions were set with a 1.0 nm cutoff, while the Particle Mesh Ewald algorithm was employed for long-range electrostatic interactions. Trajectory snapshots were recorded every 10 ps, documenting dynamic integrals at 2 fs intervals. Analysis and visualization of the motion trajectories, hydrogen bond characteristics, and conformational changes were facilitated using Visual Molecular Dynamics and Qtgrace tools. Model representations in the figures and movie were prepared by PyMOL.

### THP1-Dual cells

THP1-Dual cells were purchased from InvivoGen, which were derived from the human THP-1 monocyte cell line by stable integration of two inducible reporter constructs. These cells comprise a secreted Lucia luciferase reporter gene driven by an ISG54 promoter and also a NF-κB–inducible secreted SEAP gene. THP1-Dual cell lines were cultured in RPMI 1640 growth medium (containing 2 mM L-glutamine and 25 mM HEPES, Gibco), supplemented with 10% heat-inactivated FBS (Gibco), 100 μg mL^-1^ Normocin (InvivoGen), 100 μg mL^-1^ Pen-Strep (SclenCell), and selective antibiotics Blasticidin (10 μg mL^-1^, InvivoGen) and Zeocin (100 μg mL^-1^, InvivoGen).

### THP1-Dual reporter assays

THP1-Dual cells were seeded in flat-bottom 96-well plates (Nest) at 50,000 cells per well in Normocin-free and selective antibiotics-free RPMI 1640 growth media with 10% HI-FBS before transfection. The cells were transfected with LNPs encapsulating mRNA (500 ng per well) for 24 h. The corresponding mRNA and the positive control were transfected using Lipofectamine 2000 (Thermo Fisher). Additionally, empty LNPs, containing the same lipid concentration (∼9.23 mM ionizable lipids) as the LNPs encapsulating mRNA, were introduced into each well. 24 h post-transfection, the cells were centrifuged at 300 g for 5 min, and the supernatant was collected for the detection of the secreted proteins expressed by the two reporter genes. Luciferase activity was quantified using the QUANTI-Luc 4 assay (InvivoGen), while secreted embryonic alkaline phosphatase (SEAP) levels were measured using the QUANTI-Blue assay (InvivoGen) on a microplate reader (Varioskan LUX, Thermo Fisher).

### Uptake and expression of HL LNPs in vitro

In order to further assess the immunosuppressive capacity of HL LNPs on the RNA they deliver, we conducted an examination of the uptake and expression capabilities of each LNP in THP1-Dual cells, while simultaneously monitoring the associated immune responses. The number of seeded cells and the transfection dose of LNPs were consistent with those used in assays of innate immune reporter genes. Poly(I:C) and mRNA (FLuc or Cy5-labeled EGFP mRNA) were mixed with a mass ratio of 1:5 and encapsulated within HL LNPs to investigate the correlation between cellular uptake, protein expression, and the immune response elicited by mRNA-encapsulated HL LNPs.

24 h post-transfection with LNPs, the supernatant of the culture medium was utilized for the detection of immune responses as previously described. The cells were subsequently employed for assays assessing cellular uptake and expression. The luciferase expression of FLuc mRNA-transfected cells were measured using a microplate reader (Varioskan LUX, Thermo Fisher). For the transfection of Cy5-EGFP mRNA, the cells were washed twice with 100 μL PBS, and each sample was then resuspended in 50 μL PBS containing Fixable Viability Stain 510 Stock Solution (diluted at 1:500, BD Bioscience) to achieve a concentration of 1,000,000 cells mL^-1^. The samples were transferred to 96-well V-bottom plates and incubated at room temperature for 10 min, followed by the addition of 150 μL of the stain buffer. After the removal of the supernatant, the cells were resuspended in 150 μL of PBS. The percentage and mean fluorescence intensity (MFI) of Cy5 or GFP-positive cells among viable cells were analyzed using a flow cytometer (Cytek Aurora NL).

### Mice

Female BALB/c mice, weighing between 18-22 g, were procured from Sipeifu (Beijing) Biotechnology Co., Ltd. Male C57BL/6J mice, also weighing 18-22 g were obtained from Shanghai Lingchang Biotechnology Co., Ltd. All animals were provided with unrestricted access to both water and food. The animal experiments were conducted in accordance with ethical guidelines and received approval from the Institutional Animal Care and Use Committee (IACUC) of School of Life Sciences, Fudan University (Approval No. 2023JS020).

### In vivo expression of luciferase mRNA loaded in LNPs

FLuc mRNA-encapsulated LNPs were prepared and examined for particle size and encapsulation efficiency. BALB/c mice were intravenously or intramuscularly injected of LNPs encapsulating FLuc mRNA at a dose of 5 μg mRNA. Then, at 4 h, 24 h, and 48 h post-administration, the bioluminescence imaging procedure was performed using an IVIS Spectrum imaging system (PerkinElmer). Prior to each imaging session, mice received an intraperitoneal injection of 100 μL of D-luciferin (30 mg mL^-1^, dissolved in PBS) 8 min before the imaging procedure. For the repeated administration of FLuc mRNA, imaging was performed 4 h and 72 h following each administration.

### Luminex analysis for the detection of cytokines in serum

Cytokine levels in mouse serum were quantified utilizing the Mouse Premixed Multi-Analyte Kit (R&D Systems, cKlKmbpF) at 6 h post-administration. The cytokines assessed included IL-6, TNF-α, IFN-γ, MCP-1, IL-1β and IL-2. Serum samples were diluted in a 2-fold manner prior to measurement. All procedures were conducted in accordance with the manufacturer’s recommended protocol. Plates were read on a Luminex^®^200™ instrument.

### ELISA for VZV gE-specific IgG

VZV gE-specific antibody responses in immunized mice were determined by ELISA assay. In brief, 96-well plates were coated with 100 ng of recombinant VZV gE antigen (ACRO Biosystems, GLE-V52H3) dissolved in 50 μL of coating buffer per well and incubated at 4 overnight. Following this, the antigen-coated plates were blocked with a solution of 2% bovine serum albumin in PBS containing 0.1% Tween 20 (PBST) at room temperature for 1 h. Serum samples were initially diluted 250-fold and subsequently subjected to two-fold serial dilutions, resulting in a total of 11 gradients in PBS. After washing the plates with PBST, the serially diluted sera were added and incubated at room temperature for 2 h. To determine the gE-specific IgG antibody levels, the plates were incubated with HRP-conjugated goat anti-mouse IgG (Proteintech, SA00001-1) at 37 CC for 1 h and then the substrate tetramethylbenzidine (TMB) solution (Invitrogen, 002023) was then applied for color development. The color reaction was terminated by the addition of 1N sulfuric acid for about 10 min, and the optical density was measured at 450 nm using a microplate reader (Varioskan LUX, Thermo Fisher).

### ICS assays

An ICS assay was performed to assess gE-specific CD4^+^ and CD8^+^ T cell responses in the spleen of vaccinated mice. Spleen cells were stimulated with a peptide pool of gE (GenScript) at a concentration of 2 μg mL^-1^ in the presence of anti-mouse CD28 antibody (BD Bioscience, 553295) for 1 h. Then, a protein transport inhibitor containing monensin (BD Bioscience, 554724) was added into the culture, and the incubation continued for an additional 5 hours. Following stimulation, the cells were washed and stained with the Fixable Viability Stain 510 (BD Bioscience, 564406). The cells were then blocked with anti-mouse CD16/CD32 (BD Bioscience, 553140) and labeled with a surface antibody mixture including anti-mouse CD3-FITC, anti-mouse CD4-APC and anti-mouse CD8-PerCP-Cy5.5 (BD Bioscience, 555274, 553051 and 551162, respectively) for 30 min at 4. After surface staining, the cells were fixed and permeabilized for 20 min at 4 using a Fixation/Permeabilization Kit (BD Bioscience, 554714). Subsequently, the cells were stained for intracellular cytokines using anti-mouse IFN-γ-PE-Cy7, anti-mouse TNF-α-BV650, anti-mouse IL-2-BV605, anti-mouse IL-4-BV711 and anti-mouse IL-5-PE (BD Bioscience, 557649, 563943, 563911, 564005 and 554395, respectively). The stained cells were then washed, resuspended in PBS, and analyzed using a flow cytometer (Cytek Aurora NL).

### IFN-γ and IL-2 ELISpot assays

ELISpot is a more intuitive method for quantifying antigen-specific immune cells that secret IFN-γ and IL-2. In this procedure, a total of one million spleen cells, stimulated with the peptide pool, were added to each well of the plates pre-coated with antibodies specific for either IFN-γ or IL-2 from the ELISpot kits (MABTECH, 3321-4AST-10 and 3441-4APW-10, respectively). The cells were cultured at 37 °C for 48 h. Following that, the plates were washed with PBS and subsequently incubated with biotinylated anti-IFN-γ, or anti-IL-2 antibody at room temperature for 2 h. This was followed by a one-hour incubation with streptavidin-ALP for 1 h at room temperature. The development of spots was initiated by the addition of a substrate solution, after which the plates were rinsed with water. Finally, the plates were allowed to air dry in a shaded environment and were analyzed using an ELISpot analyzer (AT-SPOT, SINSAGE).

### Determination of blood biochemical indicators

Mouse blood samples were collected at certain timepoints and subsequently centrifuged at 15,000 rpm for 5 min at 4 to obtain mouse sera. In the study of the VZV vaccine, ALT, AST and Crea in the serum collected two weeks post-booster were quantified using an automatic biochemical analyzer (Mindray, BS-450). In the investigation of PCSK9 editing, LDL-C and TC in the serum collected two weeks after administration were measured using an automatic biochemical analyzer (Mindray, BS-450).

### Sequence of sgPCSK9

5′-mG*mG*mC*UGAUGAGGCCGCACAUGGUUUUAGAGCUAGAAAUAGCA AGUUAAAAUAAGGCUAGUCCGUUAUCAACUUGAAAAAGUGGCACCGAGUCGG*mU*mG*mC (* = phosphorothioate, m = 2′-O-Me). (GenScript)

### NGS analysis

DNA was extracted from the liver on Day 14 following administration, utilizing a genomic DNA kit (TIANGEN, DP304). PCR primers were specifically designed to amplify the region surrounding the target site within the PCSK9 gene. Subsequently, an amplicon library was constructed through two rounds of PCR. The resulting amplicons were sequenced using an Illumina NovaSeq S4, and the sequencing data were analyzed for the types and frequencies of indels using a CRISPResso2 software.

PCR forward primer: 5′-ACACTCTTTCCCTACACGACGCTCTTCCGATCTNNNNGGACGAGGATGGAGATTATG

PCR reverse primer: 5′-ACTGGAGTTCAGACGTGTGCTCTTCCGATCTNNNNCATCACCCCAACCCCAAAG

### ELISA for detection of serum PCSK9

Blood samples were obtained from mice on Day 14 following the administration of Cas9 mRNA. The samples were then centrifuged at 15,000 rpm for 5 min at 4 to isolate the serum. The concentrations of PCSK9 protein in the serum were measured using a mouse PCSK9 ELISA kit (Sino Biological, KIT50251).

### Data analysis

The pKa values of the lipid compounds were predicted using ChemDraw Professional 15.0. NMR spectra of the lipid compounds were analyzed by MestReNova 5.3. All data were processed and visualized using GraphPad Prism 8. The results are presented as the mean ± SD, and statistical significance was assessed using one-way ANOVA (ns or unmarked indicates no significance; *, p < 0.05; **, p < 0.01; ***, p < 0.001; ****, p < 0.0001).

## Acknowledgements

This research was financially supported by the National Key Research and Development Program of China (No. 2023YFC2306404 and No. 2023YFC2604303), the National Natural Science Foundation of China (No. 32301174), and the China Postdoctoral Science Foundation (No. 2024M750497). Additionally, this work received support from the Innovation Program of Shanghai Municipal Education Commission (No. 2021-01-07-00-07-E00074) and the Zhangjiang mRNA Innovation and Translation Center at Shanghai. The authors express their gratitude to the R&D Department of RNACure Biopharma for their assistance in the preparation and evaluation of LNPs.

## Author Contributions

J.Lin, K.C. and X.L. conceived the study and designed the experiments. K.C. designed and synthesized the HL compounds. X.L. and K.C. designed and executed the in vitro experiments and the repeated administration study. K.C. designed and participated in the study on the physicochemical properties of LNPs. X.L. conducted the vaccine against VZV. S.F. performed the gene editing of PCSK9. Y.Li carried out the molecular dynamic simulations. X.L. and N.G. conducted the in vitro experiments. K.C., X.L., T.J., Y.Liu and X.Z. participated in the preparation, characterization, and in vivo evaluation of LNPs. H.Y. provided support for the NMR study. K.C. and X.L. analyzed the data. K.C. drafted the manuscript. J.Lin and M.Q. revised the manuscript. J.Lin and J.Lu supervised the study. The final manuscript was reviewed and approved by all authors.

