## Supplementary material for "Hydroxychloroquine-functionalized Ionizable Lipids Mitigate Inflammatory Responses in mRNA Therapeutics": Supplementary Information.docx (from another version of this preprint: version 1, 2025-03-07)

**Supplementary information of lipid synthesis**

**Synthesis of HL-1**

A mixture of 2-decyltetradecan-1-ol (**h1**, 1.7732 g, 5 mmol), succinic anhydride (**h2**, 0.7005 g, 7 mmol) and 4-dimethylaminopyridine (DMAP) (0.1222 g, 1 mmol) in anhydrous dichloromethane (DCM) (30 mL) was stirred at room temperature overnight. The solvent was removed in vacuo after reaction and the residue was then purified by silica gel chromatography (petroleum ether (PE): ethyl acetate (EA)= 3:1) to afford **Compound 1** (2.0462 g) in 90% yield as white semisolid.

A mixture of **Compound 1** (0.9094 g, 2 mmol), hydroxychloroquine (**HCQ**) (1.0076 g, 3 mmol), 1,3-dicyclohexylcarbodiimide (DCC) (0.6170 g, 3 mmol) and DMAP (61 mg, 0.5 mmol) in anhydrous DCM (30 mL) was stirred at room temperature for 6 h. The reaction solution was filtered to remove suspended solids and then purified by silica gel chromatography (DCM: methanol (MeOH) = 70:1) to afford **HL-1** (0.8498 g) in 55% yield as yellow oil. The synthesis route and characterization of the final product were listed below (Supplementary Fig. 1, 7).

**Synthesis of HL-2**

A mixture of heptadecan-9-ol (**h3**, 2.5647 g, 10 mmol), **h2** (1.5010 g, 15 mmol) and DMAP (0.2443 g, 2 mmol) in anhydrous DCM (50 mL) was stirred at room temperature overnight. The solvent was removed in vacuo after reaction and the residue was then purified by silica gel chromatography (PE: EA = 3:1) to afford **Compound 2** (3.2802 g) in 92% yield as white solid.

A mixture of **Compound 2** (0.7131 g, 2 mmol), **HCQ** (1.0076 g, 3 mmol), DCC (0.6170 g, 3 mmol) and DMAP (61 mg, 0.5 mmol) in anhydrous DCM (30 mL) was stirred at room temperature for 6 h. The reaction solution was filtered to remove suspended solids and then purified by silica gel chromatography (DCM: MeOH = 60:1) to afford **HL-2** (0.7149 g) in 53% yield as yellow oil. The synthesis route and characterization of the final product were listed below (Supplementary Fig. 2, 8).

**Synthesis of HL-4**

A mixture of 2-heptylundecanoic acid (**h4**, 0.5690 g, 2 mmol), **HCQ** (1.0076 g, 3 mmol), DCC (0.6170 g, 3 mmol) and DMAP (61 mg, 0.5 mmol) in anhydrous DCM (30 mL) was stirred at room temperature for 6 h. The reaction solution was filtered to remove suspended solids and then purified by silica gel chromatography (DCM: MeOH = 60:1) to afford **HL-4** (0.5782 g) in 48% yield as white semisolid. The synthesis route and characterization of the final product were listed below (Supplementary Fig. 4, 11).

**Synthesis of TL-4 and HL-3**

To a mixture of **h3** (2.5647 g, 10 mmol) and triethylamine (TEA) (1.5178 g, 15 mmol) in anhydrous DCM (30 mL) cooled in ice, acryloyl chloride (**h5**, 1.3576 g, 15 mmol) dissolved in anhydrous DCM (20 mL) was added dropwise. The mixture was then stirred in an ice-water bath for 4 h. The reaction solution was filtered to remove suspended solids and then purified by silica gel chromatography (PE: DCM = 3:1) to afford **Compound 3** (2.7410 g) in 88% yield as colorless liquid.

A mixture of **Compound 3** (1.8361 g, 6 mmol) and ethanolamine (**h6**, 0.1832 g, 3 mmol) in propan-2-ol (Me_2_CHOH, 30 mL) was stirred at 80 ℃ overnight. The solvent was removed in vacuo after reaction and the residue was then purified by silica gel chromatography (PE: EA = 6:1) to afford **TL-4** (1.6827 g) in 82% yield as colorless oil.

A mixture of **TL-4** (1.3643 g, 2 mmol)), **h2** (0.6109 g, 4 mmol) and DMAP (61 mg, 0.5 mmol) in anhydrous DCM (30 mL) was stirred at room temperature overnight. The solvent was removed in vacuo after reaction and the residue then was purified by silica gel chromatography (PE: EA = 1:1) to afford **Compound 4** (1.4428 g) in 92% yield as white solid.

A mixture of **Compound 4** (0.7841 g, 1 mmol), **HCQ** (0.6718 g, 2 mmol), DCC (0.4127 g, 2 mmol) and DMAP (61 mg, 0.5 mmol) in anhydrous DCM (30 mL) was stirred at room temperature for 6 h. The reaction solution was filtered to remove suspended solids and then purified by silica gel chromatography (DCM: MeOH = 70:1) to afford **HL-3** (0.5401 g) in 49% yield as white semisolid. The synthesis route and characterization of the final products were listed below (Supplementary Fig. 3, 9 and 10).

**Synthesis of HL-5**

A mixture of **SM-102** (1.4204 g, 2 mmol)), **h2** (0.6109 g, 4 mmol) and DMAP (61 mg, 0.5 mmol) in anhydrous DCM (30 mL) was stirred at room temperature overnight. The solvent was removed in vacuo after reaction and the residue was then purified by silica gel chromatography (DCM: MeOH = 30:1) to afford **Compound 5** (1.5087 g) in 93% yield as white oil.

A mixture of **Compound 5** (0.8102 g, 1 mmol), **HCQ** (0.6718 g, 2 mmol), DCC (0.4127 g, 2 mmol) and DMAP (61 mg, 0.5 mmol) in anhydrous DCM (30 mL) was stirred at room temperature for 6 h. The reaction solution was filtered to remove suspended solids and then purified by silica gel chromatography (DCM: MeOH = 50:1) to afford **HL-5** (0.5189 g) in 46% yield as yellow oil. The synthesis route and characterization of the final products were listed below (Supplementary Fig. 3, 12 and 13).

**Synthesis of HL-6 and CL-17**

To a mixture of 2-hydroxymethyl-1,3-propanediol (**h7**, 0.5306 g, 5 mmol), DCC (3.0949 g, 15 mmol) and DMAP (0.1222 g, 1 mmol) in a mixed solvent of DCM and N,N-dimethylformamide (DMF) (30 mL, DCM: DMF = 2:1), **Compound 2** (3.5654 g, 10 mmol) dissolved in anhydrous DCM (20 mL) was added dropwise. The mixture was then stirred at room temperature overnight. The reaction solution was filtered to remove suspended solids and the mixed solvent was removed in vacuo. Then, the residue was purified by silica gel chromatography (PE: EA = 7:1) to afford **Compound 6** (2.0755 g) in 53% yield as colorless oil.

A mixture of **Compound 6** (1.5664 g, 2 mmol)), **h2** (0.6109 g, 4 mmol) and DMAP (61 mg, 0.5 mmol) in anhydrous DCM (30 mL) was stirred at room temperature overnight. The solvent was removed in vacuo after reaction and the residue was then purified by silica gel chromatography (PE: EA = 5:1) to afford **Compound 7** (1.5015 g) in 85% yield as white semisolid.

A mixture of **Compound 7** (0.8833 g, 1 mmol), **HCQ** (0.6718 g, 2 mmol), DCC (0.4127 g, 2 mmol) and DMAP (61 mg, 0.5 mmol) in anhydrous DCM (30 mL) was stirred at room temperature for 6 h. The reaction solution was filtered to remove suspended solids and then purified by silica gel chromatography (DCM: MeOH = 70:1) to afford **HL-6** (0.5405 g) in 45% yield as white semisolid.

For the synthesis of CL-17, a mixture of **Compound 7** (0.8833 g, 1 mmol), 2-diethylaminoethanol (**h8**, 0.2344 g, 2 mmol), DCC (0.4127 g, 2 mmol) and DMAP (61 mg, 0.5 mmol) in anhydrous DCM (30 mL) was stirred at room temperature for 6 h. The reaction solution was filtered to remove suspended solids and then purified by silica gel chromatography (DCM: MeOH = 80:1) to afford **CL-17** (0.4617 g) in 47% yield as white semisolid. The synthesis route and characterization of the final products were listed below (Supplementary Fig. 3, 14 and 15).

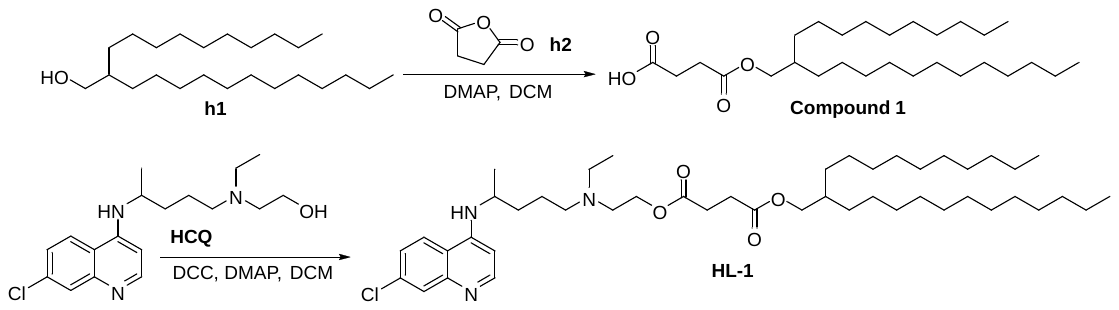

**Figure S1 Synthesis route of HL-1.**

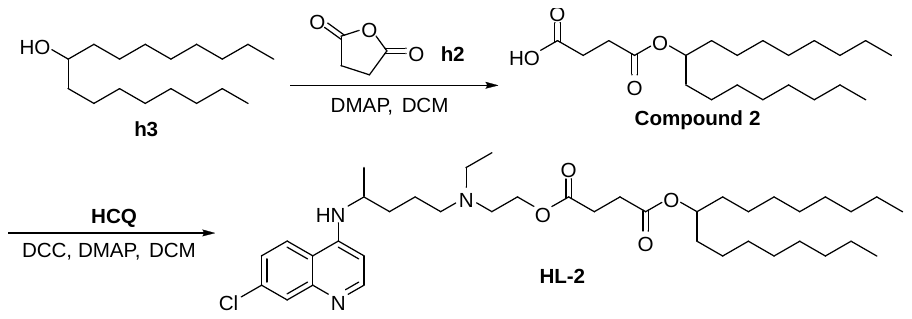

**Supplementary Fig. 2 Synthesis route of HL-2.**

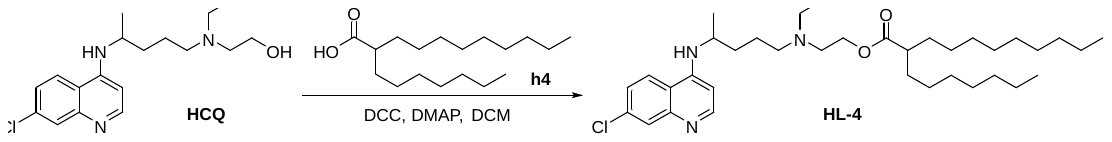

**Supplementary Fig. 3 Synthesis route of HL-4.**

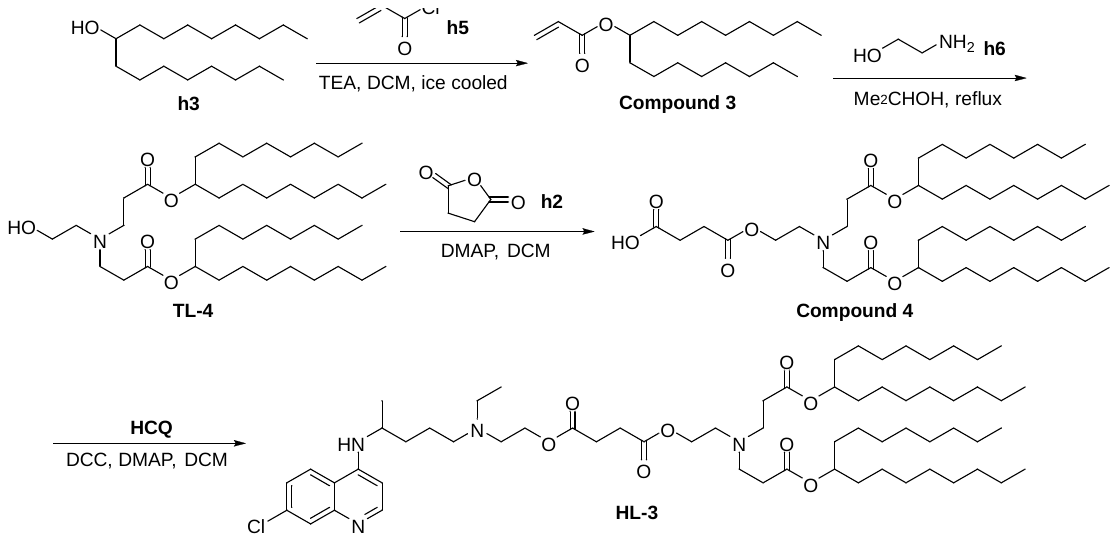

**Supplementary Fig. 4 Synthesis route of TL-4 and HL-3.**

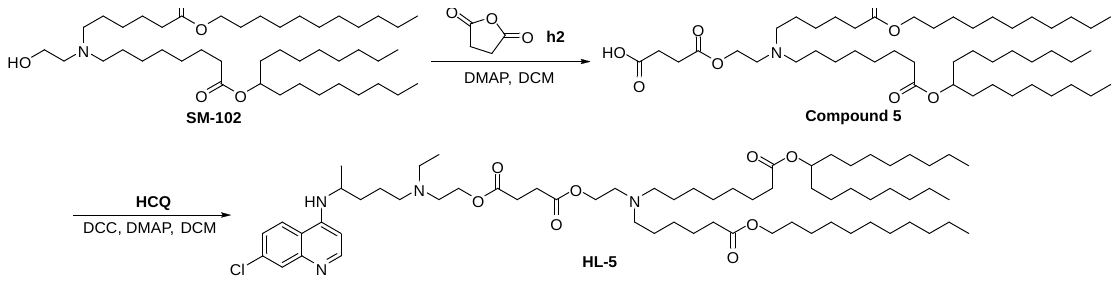

**Supplementary Fig. 5 Synthesis route of HL-5.**

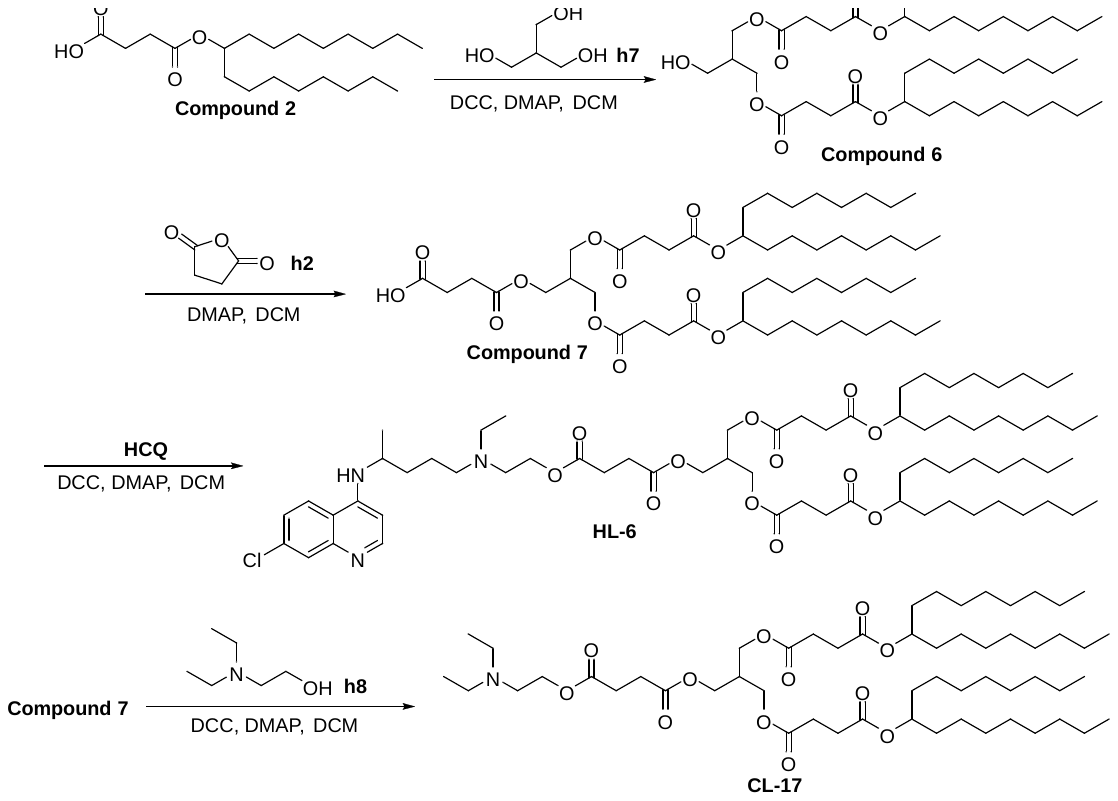

**Supplementary Fig. 6 Synthesis route of HL-6 and CL-17.**

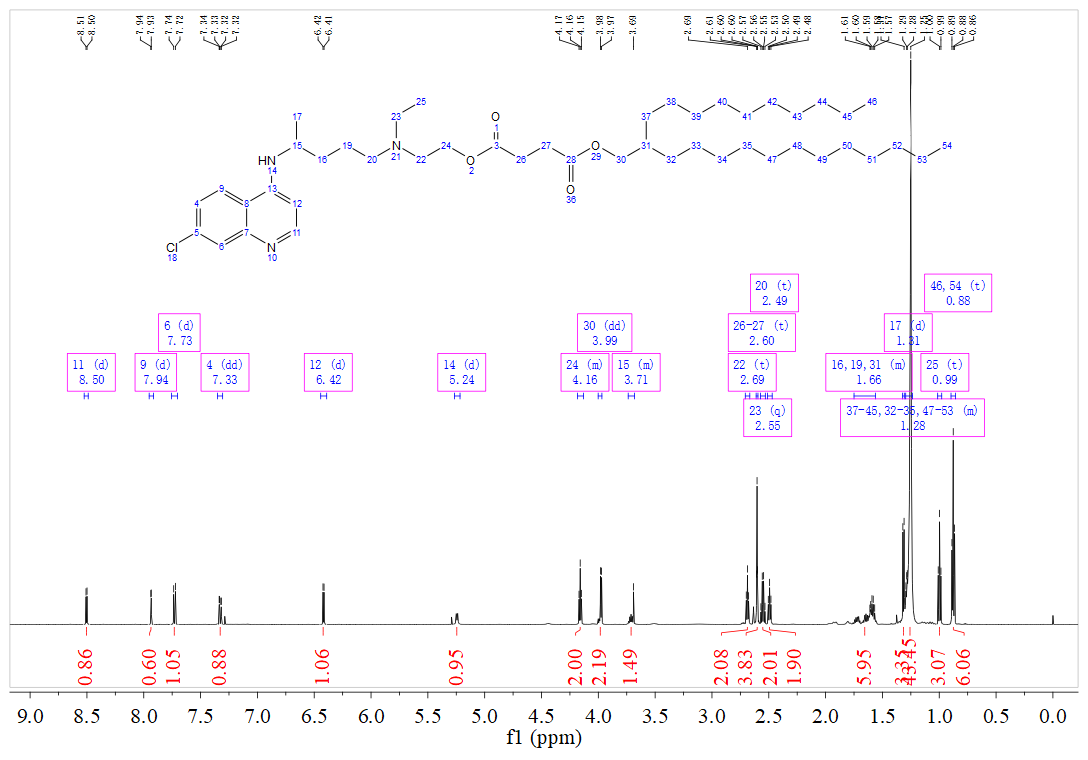

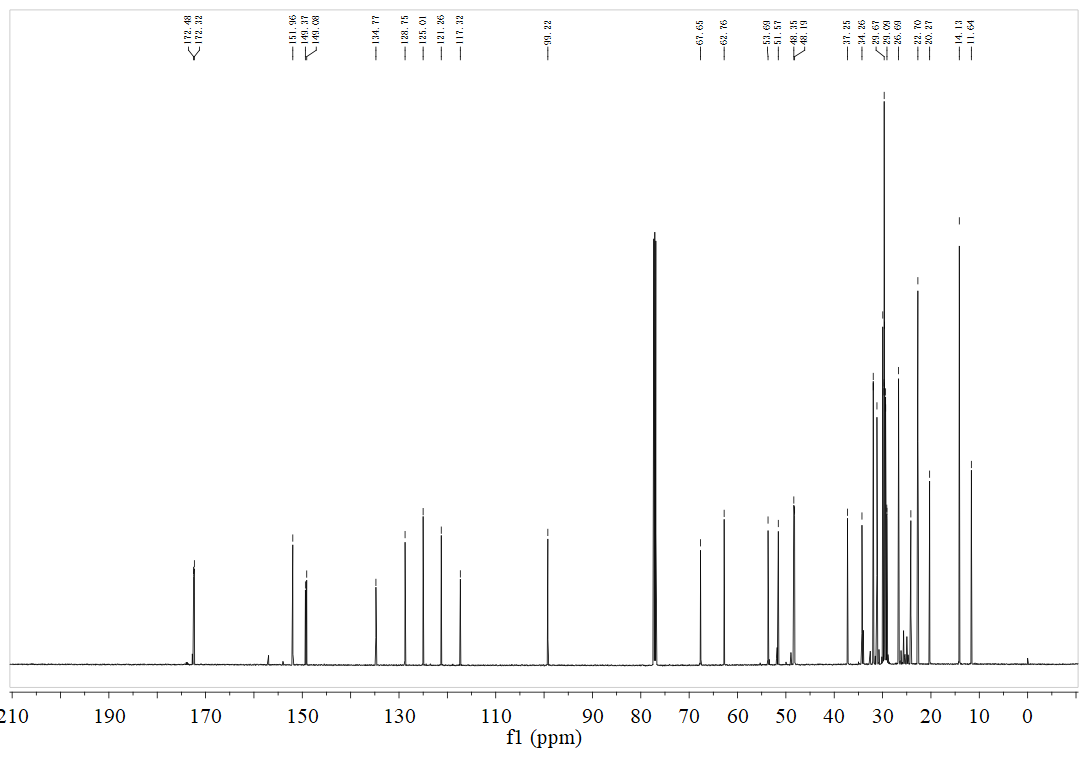

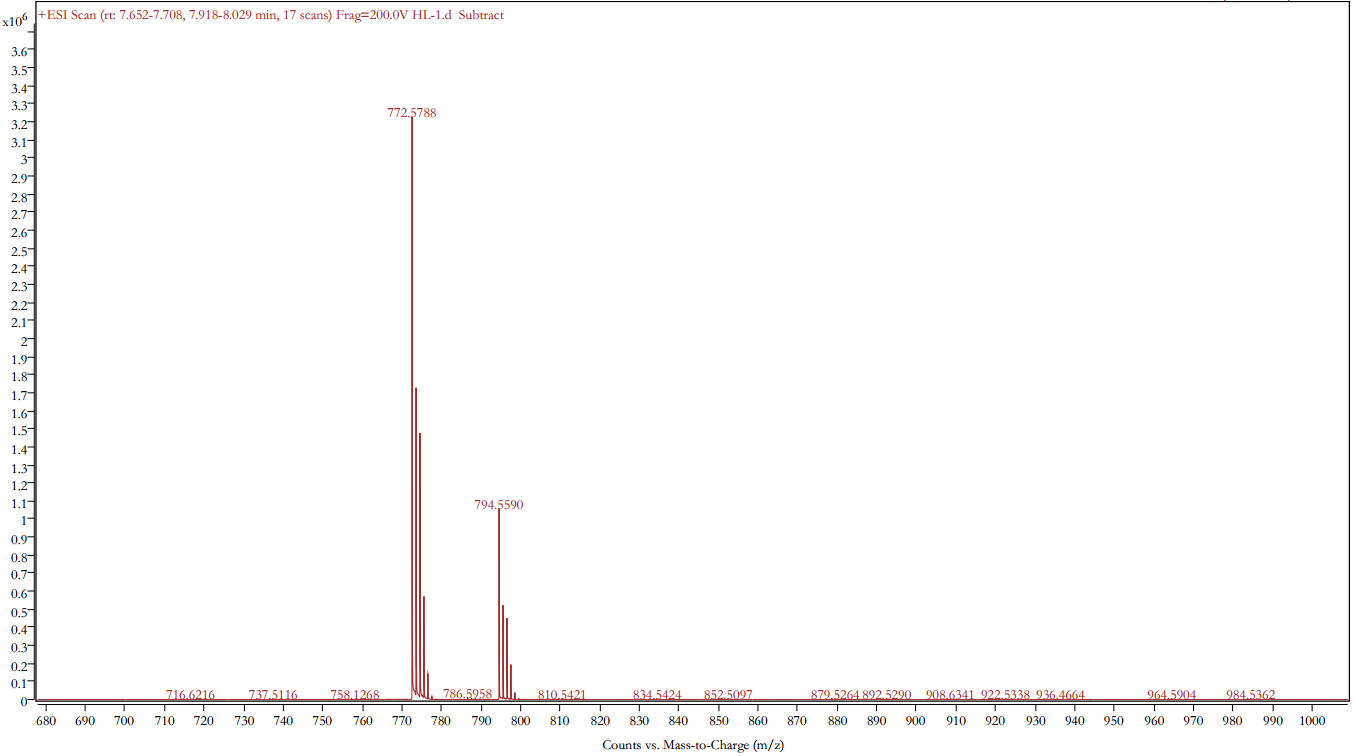

**Supplementary Fig. 7 Characterization of HL-1. ^1^H NMR** (600 MHz, CDCl_3_) δ 8.50 (d, *J* = 5.4 Hz, 1H), 7.94 (d, *J* = 2.2 Hz, 1H), 7.73 (d, *J* = 9.0 Hz, 1H), 7.33 (dd, *J* = 8.9, 2.2 Hz, 1H), 6.42 (d, *J* = 5.5 Hz, 1H), 5.24 (d, *J* = 7.4 Hz, 1H), 4.18 – 4.13 (m, 2H), 3.99 (dd, *J* = 11.5, 5.8 Hz, 2H), 3.74 – 3.68 (m, 1H), 2.69 (t, *J* = 6.2 Hz, 2H), 2.60 (t, *J* = 1.7 Hz, 4H), 2.55 (q, *J* = 7.1 Hz, 2H), 2.49 (t, *J* = 6.9 Hz, 2H), 1.75 – 1.56 (m, 6H), 1.31 (d, *J* = 6.4 Hz, 3H), 1.30 – 1.24 (m, 43H), 0.99 (t, 3H), 0.88 (t, 6H). **^13^C NMR** (151 MHz, CDCl_3_) δ 172.48, 172.32, 151.96, 149.37, 149.08, 134.77, 128.75, 125.01, 121.26, 117.32, 99.22, 67.65, 62.76, 53.69, 51.57, 48.35, 48.19, 37.25, 34.26, 31.93, 31.93, 31.16, 29.96, 29.70, 29.69, 29.67, 29.65, 29.62, 29.37, 29.36, 29.13, 29.09, 26.69, 24.15, 22.70, 20.27, 14.13, 11.64. **MS** (ESI, m/z) [M+H]^+^ calculated for C_46_H_78_ClN_3_O_4_, 772.57; Found, 772.5788.

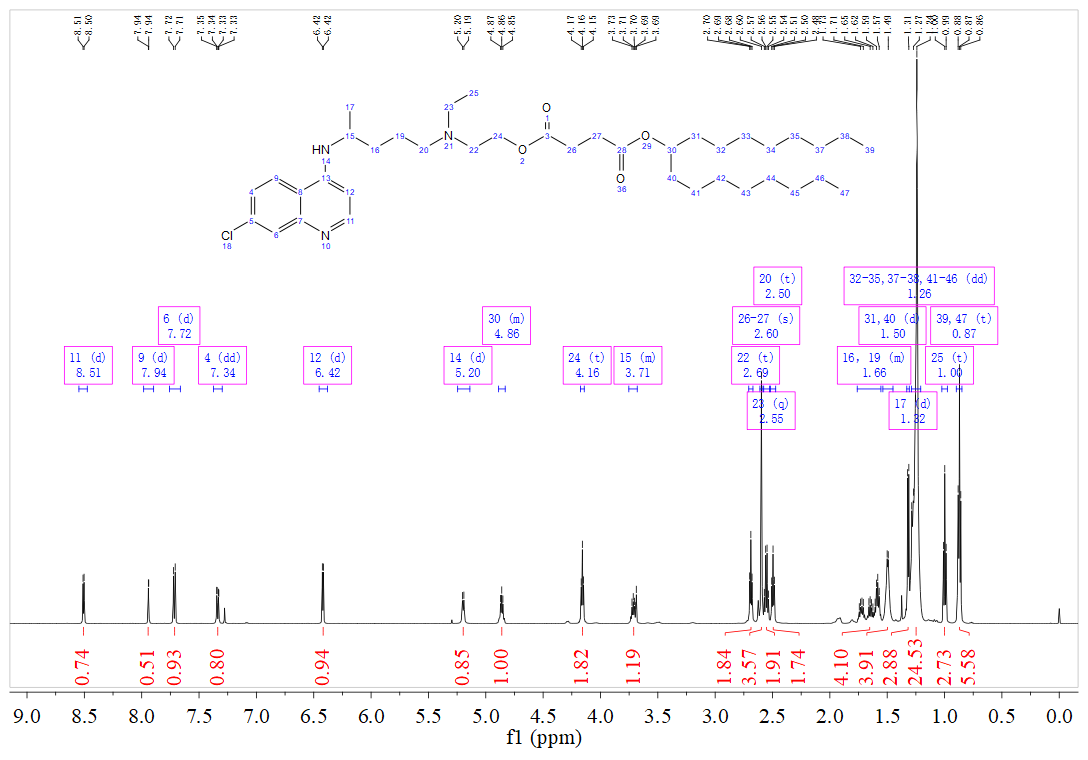

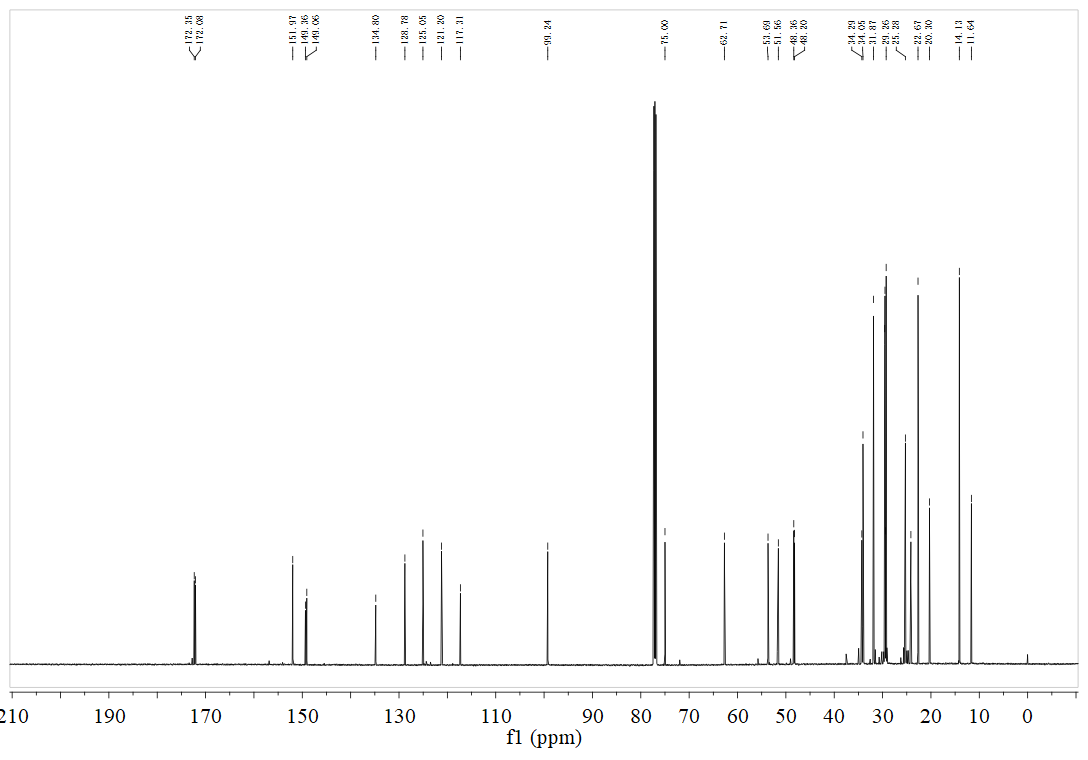

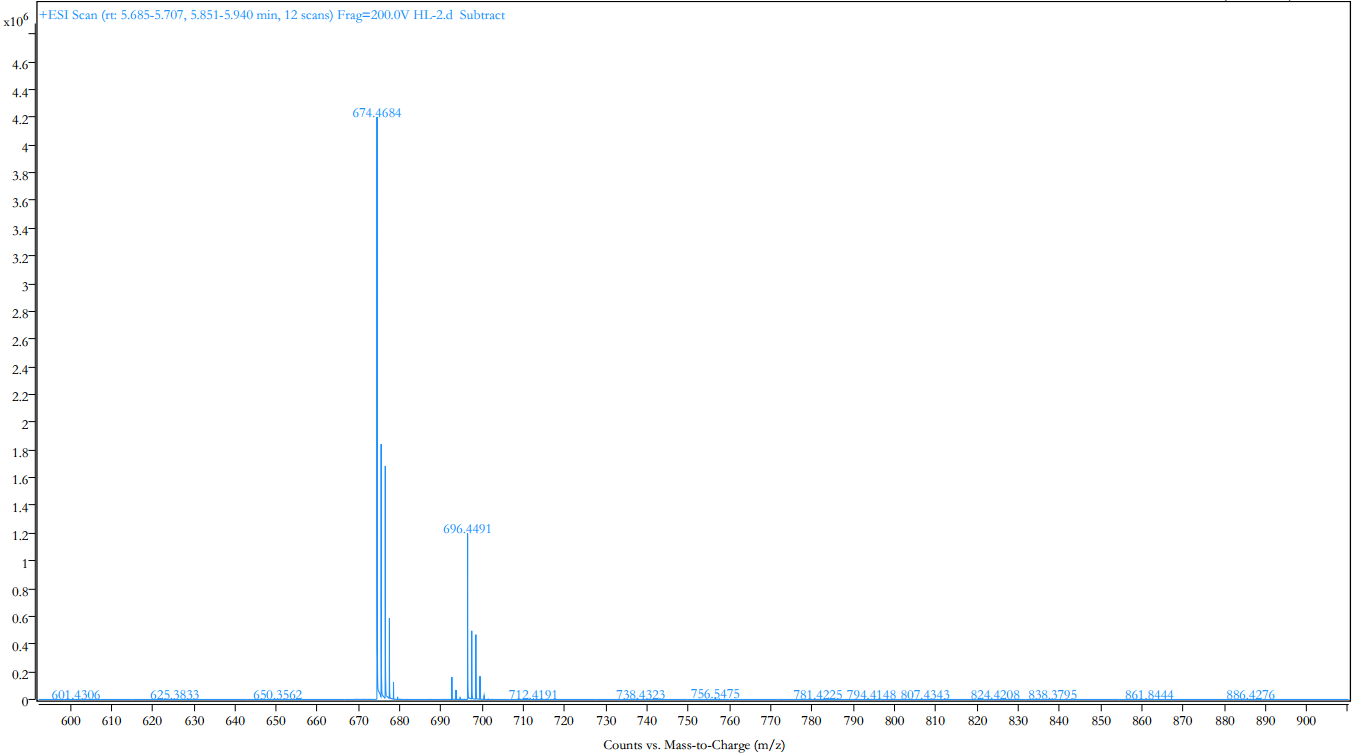

**Supplementary Fig. 8 Characterization of HL-2. ^1^H NMR** (600 MHz, CDCl_3_) δ 8.51 (d, *J* = 5.4 Hz, 1H), 7.94 (d, *J* = 2.0 Hz, 1H), 7.72 (d, *J* = 9.0 Hz, 1H), 7.34 (dd, *J* = 8.9, 2.0 Hz, 1H), 6.42 (d, *J* = 5.4 Hz, 1H), 5.20 (d, *J* = 7.3 Hz, 1H), 4.89 – 4.83 (m, 1H), 4.16 (t, *J* = 6.1 Hz, 2H), 3.75 – 3.68 (m, 1H), 2.69 (t, *J* = 6.2 Hz, 2H), 2.60 (s, 4H), 2.55 (q, *J* = 7.1 Hz, 2H), 2.50 (t, *J* = 6.9 Hz, 2H), 1.76 – 1.55 (m, 4H), 1.50 (d, *J* = 5.8 Hz, 4H), 1.32 (d, *J* = 6.3 Hz, 3H), 1.26 (dd, *J* = 18.7, 8.1 Hz, 25H), 1.00 (t, *J* = 7.1 Hz, 3H), 0.87 (t, *J* = 7.0 Hz, 6H). **^13^C NMR** (151 MHz, CDCl_3_) δ 172.35, 172.08, 151.97, 149.36, 149.06, 134.80, 128.78, 125.05, 121.20, 117.31, 99.24, 75.00, 62.71, 53.69, 51.56, 48.36, 48.20, 34.29, 34.05, 31.87, 29.55, 29.50, 29.37, 29.26, 25.28, 24.14, 22.67, 20.30, 14.13, 11.64. **MS** (ESI, m/z) [M+H]^+^ calculated for C_39_H_64_ClN_3_O_4_, 674.46; Found, 674.4684.

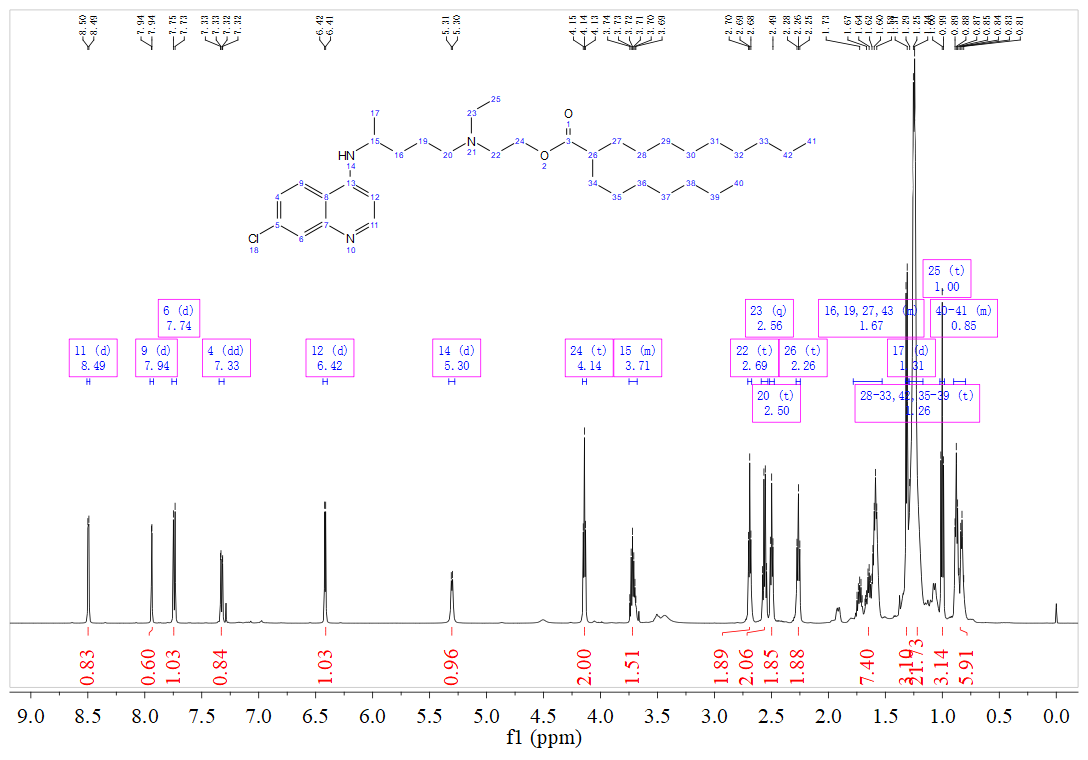

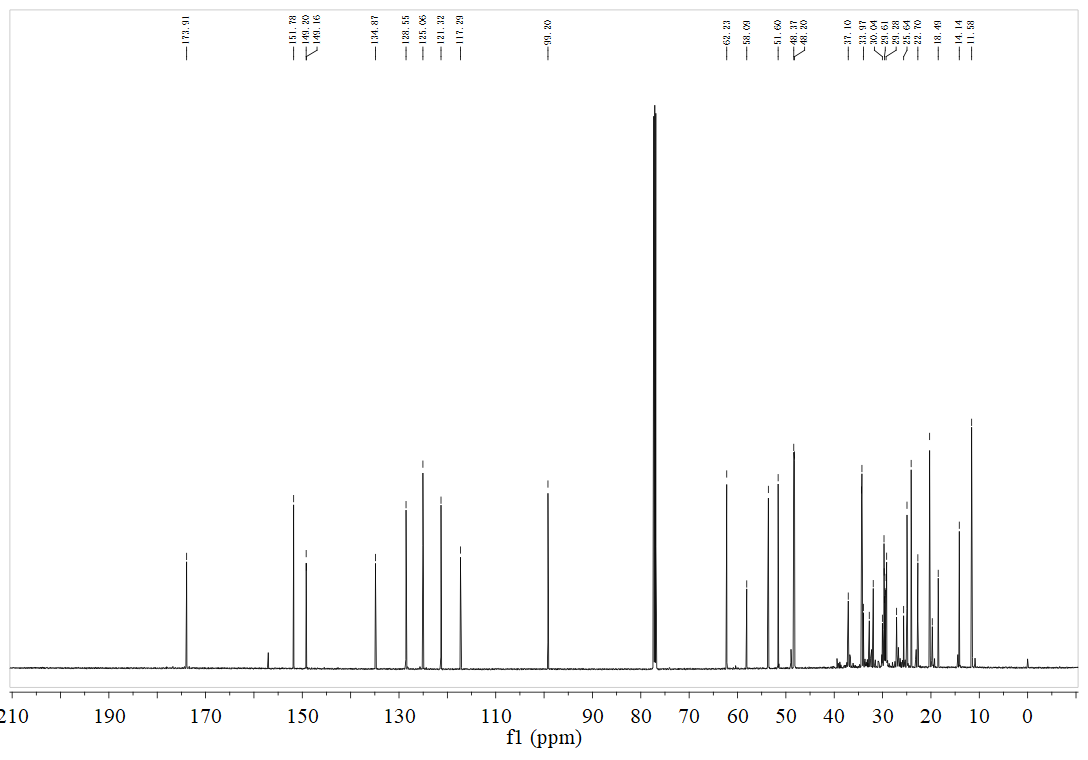

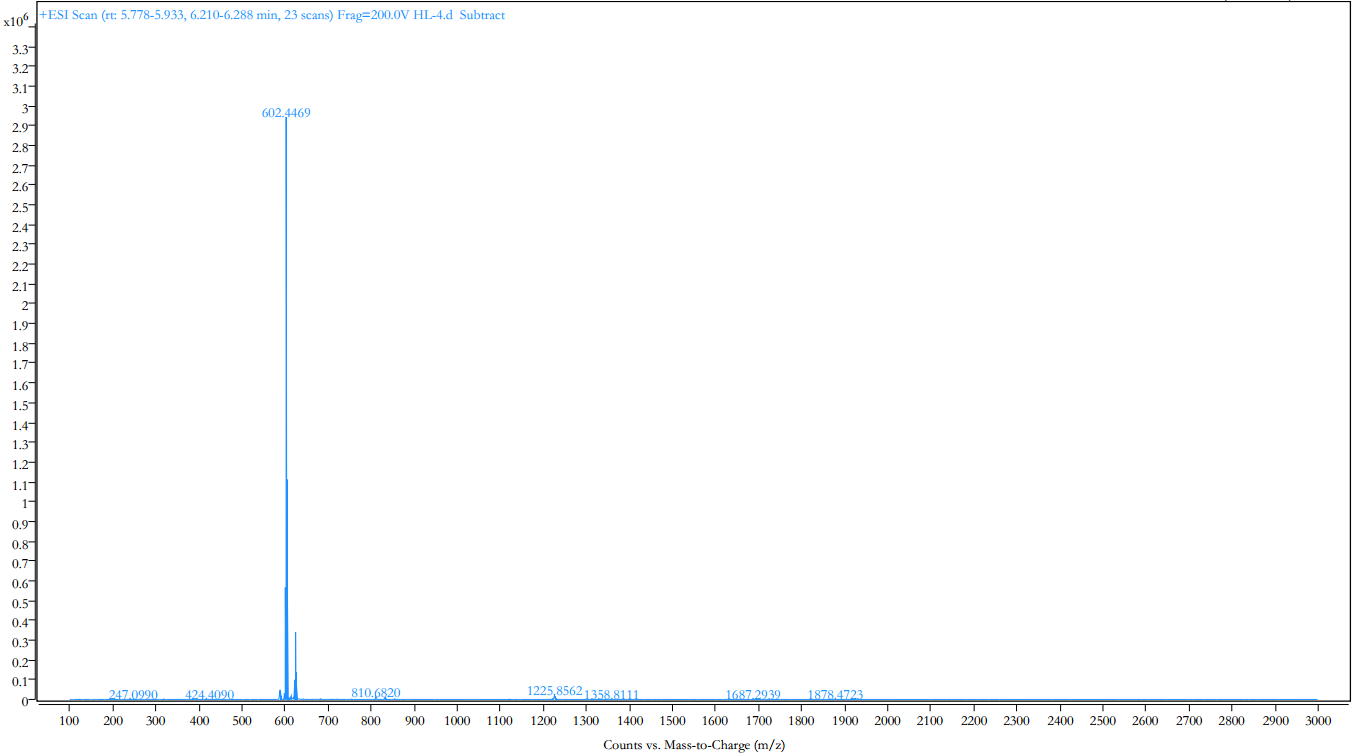

**Supplementary Fig. 9 Characterization of HL-4. ^1^H NMR** (600 MHz, CDCl_3_) δ 8.49 (d, *J* = 5.4 Hz, 1H), 7.94 (d, *J* = 2.0 Hz, 1H), 7.74 (d, *J* = 9.0 Hz, 1H), 7.33 (dd, *J* = 8.9, 2.0 Hz, 1H), 6.42 (d, *J* = 5.5 Hz, 1H), 5.30 (d, *J* = 6.9 Hz, 1H), 4.14 (t, *J* = 6.2 Hz, 2H), 3.75 – 3.68 (m, 2H), 2.69 (t, *J* = 6.2 Hz, 2H), 2.56 (q, *J* = 7.1 Hz, 2H), 2.50 (t, *J* = 6.8 Hz, 2H), 2.26 (t, *J* = 7.5 Hz, 2H), 1.78 – 1.53 (m, 7H), 1.31 (d, *J* = 6.3 Hz, 3H), 1.26 (t, *J* = 13.0 Hz, 22H), 1.00 (t, *J* = 7.1 Hz, 3H), 0.90 – 0.80 (m, 6H). **^13^C NMR** (151 MHz, CDCl_3_) δ 173.91, 151.78, 149.20, 149.16, 134.87, 128.55, 125.06, 121.32, 117.29, 99.20, 62.23, 58.09, 53.62, 51.60, 48.37, 48.20, 37.10, 34.34, 34.27, 33.97, 32.75, 31.93, 30.04, 29.71, 29.66, 29.61, 29.47, 29.37, 29.28, 29.15, 27.10, 25.64, 24.95, 24.09, 22.70, 20.27, 19.72, 18.49, 14.14, 11.58. **MS** (ESI, m/z) [M+H]^+^ calculated for C_36_H_60_ClN_3_O_2_, 602.44; Found, 602.4469.

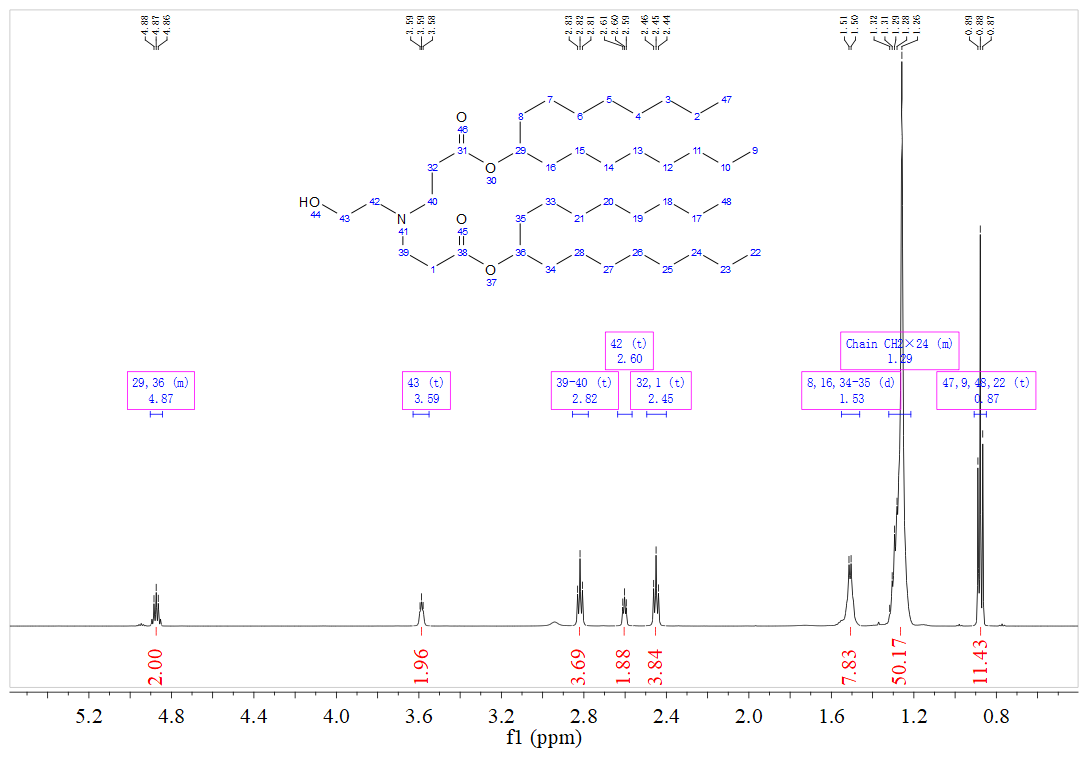

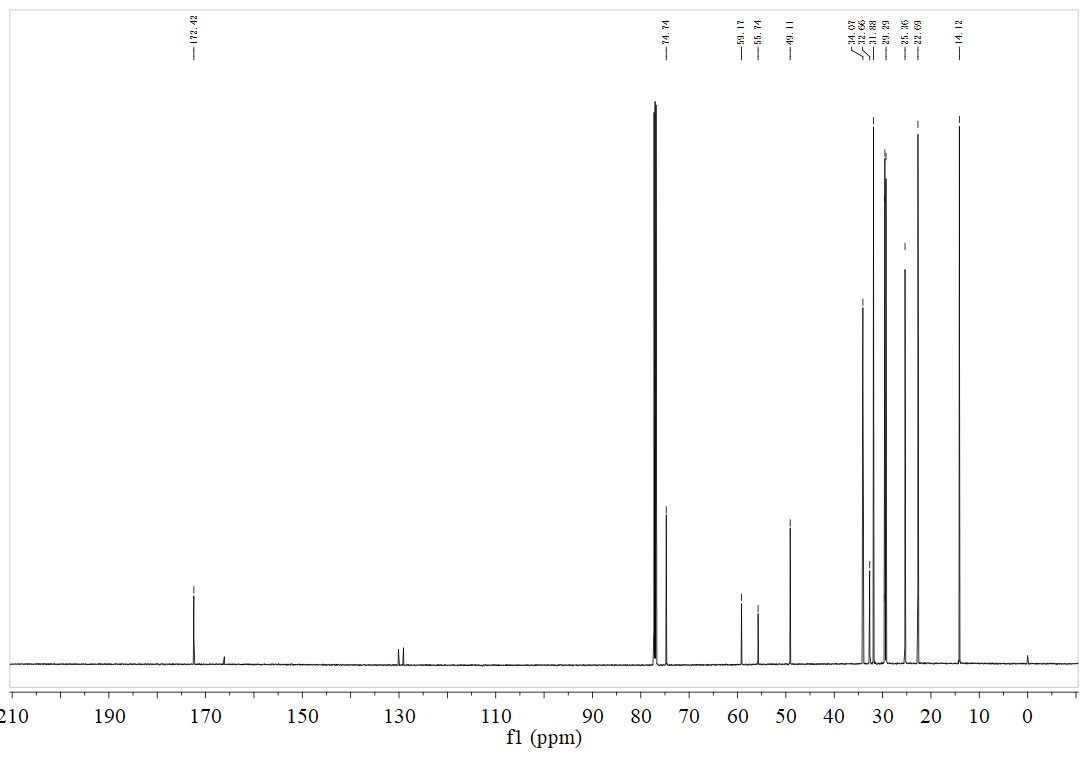

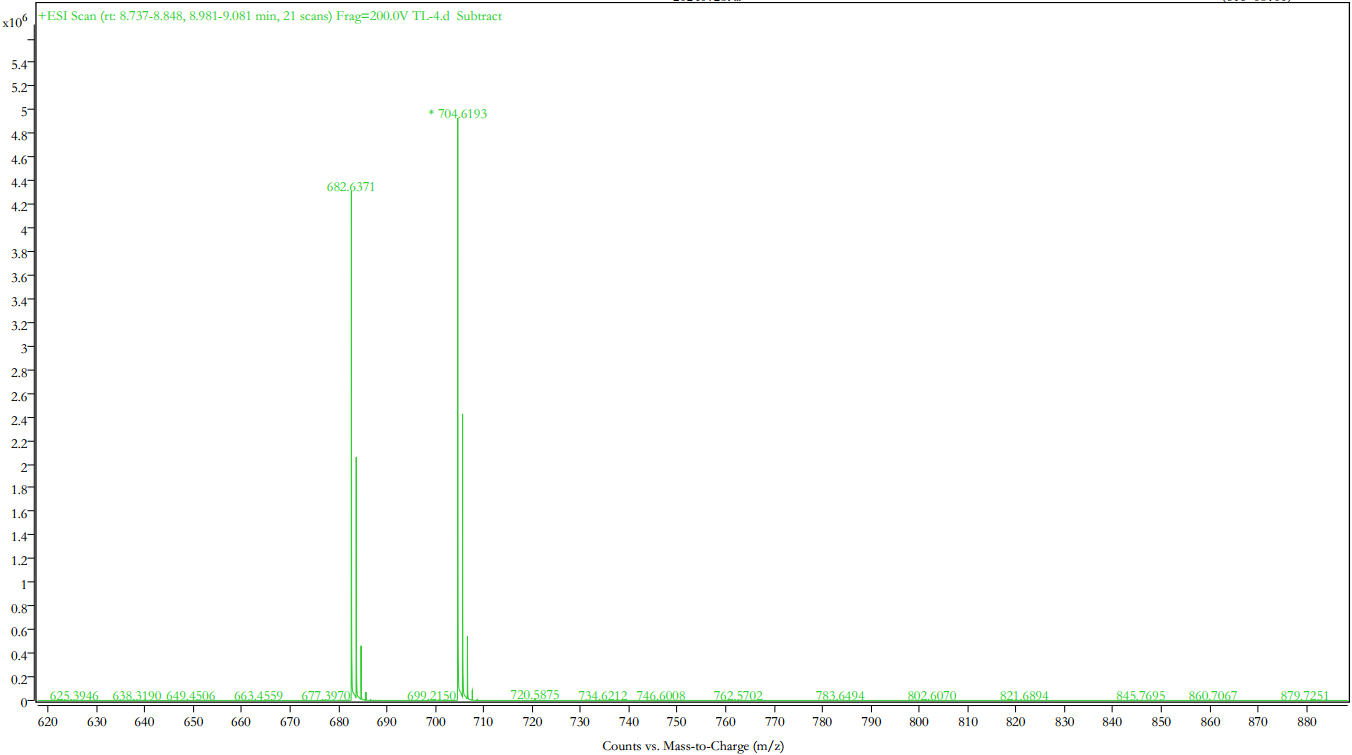

**Supplementary Fig. 10 Characterization of TL-4. ^1^H NMR** (600 MHz, CDCl_3_) δ 4.90 – 4.84 (m, 2H), 3.59 (t, *J* = 4.5 Hz, 2H), 2.82 (t, *J* = 7.0 Hz, 4H), 2.60 (t, *J* = 5.0 Hz, 2H), 2.45 (t, *J* = 7.0 Hz, 4H), 1.53 (d, *J* = 20.5, 4.8 Hz, 8H), 1.32 – 1.21 (m, 50H), 0.87 (t, 11H). **^13^C NMR** (151 MHz, CDCl_3_) δ 172.42, 74.74, 59.17, 55.74, 49.11, 34.07, 32.66, 31.88, 29.57, 29.52, 29.50, 29.29, 25.36, 22.69, 14.12. **MS** (ESI, m/z) [M+H]^+^ calculated for C_42_H_83_NO_5_, 682.63; Found, 682.6371.

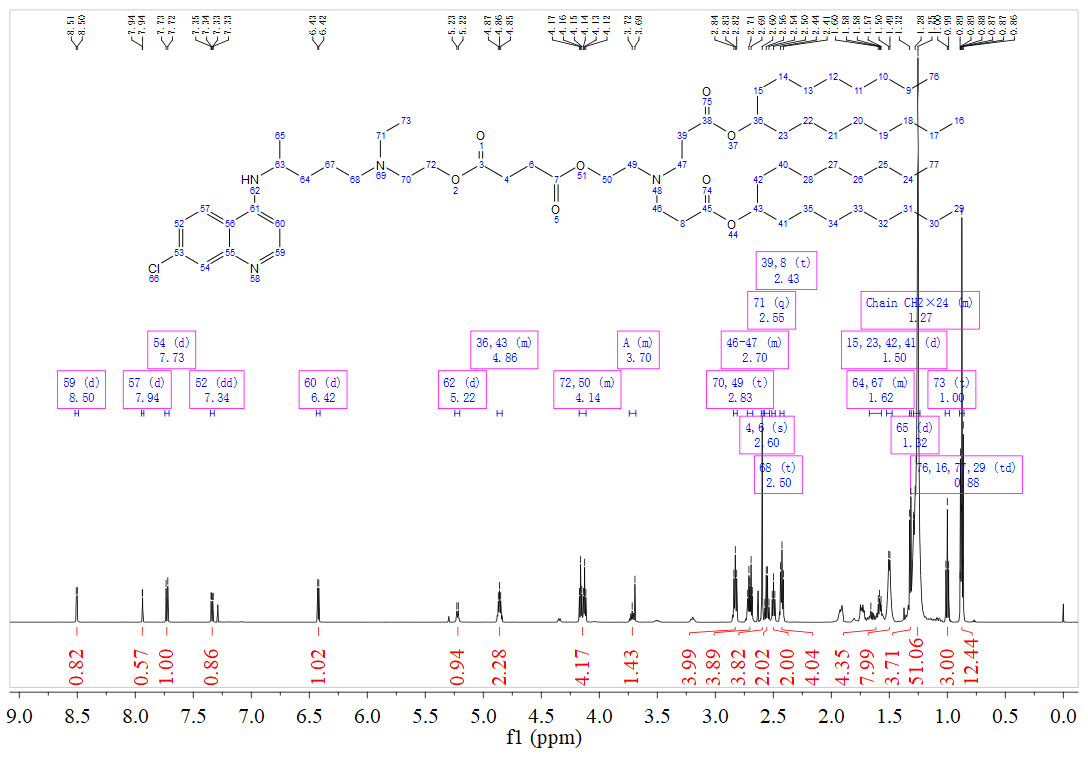

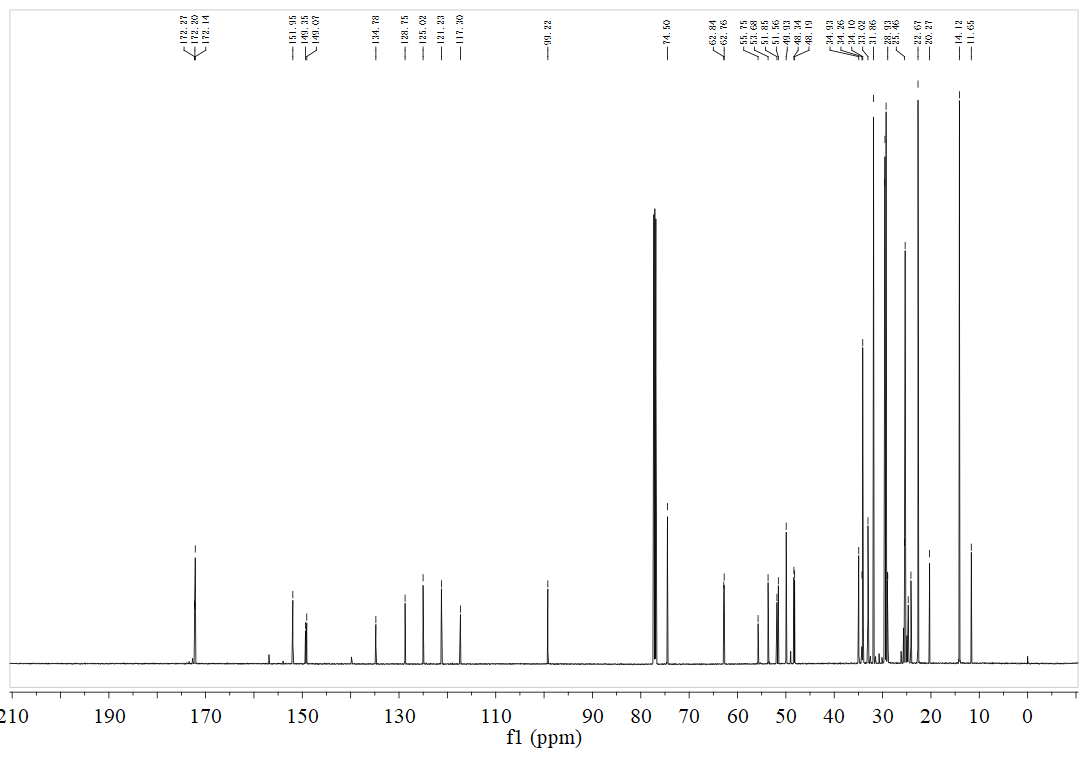

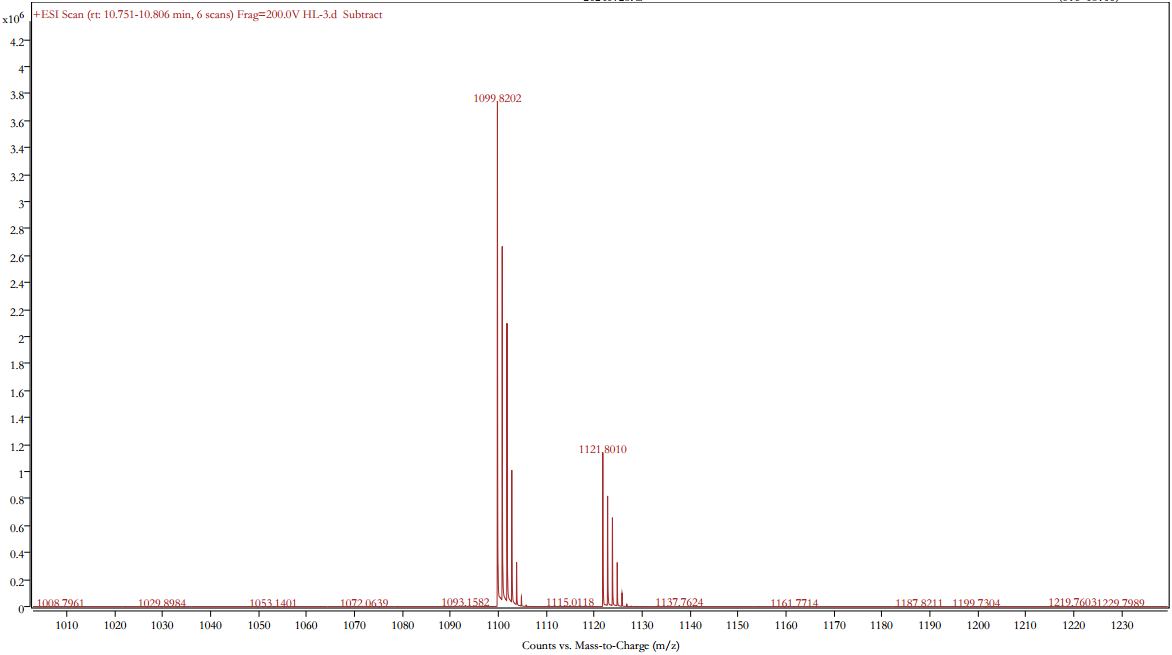

**Supplementary Fig. 11 Characterization of HL-3. ^1^H NMR** (600 MHz, CDCl_3_) δ 8.50 (d, *J* = 5.4 Hz, 1H), 7.94 (d, *J* = 2.1 Hz, 1H), 7.73 (d, *J* = 9.0 Hz, 1H), 7.34 (dd, *J* = 8.9, 2.2 Hz, 1H), 6.42 (d, *J* = 5.5 Hz, 1H), 5.22 (d, *J* = 7.4 Hz, 1H), 4.88 – 4.84 (m, 2H), 4.17 – 4.11 (m, 4H), 3.74 – 3.68 (m, 1H), 2.83 (t, *J* = 7.3 Hz, 4H), 2.72 – 2.68 (m, 4H), 2.60 (s, 4H), 2.55 (q, *J* = 7.1 Hz, 2H), 2.50 (t, *J* = 6.9 Hz, 2H), 2.43 (t, *J* = 9.8, 5.0 Hz, 4H), 1.67 – 1.57 (m, 4H), 1.50 (d, *J* = 6.0 Hz, 8H), 1.32 (d, *J* = 6.3 Hz, 4H), 1.29 – 1.24 (m, 51H), 1.00 (t, *J* = 9.1, 5.1 Hz, 3H), 0.88 (td, *J* = 7.0, 2.3 Hz, 12H). **^13^C NMR** (151 MHz, CDCl_3_) δ 172.27, 172.20, 172.14, 151.95, 149.35, 149.07, 134.78, 128.75, 125.02, 121.23, 117.30, 99.22, 74.50, 62.84, 62.76, 55.75, 53.68, 51.85, 51.56, 49.93, 48.34, 48.19, 34.93, 34.26, 34.10, 33.02, 31.86, 29.55, 29.51, 29.27, 29.01, 28.93, 25.46, 25.33, 24.70, 24.13, 22.67, 20.27, 14.12, 11.65. **MS** (ESI, m/z) [M+H]^+^ calculated for C_64_H_111_ClN_4_O_8_, 1099.81; Found, 1099.8202

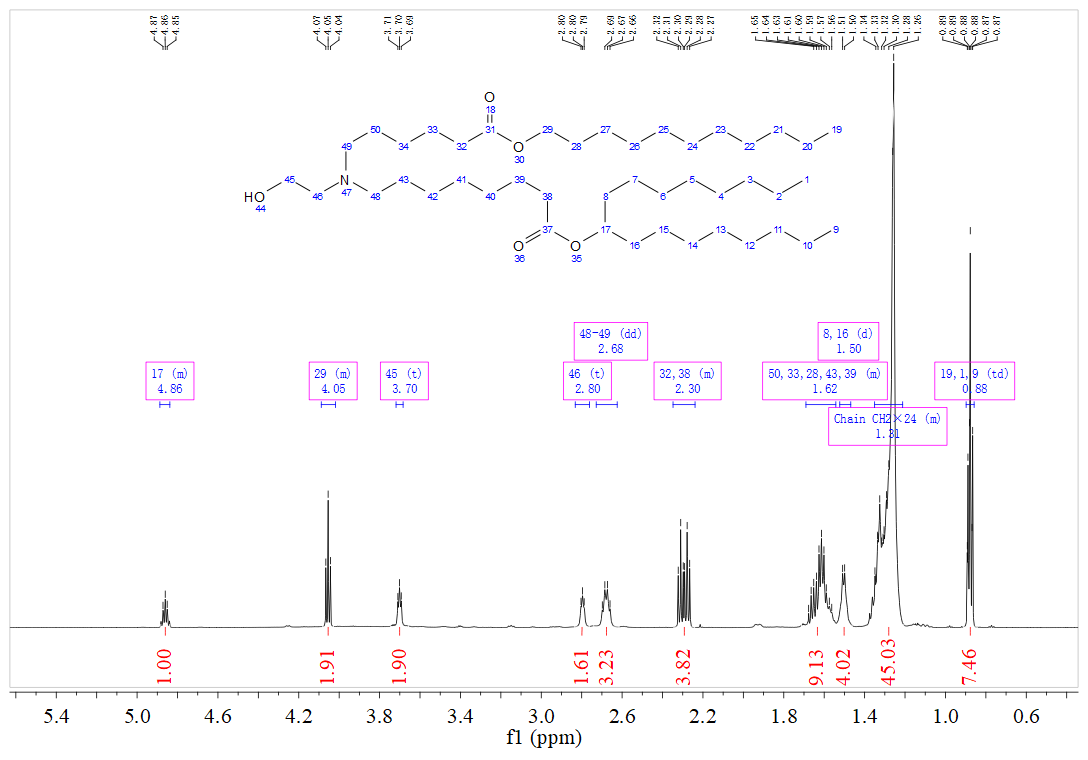

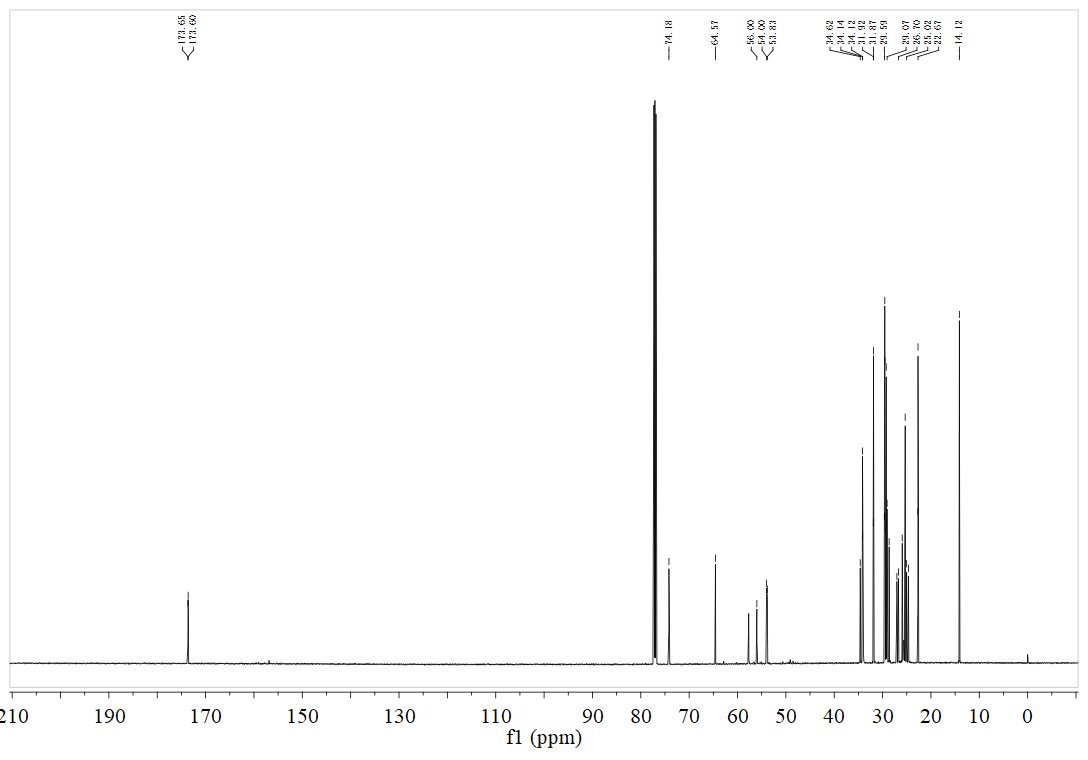

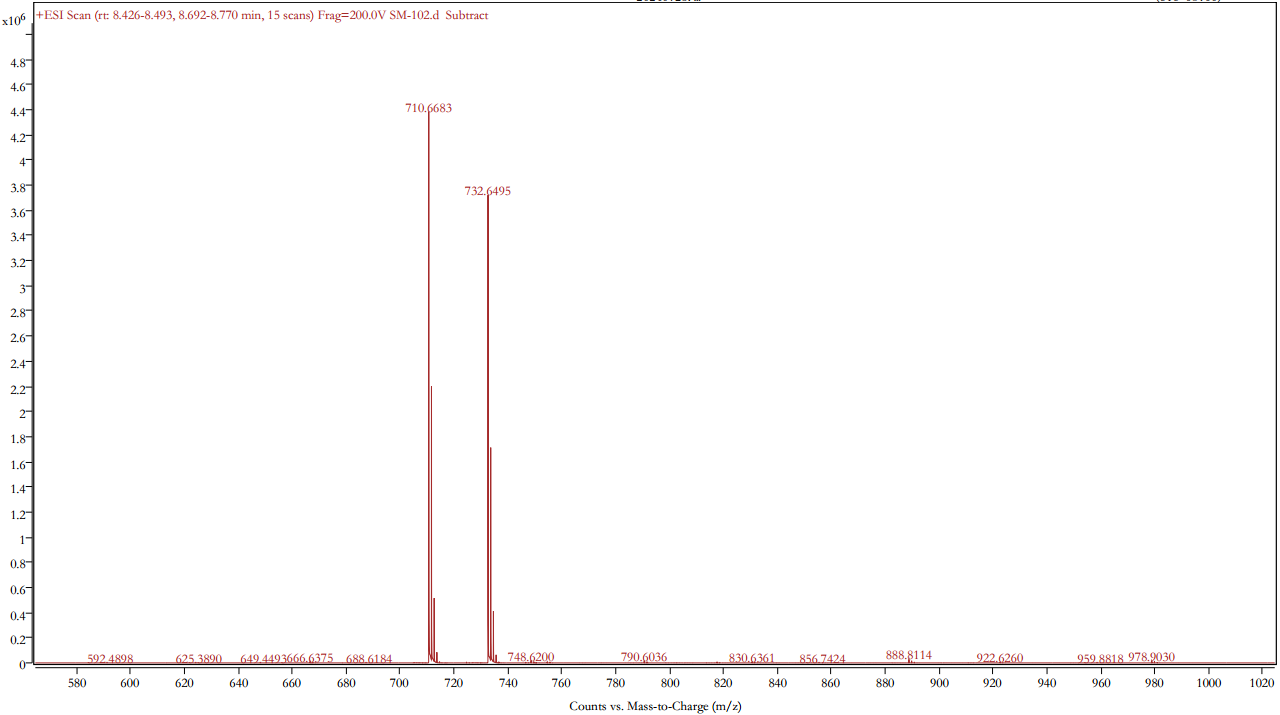

**Supplementary Fig. 12 Characterization of SM-102. ^1^H NMR** (600 MHz, CDCl_3_) δ 4.89 – 4.84 (m, 1H), 4.09 – 4.02 (m, 2H), 3.70 (t, *J* = 5.0 Hz, 2H), 2.80 (t, *J* = 4.7 Hz, 2H), 2.68 (dd, *J* = 15.2, 7.6 Hz, 3H), 2.35 – 2.24 (m, 4H), 1.69 – 1.54 (m, 9H), 1.50 (d, *J* = 6.0 Hz, 4H), 1.35 – 1.21 (m, 45H), 0.88 (td, *J* = 7.0, 1.9 Hz, 7H). **^13^C NMR** (151 MHz, CDCl_3_) δ 173.65, 173.60, 74.18, 64.57, 56.00, 54.00, 53.83, 34.62, 34.14, 34.12, 31.92, 31.87, 29.61, 29.59, 29.54, 29.51, 29.34, 29.27, 29.25, 29.07, 28.65, 27.04, 26.70, 25.94, 25.33, 25.02, 24.65, 22.69, 22.67, 14.12. **MS** (ESI, m/z) [M+H]^+^ calculated for C_44_H_87_NO_5_, 710.66; Found, 710.6683.

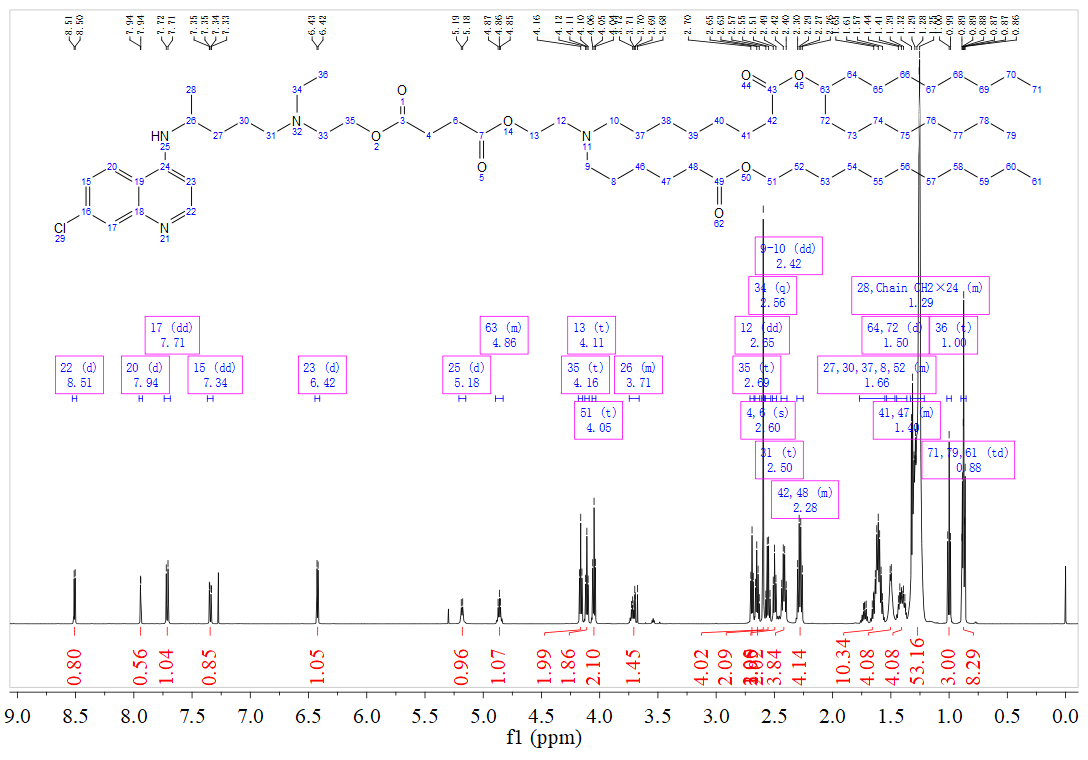

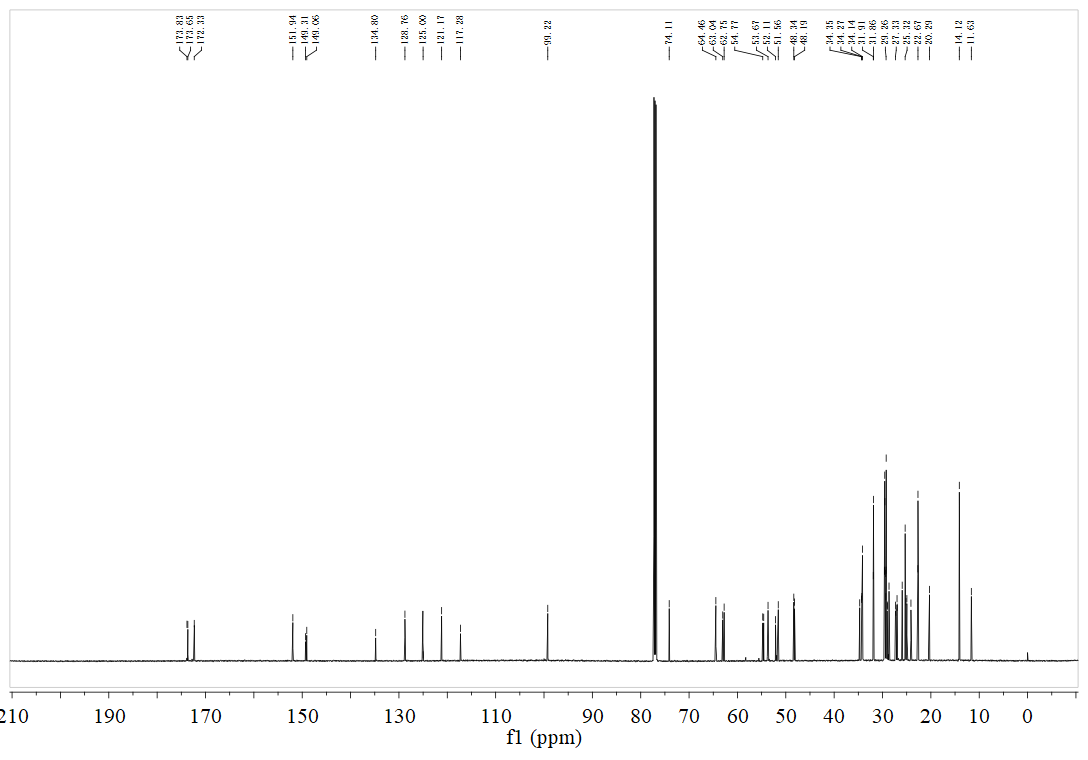

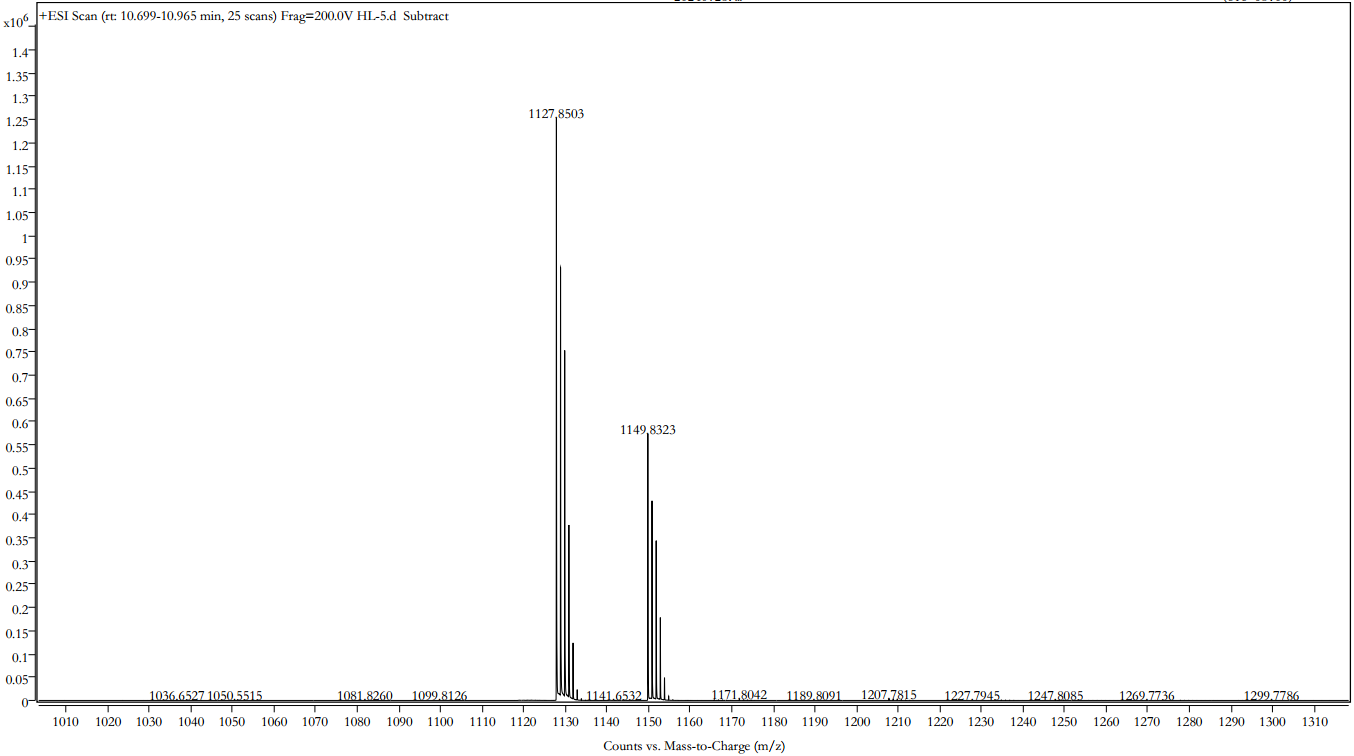

**Supplementary Fig. 13 Characterization of HL-5. ^1^H NMR** (600 MHz, CDCl_3_) δ 8.51 (d, *J* = 5.4 Hz, 1H), 7.94 (d, *J* = 2.1 Hz, 1H), 7.71 (dd, *J* = 9.0, 3.7 Hz, 1H), 7.34 (dd, *J* = 8.9, 2.2 Hz, 1H), 6.42 (d, *J* = 5.5 Hz, 1H), 5.18 (d, *J* = 7.4 Hz, 1H), 4.89 – 4.83 (m, 1H), 4.16 (t, *J* = 6.2 Hz, 2H), 4.11 (t, *J* = 6.4 Hz, 2H), 4.05 (t, *J* = 9.2, 4.4 Hz, 2H), 3.75 – 3.66 (m, 1H), 2.69 (t, *J* = 6.2 Hz, 2H), 2.65 (dd, *J* = 12.9, 6.2 Hz, 2H), 2.60 (s, 4H), 2.56 (q, *J* = 7.1 Hz, 2H), 2.50 (t, *J* = 6.9 Hz, 2H), 2.42 (dd, *J* = 15.1, 7.6 Hz, 4H), 2.31 – 2.25 (m, 4H), 1.77 – 1.55 (m, 10H), 1.50 (d, *J* = 5.9 Hz, 4H), 1.45 – 1.36 (m, 4H), 1.33 – 1.21 (m, 53H), 1.00 (t, *J* = 9.1, 5.1 Hz, 3H), 0.88 (td, *J* = 7.0, 2.4 Hz, 8H). **^13^C NMR** (151 MHz, CDCl_3_) δ 173.83, 173.65, 172.33, 151.94, 149.31, 149.06, 134.80, 128.76, 125.00, 121.17, 117.28, 99.22, 74.11, 64.46, 63.04, 62.75, 54.77, 54.56, 53.67, 52.11, 51.56, 48.34, 48.19, 34.71, 34.35, 34.27, 34.14, 31.91, 31.86, 29.60, 29.58, 29.53, 29.51, 29.33, 29.32, 29.26, 29.24, 29.04, 29.00, 28.65, 27.33, 27.21, 26.99, 26.97, 25.93, 25.32, 25.14, 24.96, 24.12, 22.69, 22.67, 20.29, 14.12, 11.63. **MS** (ESI, m/z) [M+H]^+^ calculated for C_66_H_115_ClN_4_O_8_, 1127.84; Found, 1127.8503.

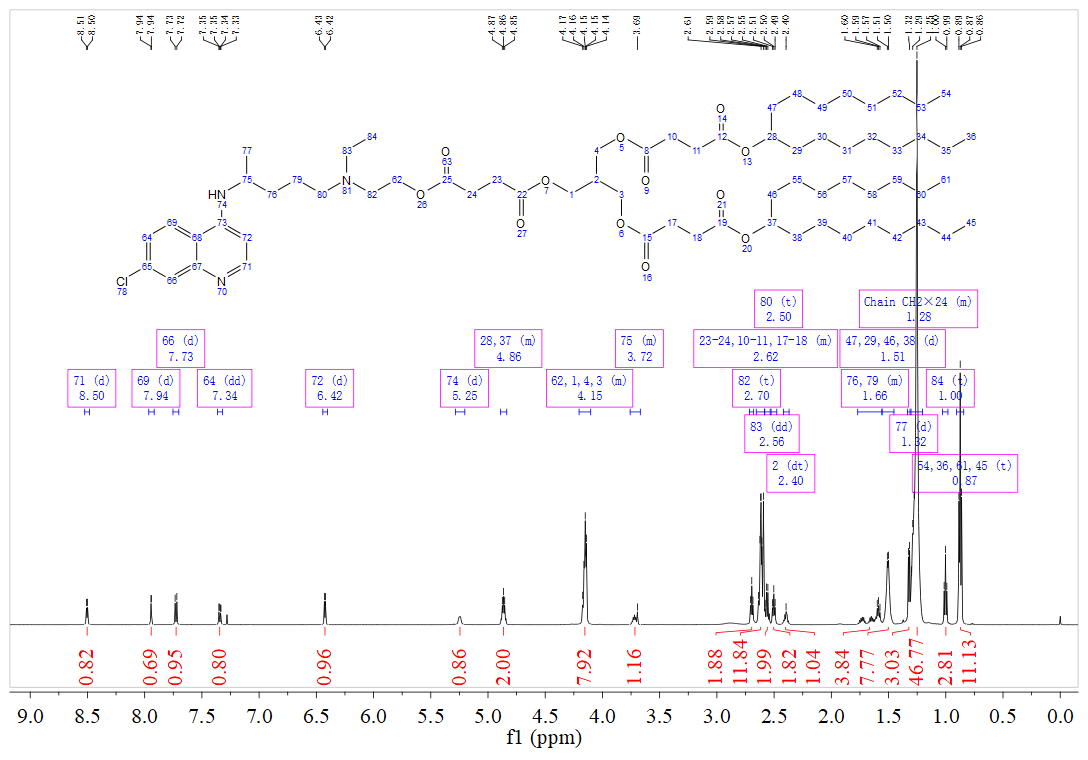

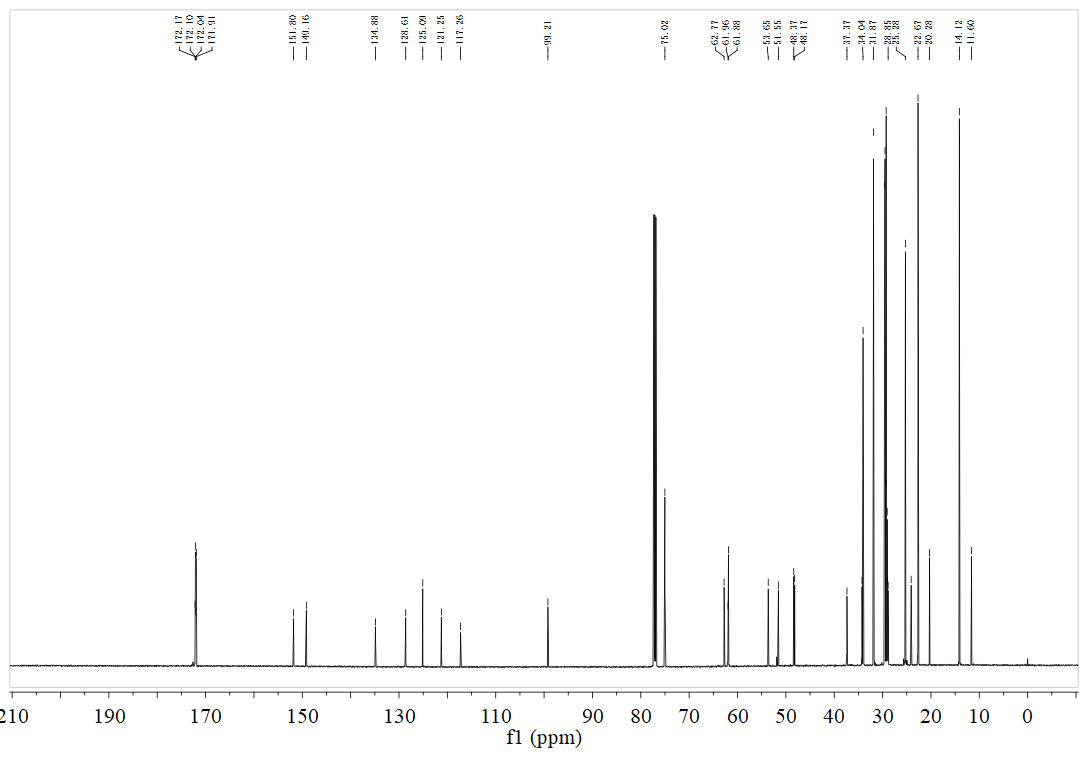

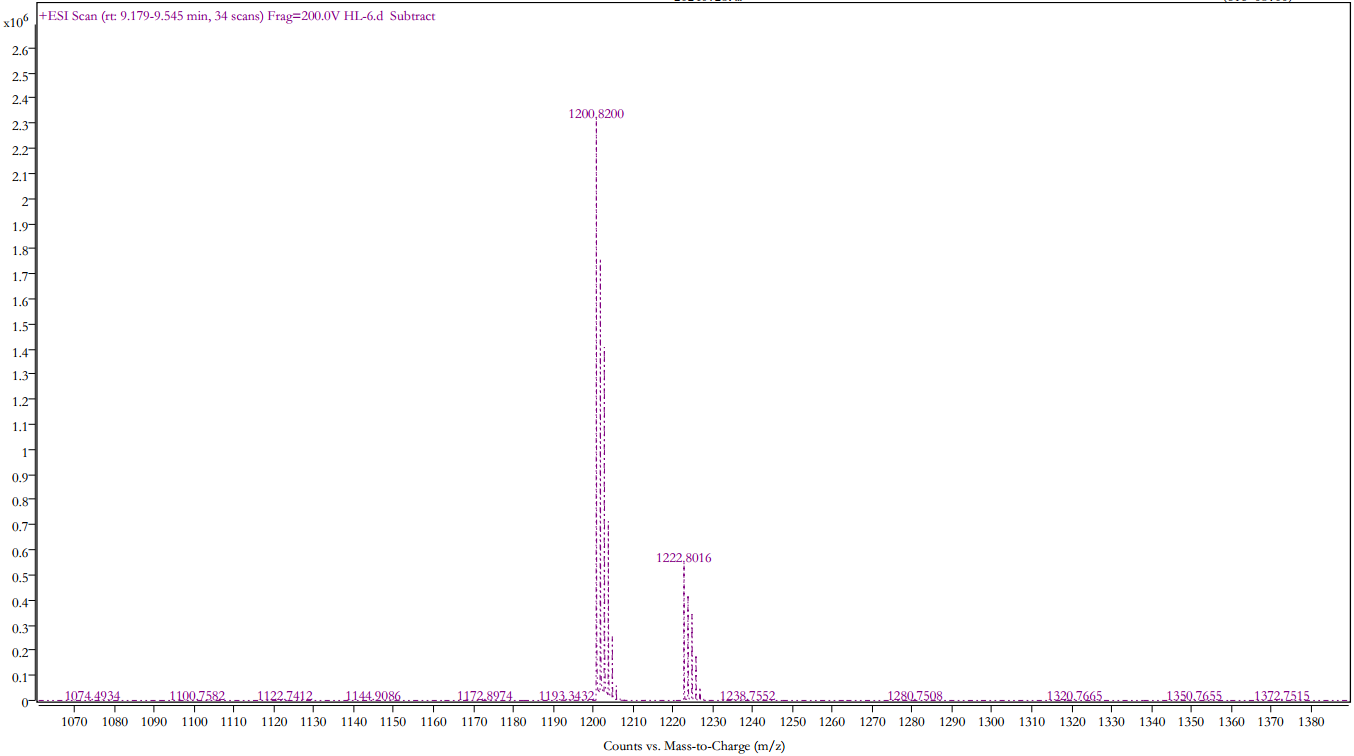

**Supplementary Fig. 14 Characterization of HL-6. ^1^H NMR** (600 MHz, CDCl_3_) δ 8.50 (d, *J* = 5.4 Hz, 1H), 7.94 (d, *J* = 1.7 Hz, 1H), 7.73 (d, *J* = 9.0 Hz, 1H), 7.34 (dd, *J* = 8.9, 1.6 Hz, 1H), 6.42 (d, *J* = 5.5 Hz, 1H), 5.25 (d, *J* = 5.2 Hz, 1H), 4.89 – 4.84 (m, 2H), 4.20 – 4.10 (m, 8H), 3.76 – 3.67 (m, 1H), 2.70 (t, *J* = 6.1 Hz, 2H), 2.66 – 2.58 (m, 12H), 2.56 (dd, *J* = 14.3, 7.2 Hz, 2H), 2.50 (t, *J* = 6.9 Hz, 2H), 2.40 (dt, *J* = 12.0, 5.9 Hz, 1H), 1.77 – 1.56 (m, 4H), 1.51 (d, *J* = 5.8 Hz, 8H), 1.32 (d, *J* = 6.3 Hz, 3H), 1.30 – 1.20 (m, 47H), 1.00 (t, *J* = 7.1 Hz, 3H), 0.87 (t, *J* = 7.0 Hz, 11H). **^13^C NMR** (151 MHz, CDCl_3_) δ 172.17, 172.10, 172.04, 171.91, 151.80, 149.16, 134.88, 128.61, 125.09, 121.25, 117.26, 99.21, 75.02, 62.77, 61.96, 61.88, 53.65, 51.55, 48.37, 48.17, 37.37, 34.26, 34.04, 31.87, 29.54, 29.50, 29.28, 29.26, 29.04, 28.95, 28.85, 25.28, 24.09, 22.67, 20.28, 14.12, 11.60. **MS** (ESI, m/z) [M+H]^+^ calculated for C_68_H_114_ClN_3_O_12_, 1200.81; Found, 1200.8200.

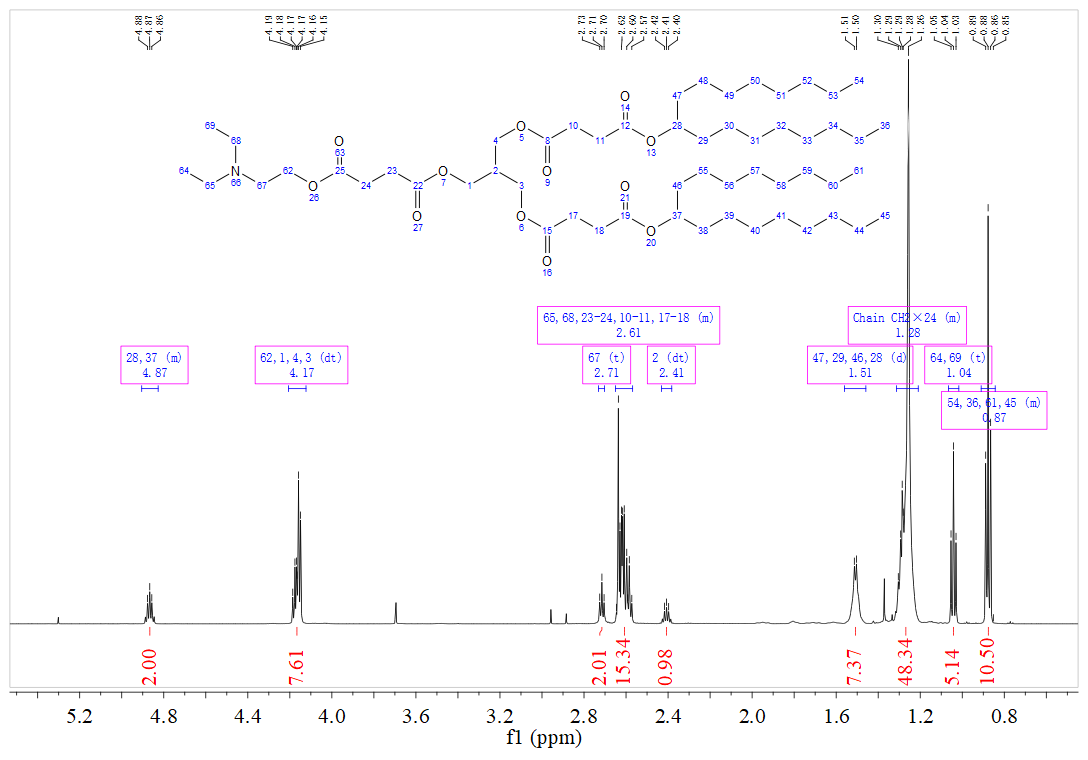

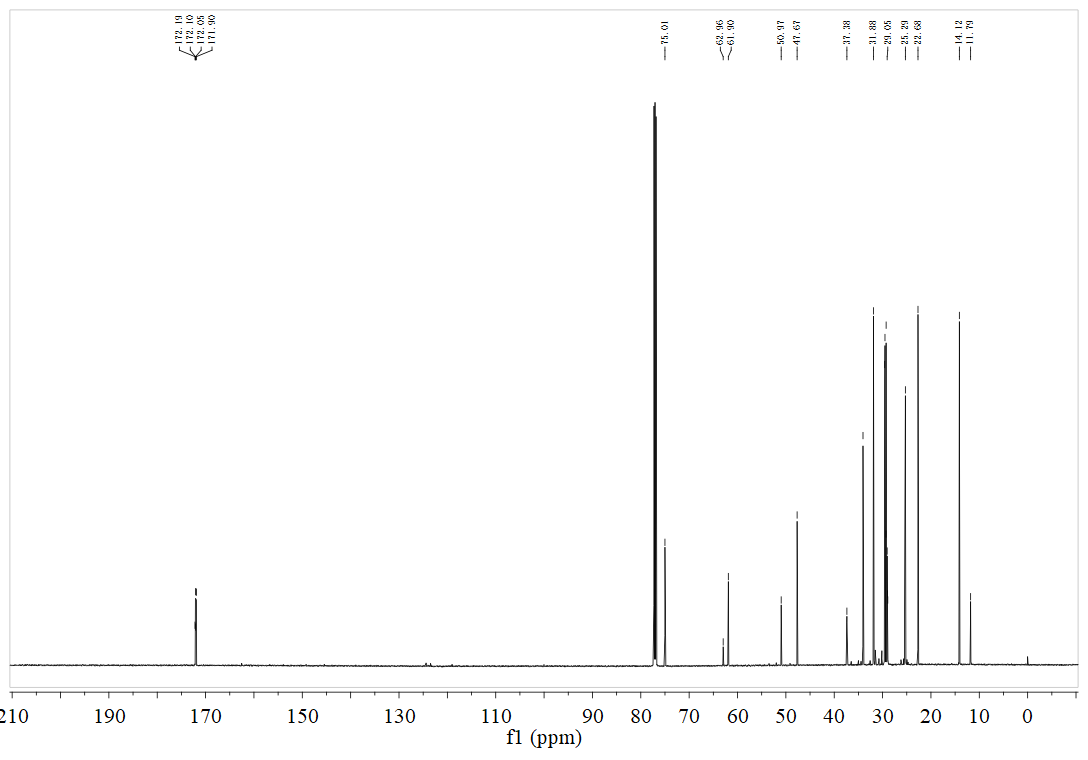

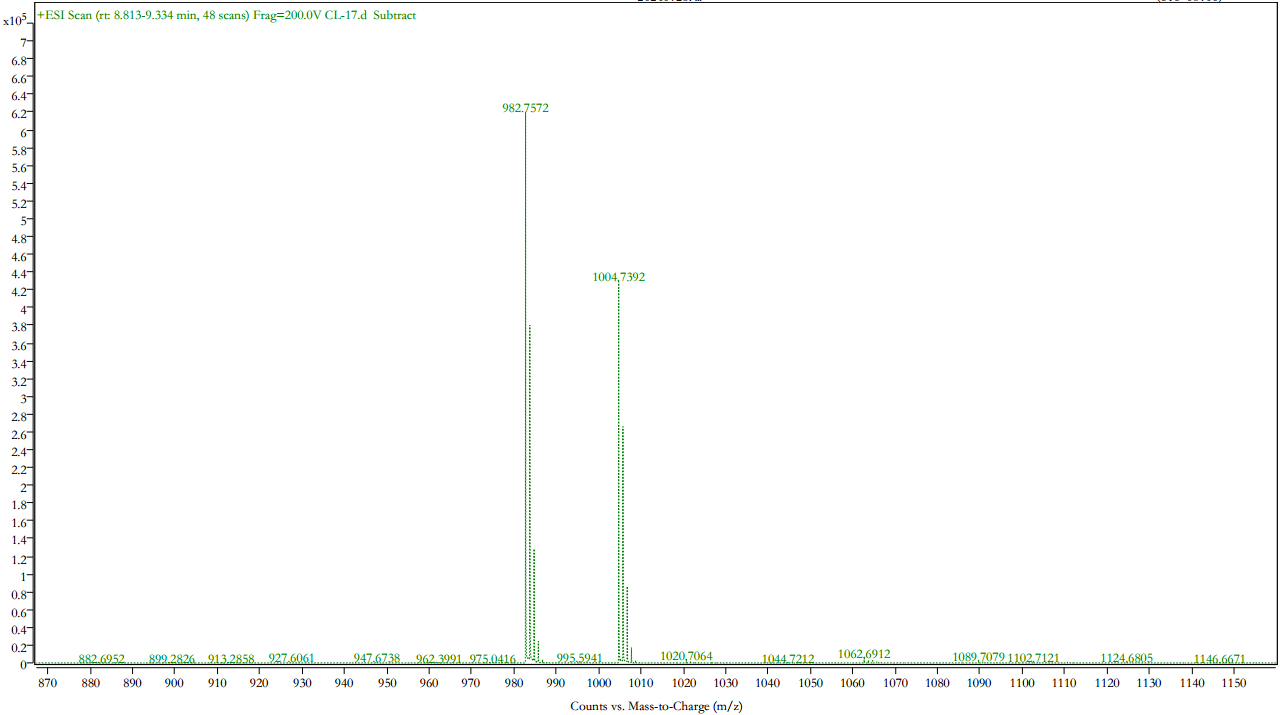

**Supplementary Fig. 15 Characterization of CL-17. ^1^H NMR** (600 MHz, CDCl_3_) δ 4.90 – 4.83 (m, 2H), 4.17 (dt, *J* = 10.1, 6.1 Hz, 8H), 2.71 (t, *J* = 6.2 Hz, 2H), 2.65 – 2.57 (m, 15H), 2.41 (dt, *J* = 12.1, 6.0 Hz, 1H), 1.51 (d, *J* = 6.0 Hz, 7H), 1.31 – 1.21 (m, 48H), 1.04 (t, *J* = 7.1 Hz, 5H), 0.91 – 0.84 (m, 11H). **^13^C NMR** (151 MHz, CDCl_3_) δ 172.19, 172.10, 172.05, 171.90, 75.01, 62.96, 61.90, 50.97, 47.67, 37.38, 34.05, 31.88, 29.55, 29.50, 29.29, 29.27, 29.05, 29.00, 28.92, 25.29, 22.68, 14.12, 11.79. **MS** (ESI, m/z) [M+H]^+^ calculated for C_56_H_103_NO_12_, 982.75; Found, 982.7572.

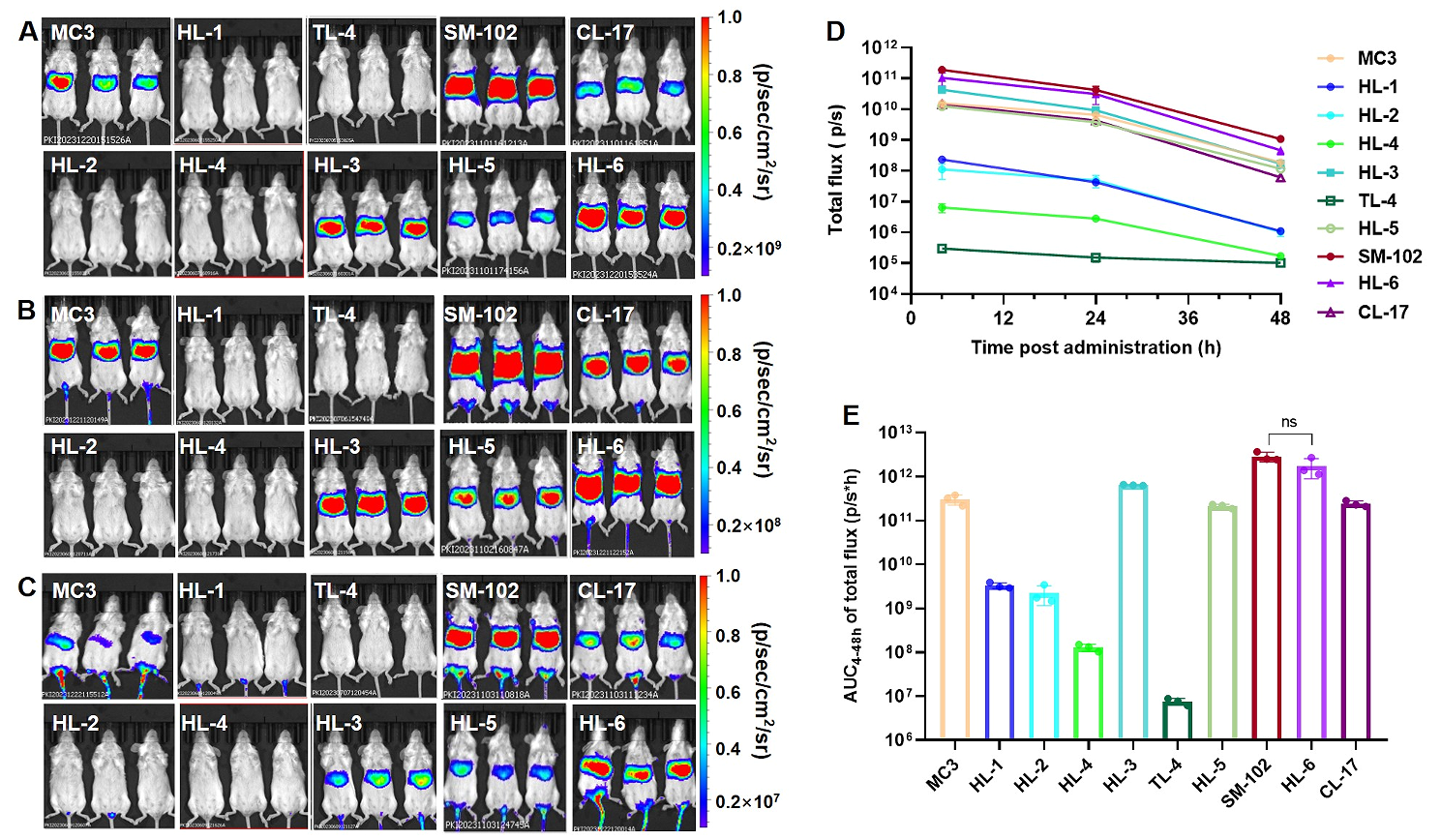

**Supplementary Fig. 16 Bioluminescence-time curve and AUC after intravenous administration.** A-C) Bioluminescence images at 4 h, 24 h and 48 h, respectively. D) Bioluminescence-time curve of different LNPs. E) AUC_4-48 h_ of bioluminescence. The result of AUC matched well with the bioluminescence intensity at 4 h.

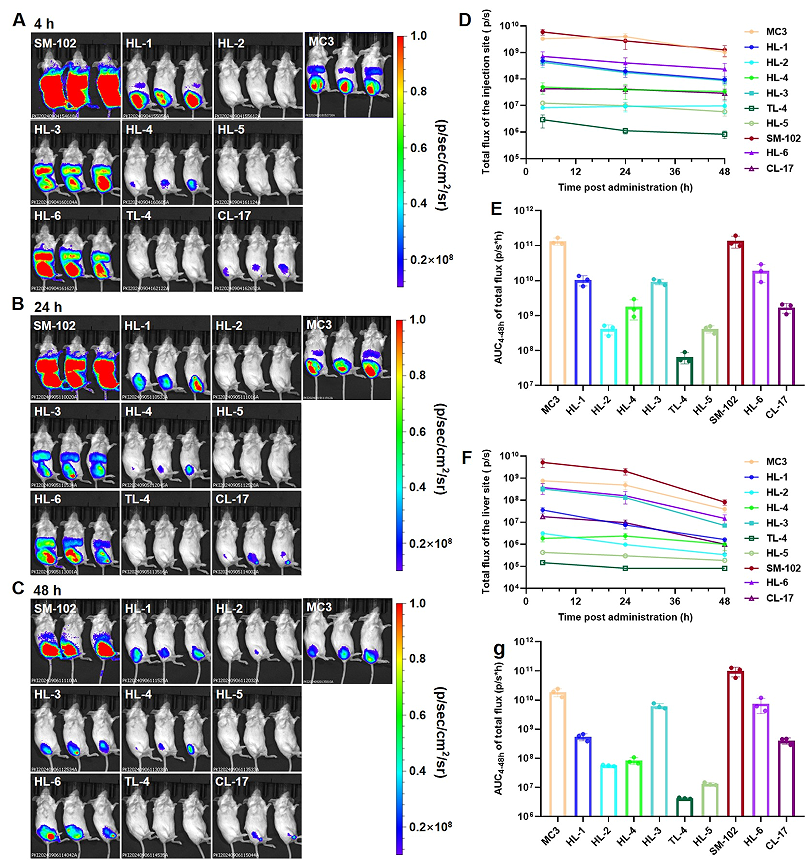

**Supplementary Fig. 17** **Bioluminescence-time curve and AUC after intramuscular administration.** A-C) Bioluminescence images at 4 h, 24 h and 48 h, respectively. D) Bioluminescence-time curve of the injection site. E) AUC_4-48 h_ of bioluminescence of the injection site. F) Bioluminescence-time curve of the liver site. G) AUC_4-48 h_ of bioluminescence of the liver site. The result of AUC matched well with the bioluminescence intensity at 4 h.

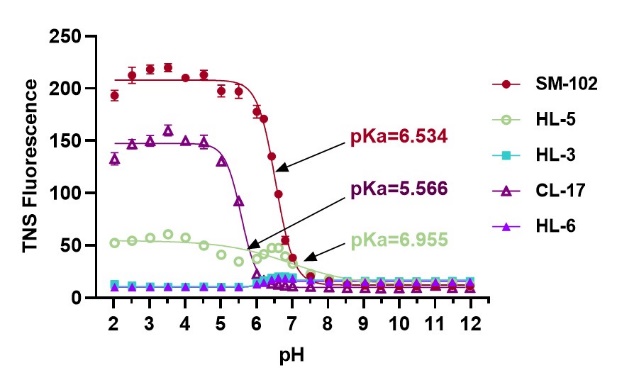

**Supplementary Fig. 18 TNS fluorescence-pH curve and Apparent pKa values of LNPs without mRNA.**

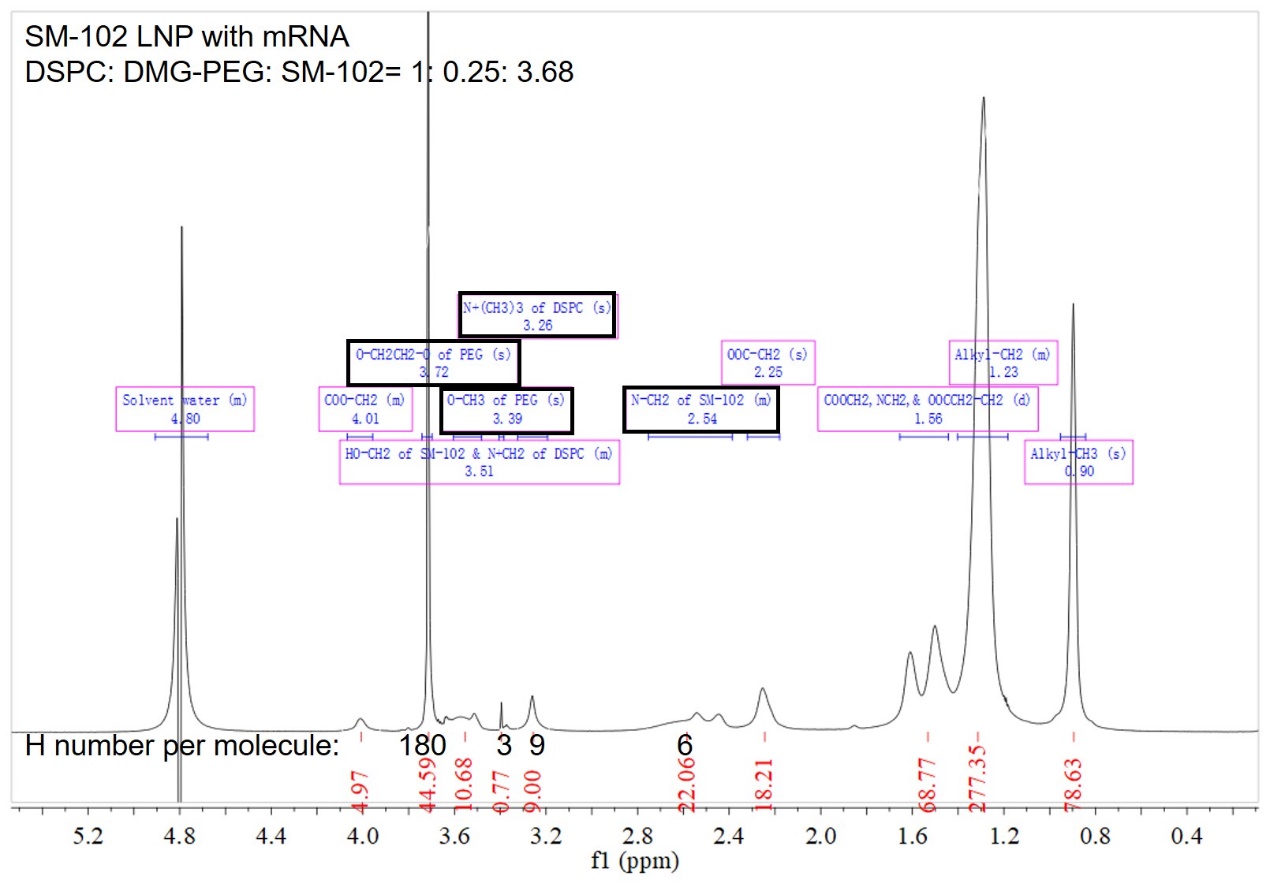

**Supplementary Fig. 19 ^1^H NMR spectrum of SM-102 LNP with mRNA and the proportion of the lipids distributed on the surface of SM-102 LNP with mRNA.**

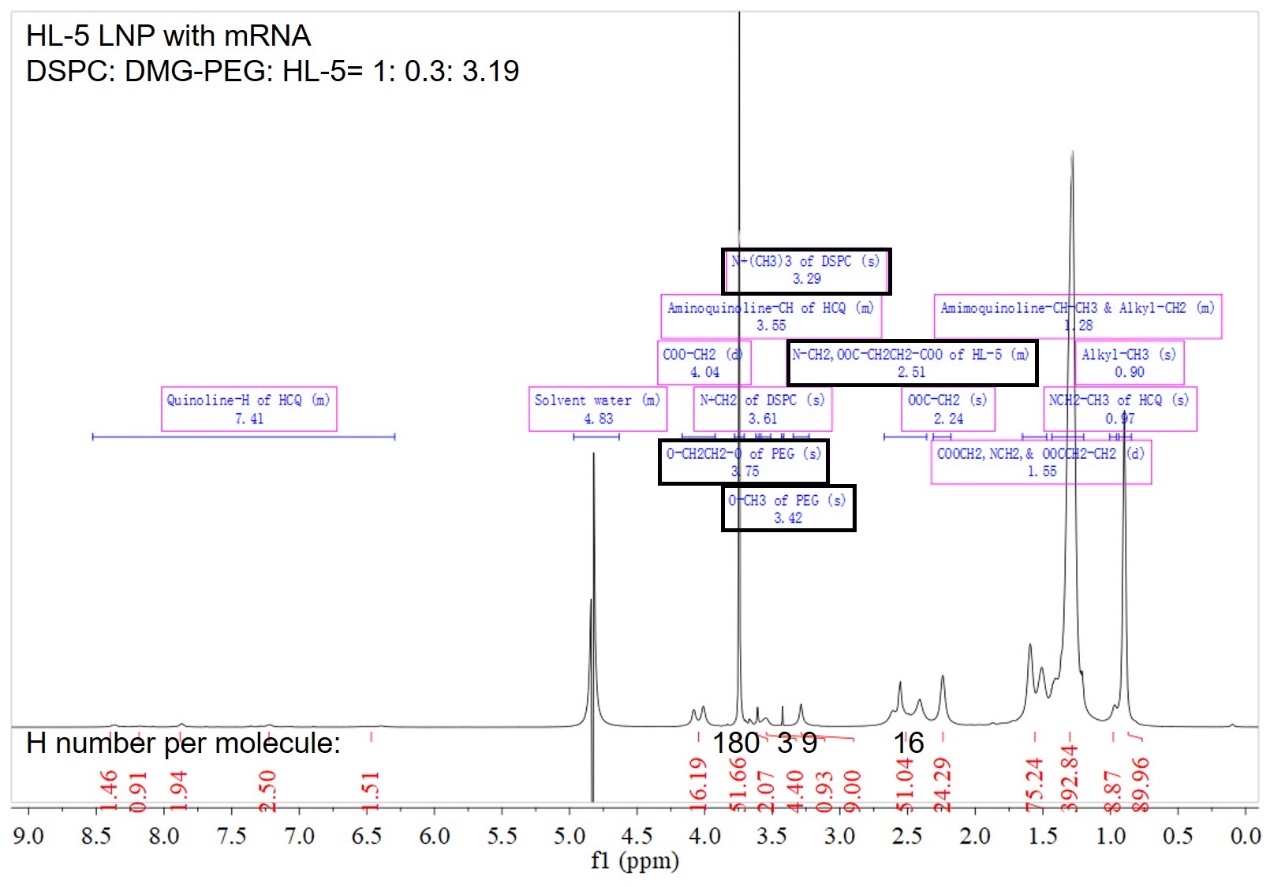

**Supplementary Fig. 20 ^1^H NMR spectrum of HL-5 LNP with mRNA and the proportion of the lipids distributed on the surface of HL-5 LNP with mRNA.**

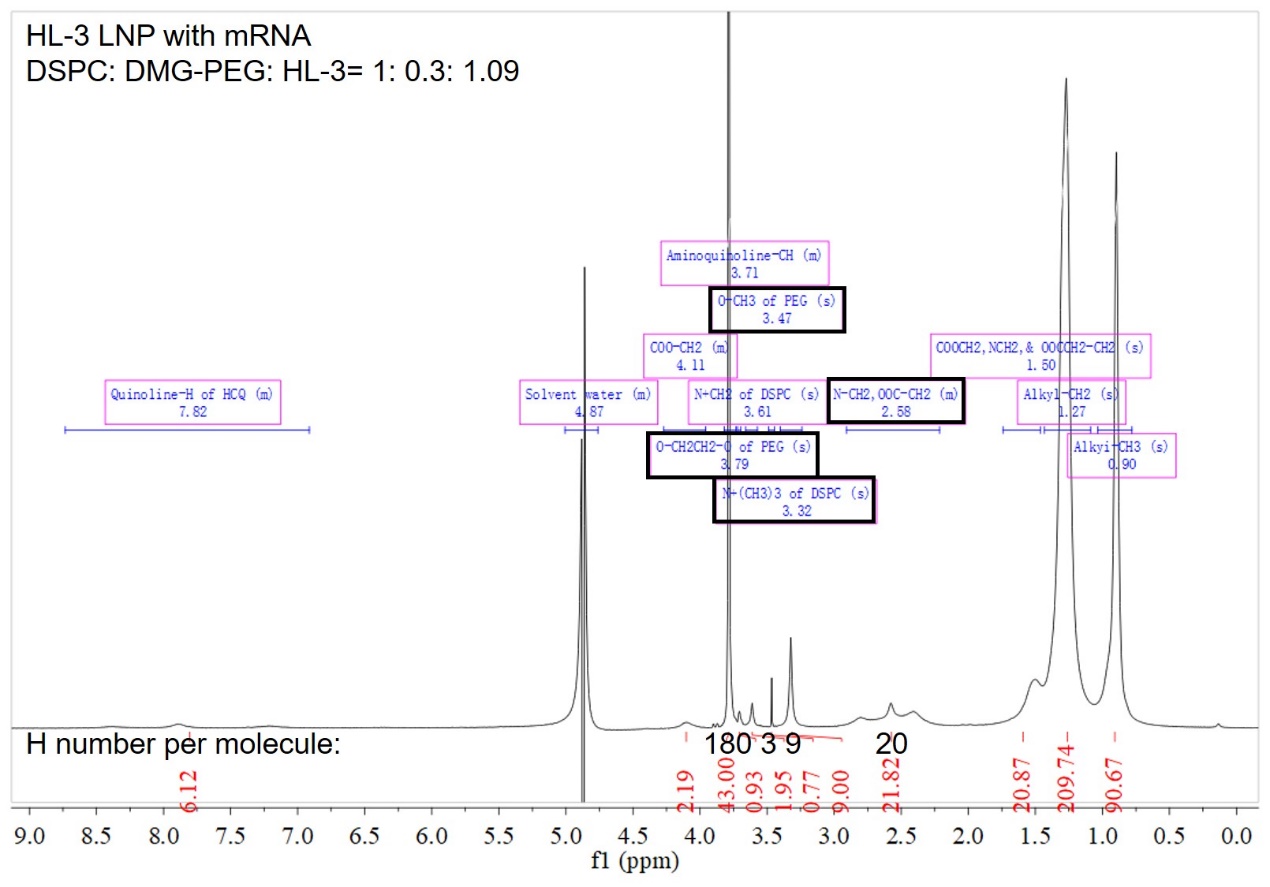

**Supplementary Fig. 21 ^1^H NMR spectrum of HL-3 LNP with mRNA and the proportion of the lipids distributed on the surface of HL-3 LNP with mRNA.**

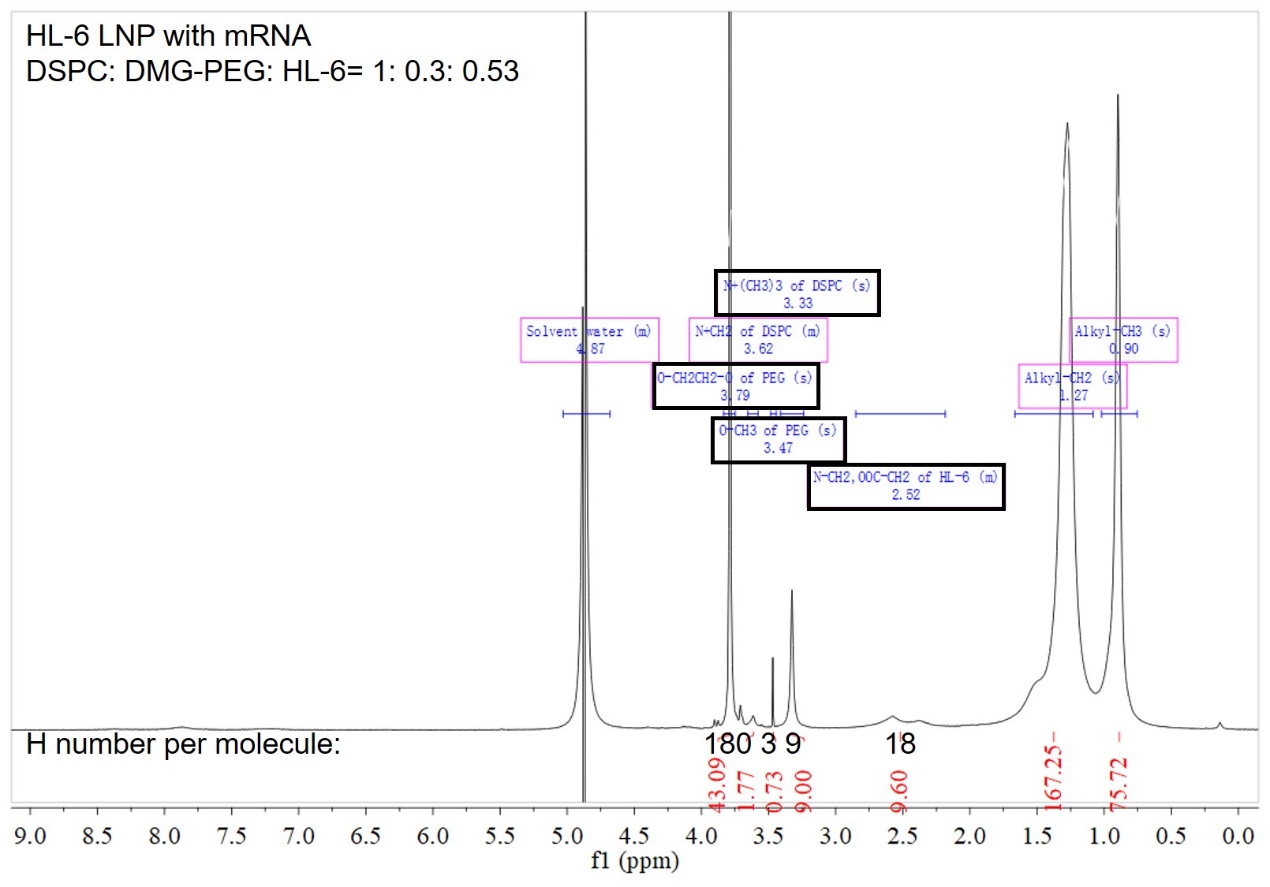

**Supplementary Fig. 22 ^1^H NMR spectrum of HL-6 LNP with mRNA and the proportion of the lipids distributed on the surface of HL-3 LNP with mRNA.**

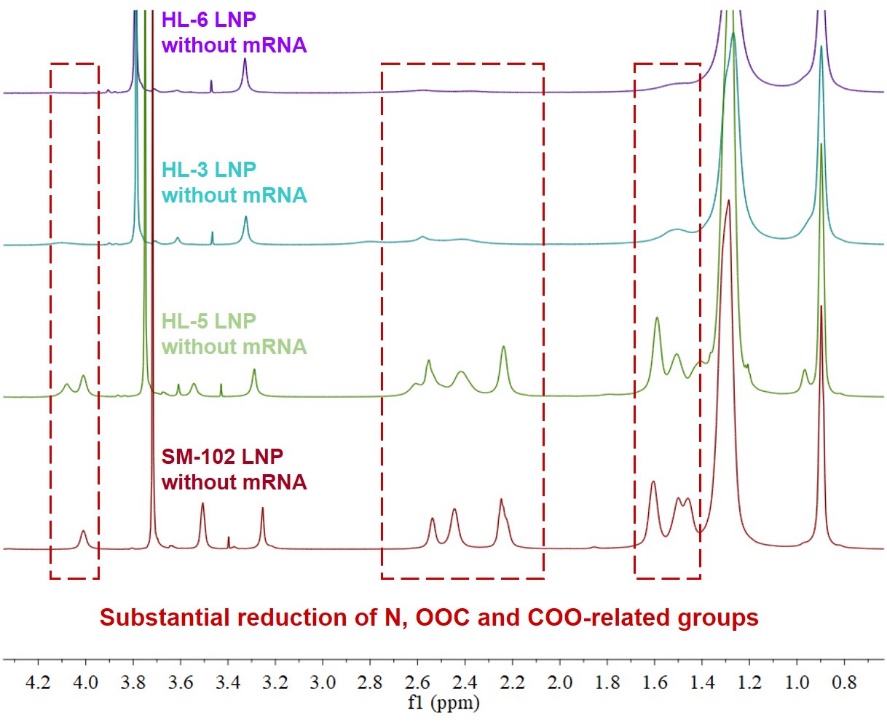

**Supplementary Fig. 23 Merged ^1^H NMR spectra of different LNPs without mRNA.**

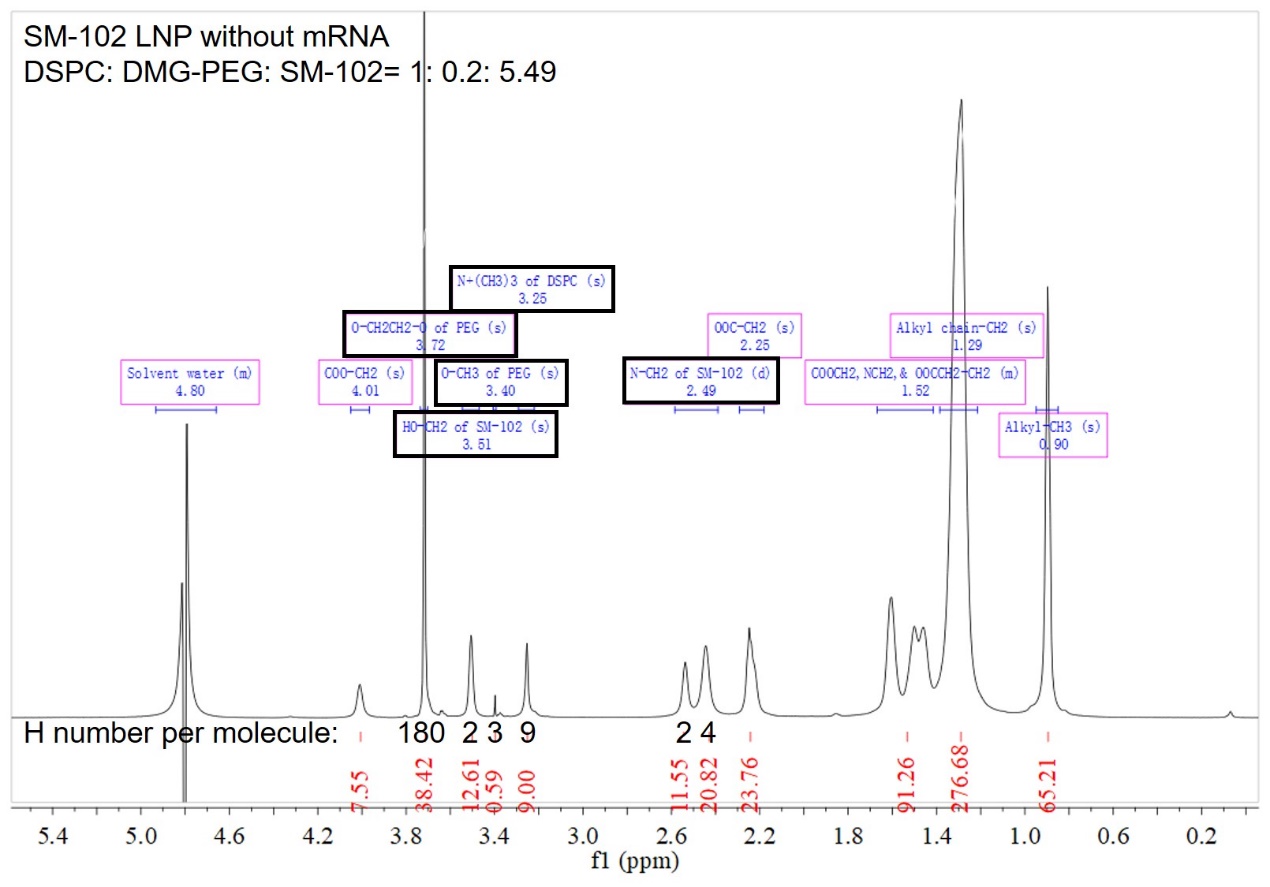

**Supplementary Fig. 24 ^1^H NMR spectrum of SM-102 LNP without mRNA and the proportion of the lipids distributed on the surface of SM-102 LNP without mRNA.**

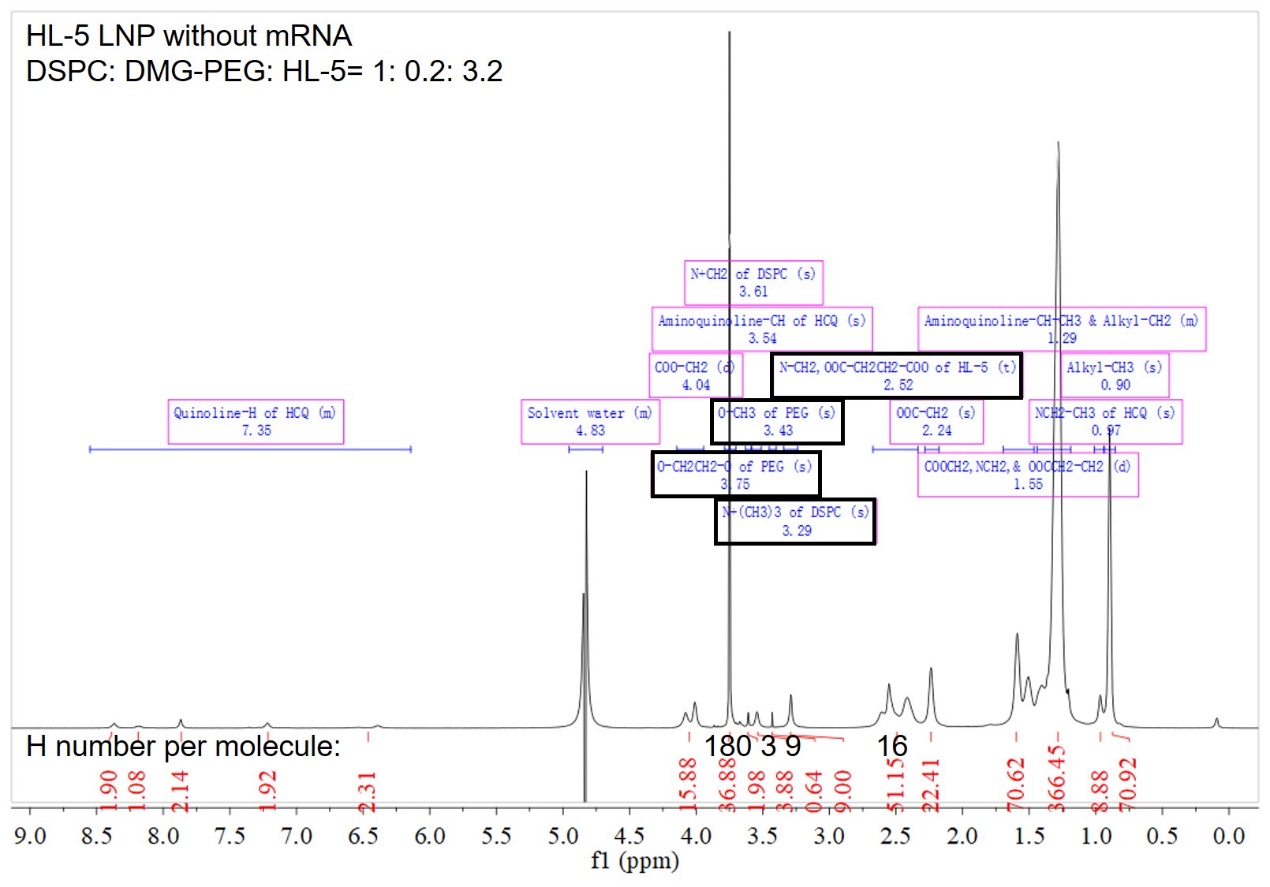

**Supplementary Fig. 25 ^1^H NMR spectrum of HL-5 LNP without mRNA and the proportion of the lipids distributed on the surface of HL-5 LNP without mRNA.**

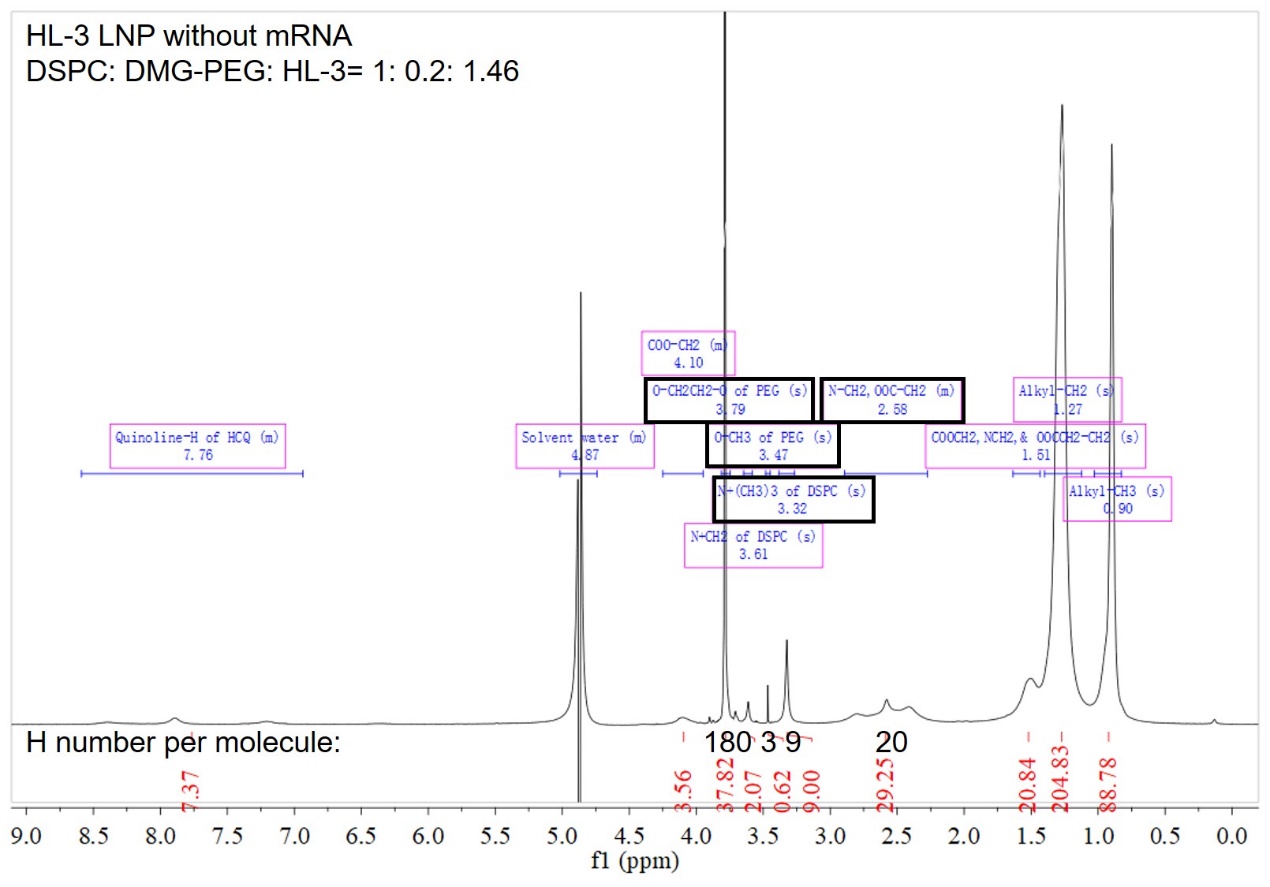

**Supplementary Fig. 26 ^1^H NMR spectrum of HL-3 LNP without mRNA and the proportion of the lipids distributed on the surface of HL-3 LNP without mRNA.**

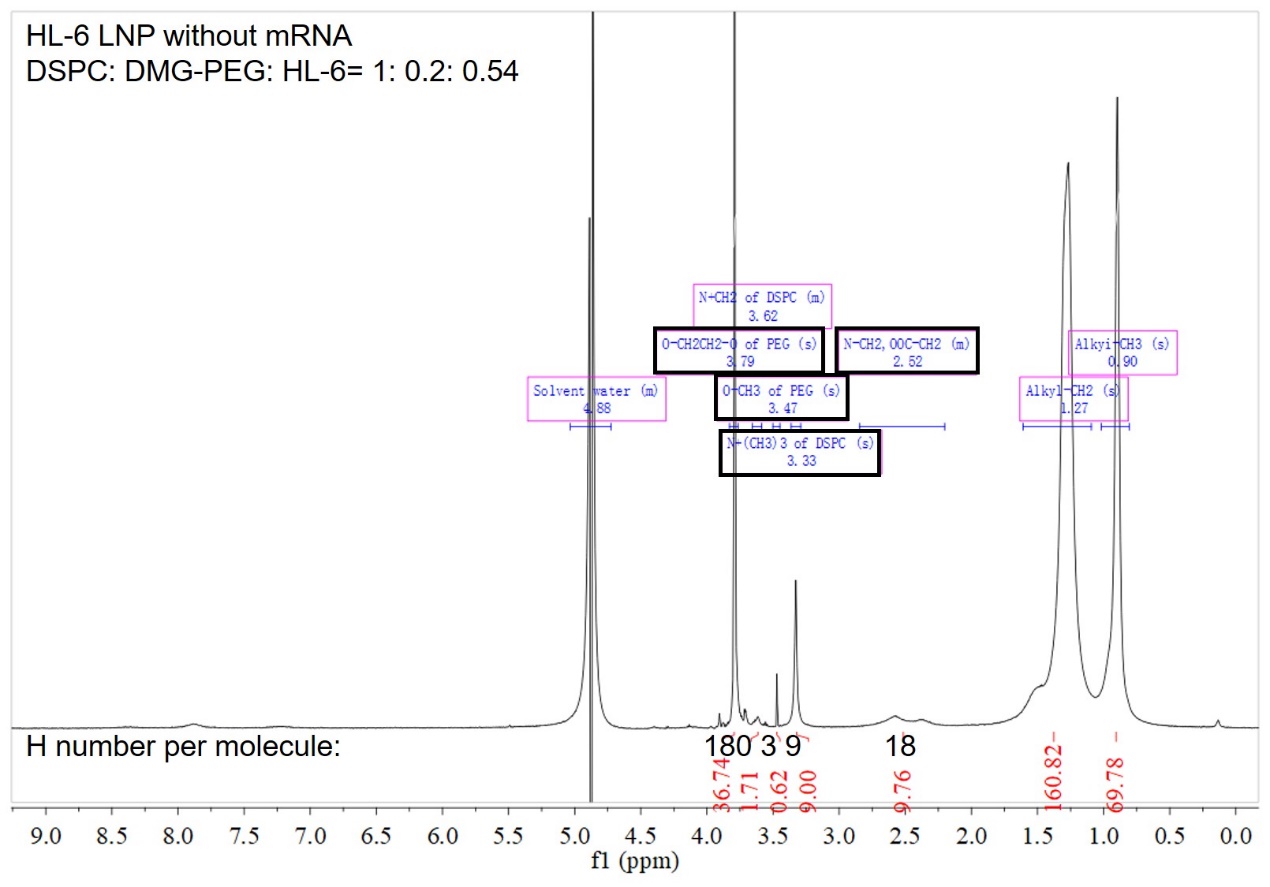

**Supplementary Fig. 27 ^1^H NMR spectrum of HL-6 LNP without mRNA and the proportion of the lipids distributed on the surface of HL-3 LNP without mRNA.**

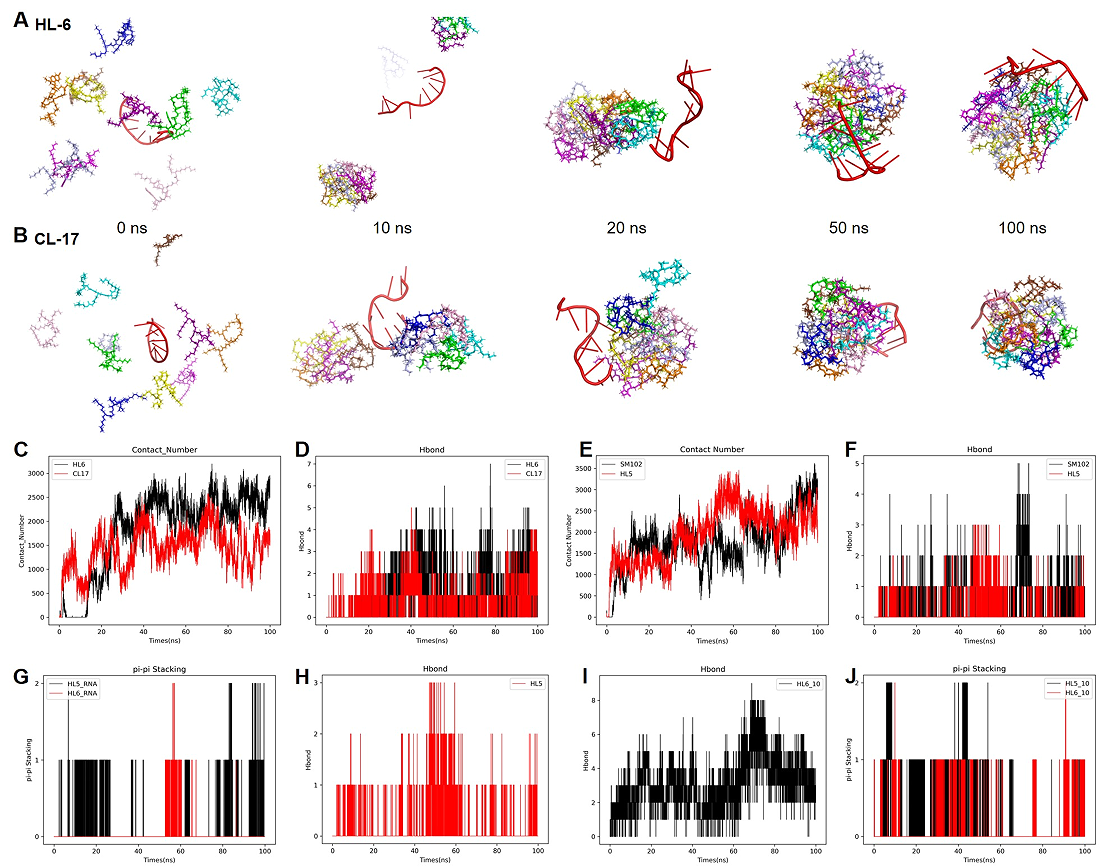

**Supplementary Fig. 28 Molecular dynamics simulation.** A-B) Simulation snapshots for HL-6 (A) and CL-17 (B) captured at 0 ns, 10 ns, 50 ns, and 100 ns. C) Number of contacts between nucleotides and either HL-6 or CL-17. D) Hydrogen bonding between nucleotides and either HL-6 or CL-17. E) Number of contacts between nucleotides and either SM-102 or HL-5. F) Hydrogen bonding between nucleotides and either SM-102 or HL-5. G) π-π stacking interaction between nucleotides and the aminoquinoline ring of either HL-5 or HL-6. H) Hydrogen bonding among HL-5 themselves. I) Hydrogen bonding among HL-6 themselves. J) π-π stacking interaction among HL themselves.

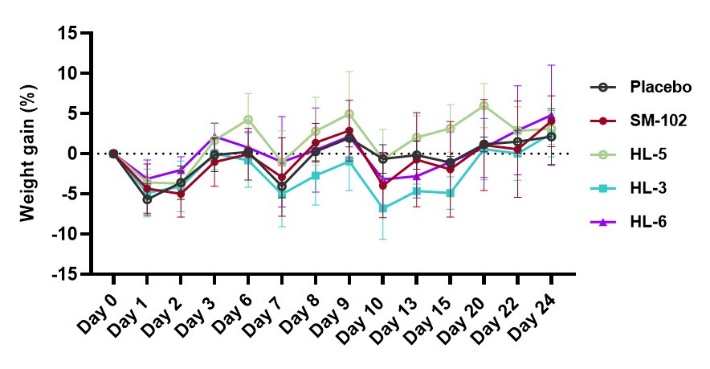

**Supplementary Fig. 29 Weight growth curve during the repeated administration.**

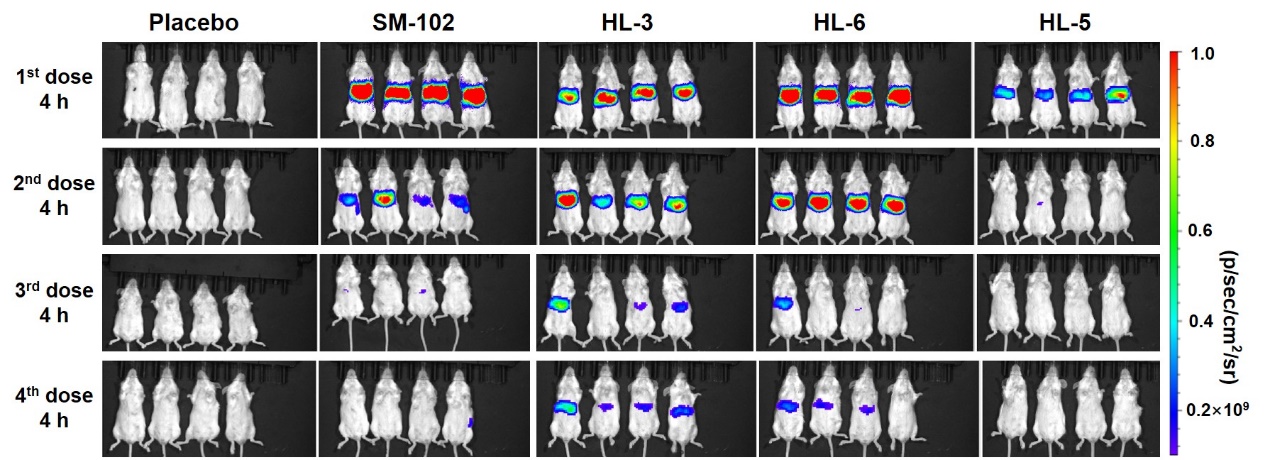

**Supplementary Fig. 30 Bioluminescence images at 4 h after each dose.**

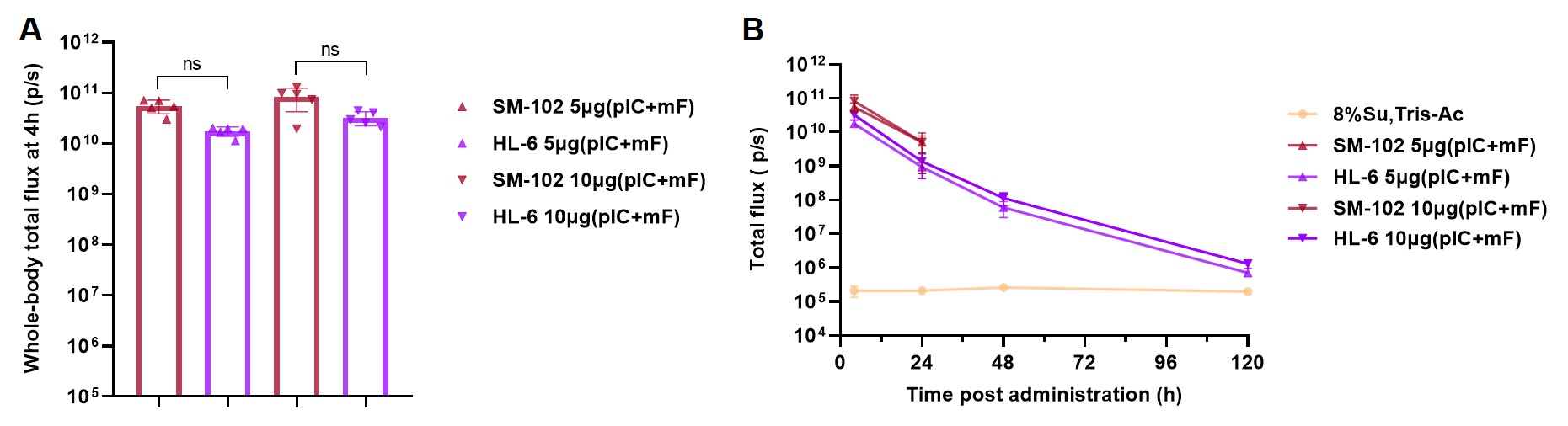

**Supplementary Fig. 31** **Bioluminescence monitoring after the administration of the mixture of poly(I:C) and FLuc mRNA.** BALB/c mice were injected with LNPs co-loaded with the mixture nucleic acid at a total dose of 5μg or 10 μg per mouse (n = 5). A) Bioluminescence at 4 h after administration. B) Bioluminescence curves from 4 h to 120h.

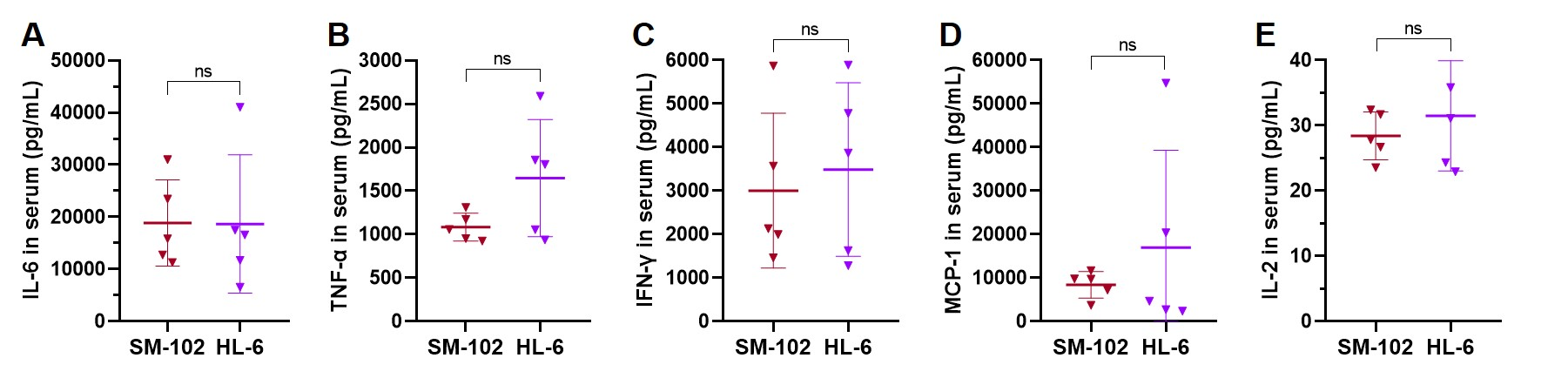
**Supplementary Fig. 32 Serum concentrations of proinflammatory cytokines at 4 h after administration in the 5 μg groups.** A) IL-6. B) TNF-α. C) IFN-γ. D) MCP-1. E) IL-2.

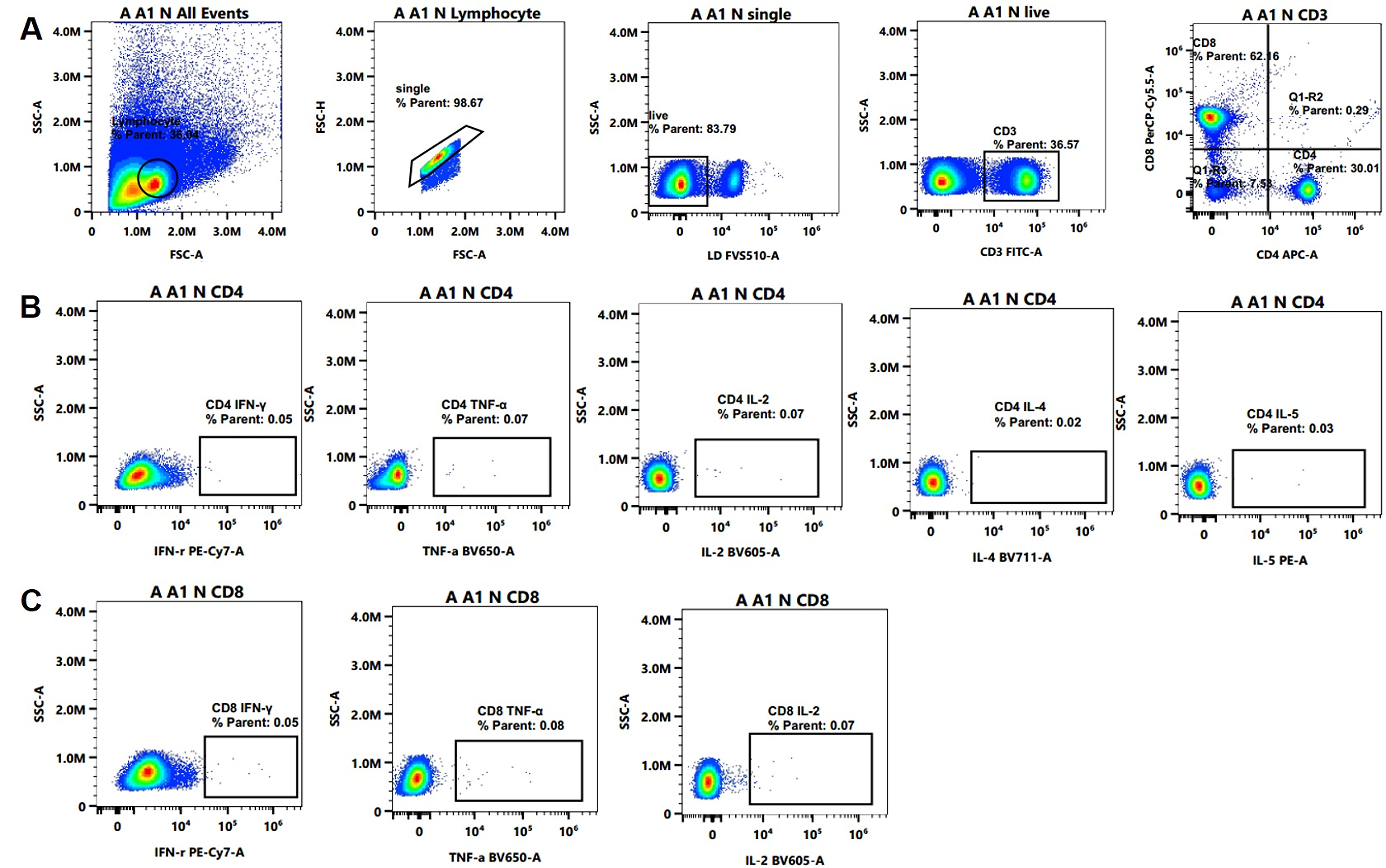

**Supplementary Fig. 33 Gating strategy for the intracellular cytokine staining.** A) Gating scheme for the identification of CD4^+^ and CD8^+^ T cells. B) Identification of antigen-specific cytokine-expressing CD4^+^ T cells. C) Identification of antigen-specific cytokine-expressing CD8^+^ T cells.

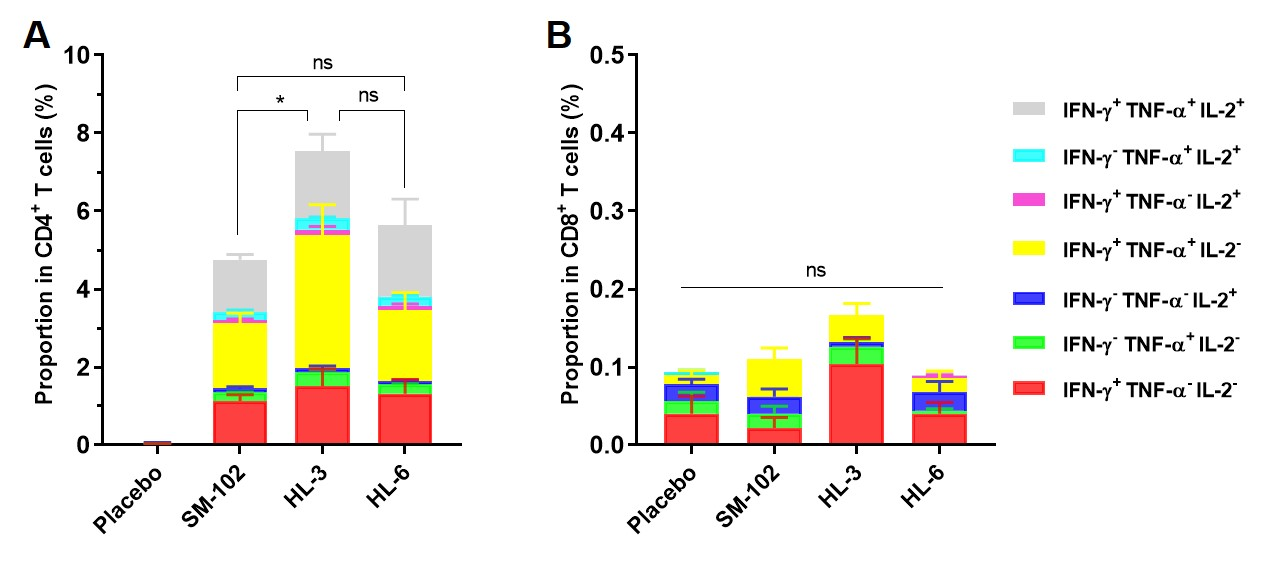

**Supplementary Fig. 34 Total gE-specific T cells quantified by ICS.** A) Total gE-specific CD4^+^ T cells. B) Total gE-specific CD8^+^ T cells.

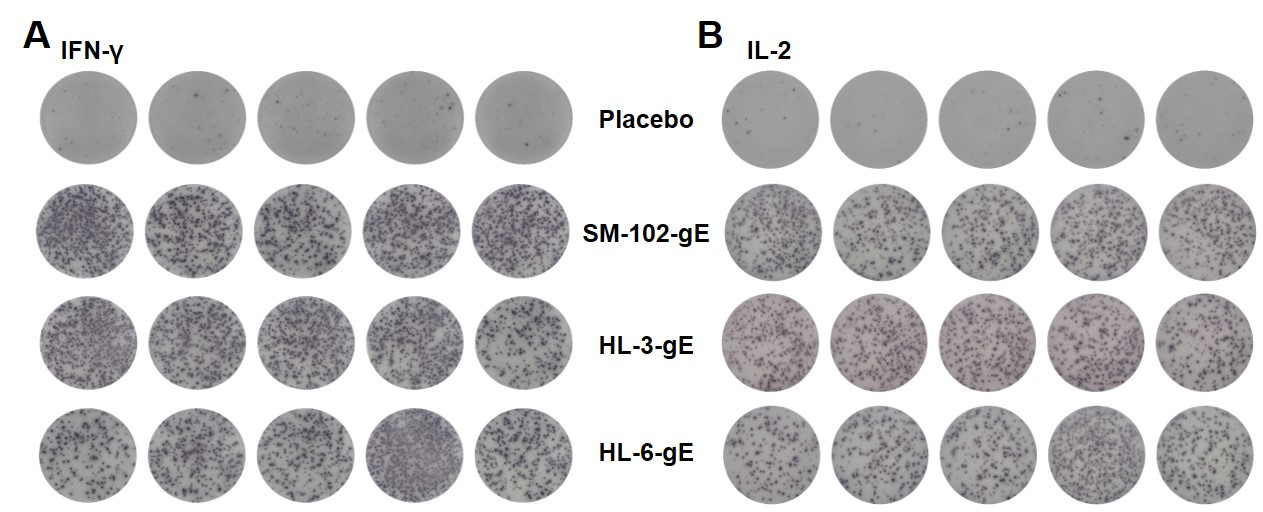

**Supplementary Fig. 35 Spot images of cytokine-secreting splenocytes.** A) IFN-γ-secreting splenocytes. B) IL-2-secreting splenocytes.

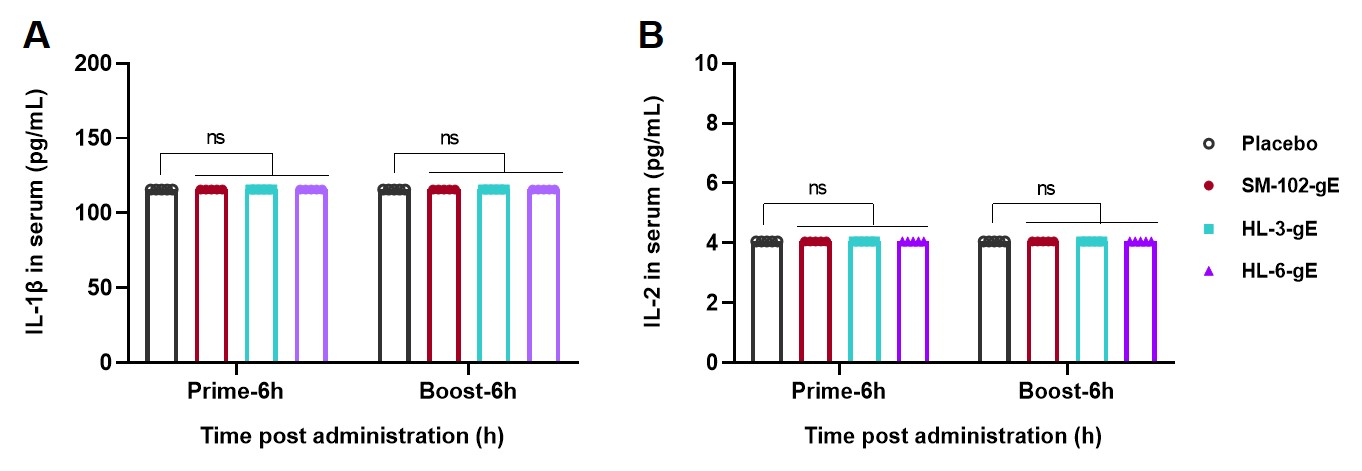

**Supplementary Fig. 36 Release of IL-1β (A) and IL-2 (B) after immunization.**

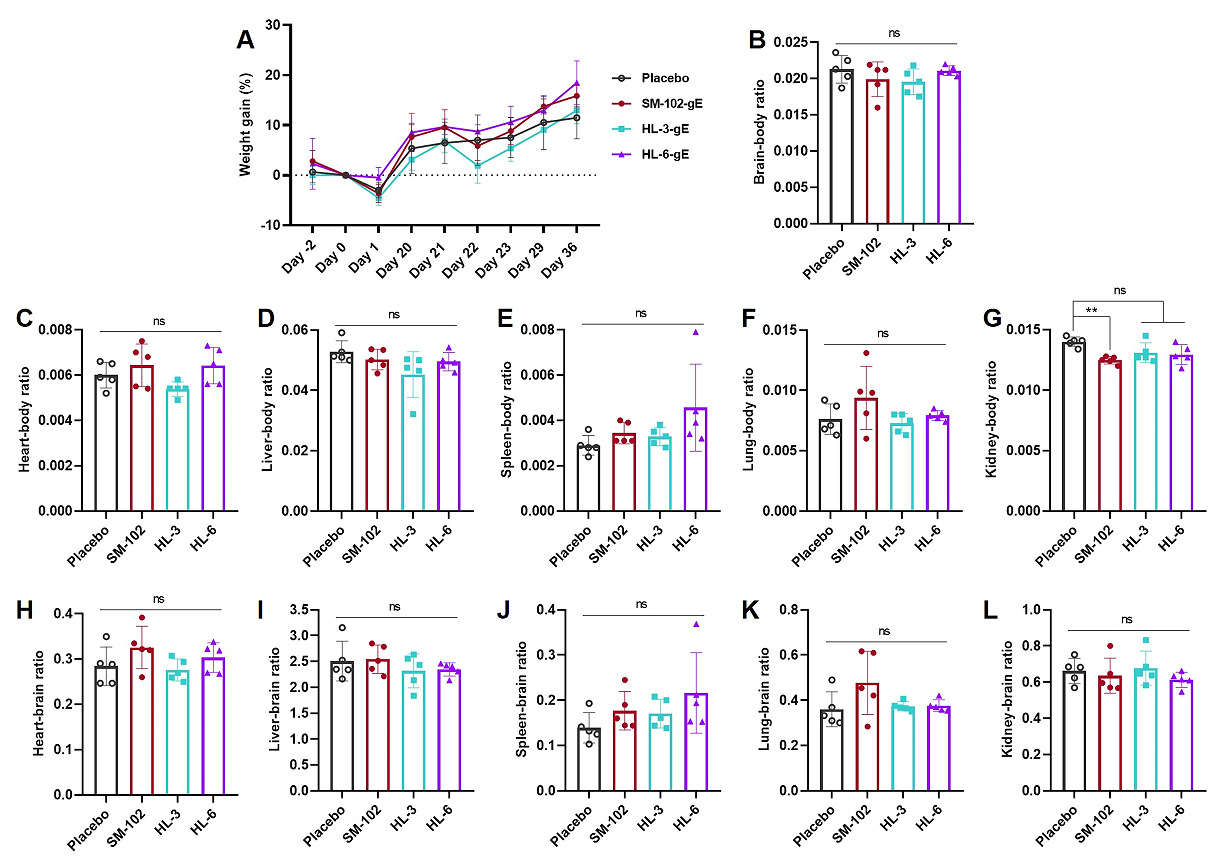

**Supplementary Fig. 37 Weight gain curve (A), coefficients of organ-body (B-G), and coefficients of organ-brain (H-L).**

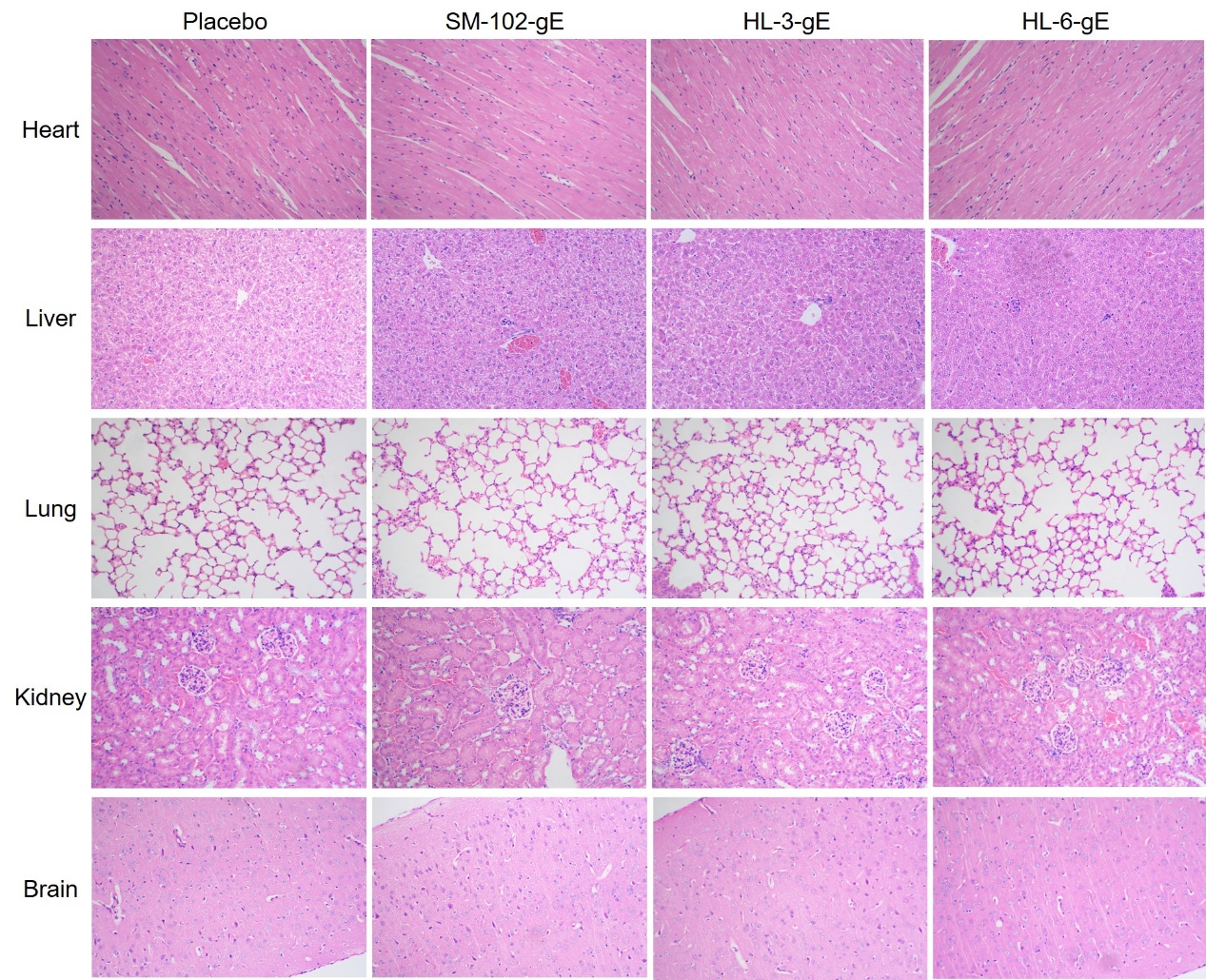

**Supplementary Fig. 38 The images of H&E-stained histopathological tissues from immunized mice.**
